# *linc-mipep* and *linc-wrb* encode micropeptides that regulate chromatin accessibility in vertebrate-specific neural cells

**DOI:** 10.1101/2022.07.21.501032

**Authors:** Valerie A. Tornini, Ho-Joon Lee, Liyun Miao, Yin Tang, Sarah E. Dube, Timothy Gerson, Valeria J. Schmidt, Katherine Du, Manik Kuchroo, François Kroll, Charles E. Vejnar, Ariel A. Bazzini, Smita Krishnaswamy, Jason Rihel, Antonio J. Giraldez

**Affiliations:** Department of Genetics, Yale University School of Medicine, New Haven, CT 06510, USA; Stowers Institute for Medical Research, Kansas City, MO 64110, USA; Department of Molecular & Integrative Physiology, University of Kansas School of Medicine, Kansas City, KS 66160, USA; Department of Computer Science, Yale University, New Haven, CT 06510, USA; Department of Cell and Developmental Biology, University College London, London WC1E 6BT, UK; Yale Stem Cell Center, Yale University School of Medicine, New Haven, CT 06510, USA; Yale Cancer Center, Yale University School of Medicine, New Haven, CT 06510, USA

## Abstract

Thousands of long intergenic non-coding RNAs (lincRNAs) are transcribed throughout the vertebrate genome. A subset of lincRNAs enriched in developing brains has recently been found to contain cryptic open reading frames and are speculated to encode micropeptides. However, systematic identification and functional assessment of these transcripts have been hindered by technical challenges caused by their small size. Here we show that two putative lincRNAs (*linc-mipep* and *linc-wrb*) encode micropeptides with homology to the vertebrate-specific chromatin architectural protein, Hmgn1, and demonstrate that they are required for development of vertebrate-specific brain cell types. Specifically, we show that NMDA receptor-mediated pathways are dysregulated in zebrafish lacking these micropeptides and that their loss preferentially alters the gene regulatory networks that establish cerebellar cells and oligodendrocytes – evolutionarily newer cell types that develop postnatally in humans. These findings highlight the power of screening for unexplored micropeptide functions by revealing a key missing link in the evolution of vertebrate brain cell development and illustrating a genetic basis for how some neural cell types are more susceptible to chromatin disruptions, with implications for neurodevelopmental disorders and disease.

## Introduction

While most of the vertebrate genome is transcribed, only a small portion encodes for functional proteins. Much of the remaining transcriptome is comprised of non-coding RNAs, including thousands of predicted long intergenic non-coding RNAs (lincRNAs). While thousands of lincRNAs have been identified, the functional significance of most remains unclear ^1^. Recent advances in ribosome profiling and mass spectrometry have identified short open reading frames (sORFs) within putative lincRNA sequences that may encode micropeptides, which were otherwise missed due to their small size (<100 aa)^2–7^. Despite conventional rules assuming that short peptides are unlikely to fold into stable structures to perform functions, and arbitrary cut offs (100 aa) used in computational identification of protein coding genes, there are several examples of these small peptides performing diverse, important cellular functions ^3,8–10^.

Remarkably, many lincRNAs are expressed in a tissue-specific manner, and about 40% of all lncRNAs identified in the human genome are specifically expressed in the central nervous system and brain ^11,12^. The vertebrate central nervous system consists of some of the most diverse and specialized cell types in the vertebrate body, and has distinct chromatin states and gene regulatory networks that have evolved to establish and maintain this diversity. Given their relatively small size, these proteins may be able to access and regulate cellular machines inaccessible to larger proteins ^13^.

Newly arising micropeptides may contribute to vertebrate-specific functions and phenotypes that have otherwise been missed due to misclassification as non-coding transcripts and lack of high-throughput phenotyping for coding functions. To tackle this question, we sought to identify micropeptides that were cryptically encoded in long non-coding RNAs but were missed due to assumptions about minimal protein sizes, dubious homologies, or mis-annotations. Here, we interrogate the function of predicted non-coding RNAs and identify two related micropeptides that regulate behavior, chromatin accessibility, and gene regulatory networks that establish evolutionarily newer neural cell types.

## Results

### Screen of long non-coding RNAs identifies micropeptide regulators of vertebrate behavior

To identify lincRNAs that may encode for micropeptides, we first analyzed ribosome profiles for previously published lincRNAs ^12^ in zebrafish embryos during early development (0-24 hours post-fertilization) (Bazzini et al., 2014) and performed *in situ* hybridization to identify brain-enriched micropeptide candidates (Extended Data Fig. 1a; Supplementary Table 1). To identify the physiological role of ten of these putative micropeptides, we used an F0 CRISPR/Cas9 behavioral screening pipeline at 4-7 days post-fertilization (dpf), during which zebrafish display a repertoire of conserved, stereotyped baseline locomotor behaviors across day:night cycles (Fig. 1a,b; Extended Data Fig. 1b, d) ^14–16 17^. This screen identified two candidate genes, *linc-mipep and linc-wrb*, that had daytime hyperactivity and correlated behavioral fingerprints when mutated in the ribosome protected ORF (Fig. 1c; Extended Data Fig. 1c). Sequence analysis revealed that *linc-mipep* (current nomenclature *si:ch73-1a9.3*, ENSDART00000158245, *lnc-rps25)*, and *linc-wrb* (current nomenclature *si:ch73-281n10.2*, ENSDART00000155252) ^2,12^ (Extended Data Fig. 2a-c; Supplementary Table 1), both had a significant homology in their sORFs and an ultra-conserved element in their non-coding sequences (Supplementary Note 1). While both *linc-mipep* and *linc-wrb* were originally identified as long non-coding RNAs, both genes have ribosome-protected fragments, suggesting they are likely translated (Fig. 1d,e). These results indicate that *linc-mipep and linc-wrb* might encode redundant or paralogous genes functioning as either lincRNAs or micropeptide-encoding genes involved in behavior.

**Fig. 1:**
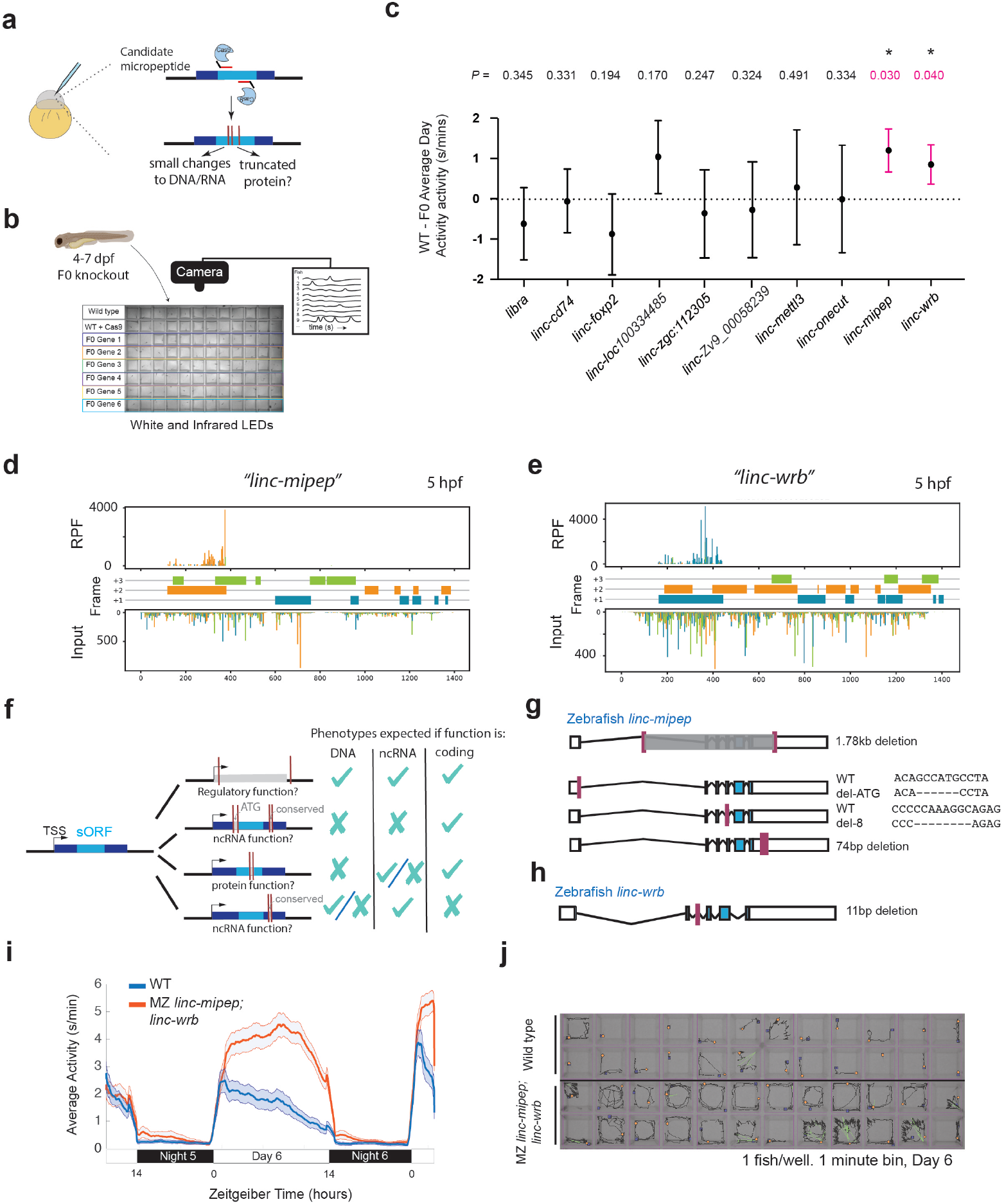
*linc-mipep* and *linc-wrb* loss-of-protein-function mutant larvae are behaviorally hyperactive. **a**. Schematic of F0 CRISPR knockout screen. Zebrafish embryos are injected at the 1-cell embryo stage with multiple sgRNAs targeting the ORF of candidate micropeptides encoded within putative lincRNAs. **b**. Schematic of behavior screening platform. A 96-well flat bottom plate contains individual larval (4-7 days post-fertilization, dpf) zebrafish per well from the same wild type (WT) clutch. Individual locomotor activity is tracked at 25 frames per second on a 14:10 light:dark cycle. **c**. Plots of the difference in average daytime activity at 6 dpf between WT and F0 knockouts of candidate micropeptides. Dotted line, wild type mean. P values (unpaired two-tailed t-test) above each targeted gene. **d**. Ribosome footprint of *linc-mipep (*also known as *lnc-rps25*) at 5 hours per fertilization (hpf) across annotated transcript length, with putative coding frames in green (+3), orange (+2), or blue (+1); input on bottom tracks. **e**. Ribosome footprint of *linc-wrb* at 5 hpf, same as d. **f**. Summary of mutagenesis strategy to decode transcript functions. Magenta bars denote CRISPR-targeted area. Mutated/removed sequence is in gray. TSS, transcription start site. sORF, short open reading frame. ATG, start codon. ncRNA, non-coding RNA. Right, phenotypes for each mutant predicted (check mark) or not predicted (x mark) if the gene functions as a regulatory region, noncoding RNA, or protein-coding gene. **g**. Stable mutants for *linc-mipep:* full region deletion (1.78kb deletion, from intron 1 – proximal 3’UTR, top); translation start site deletion that removes the ATG sequence (middle); frameshift deletion (8bp deletion at exon 4, bottom). **h**. Stable frameshift mutant for *linc-wrb* (11bp deletion, exon 3). **i**. Locomotor activity of wild type (WT, blue) or maternal-zygotic *linc-mipepdel1.8kb/del1.8bk;linc-wrbdel11bp/del11bp (linc-mipep;linc-wrb*, orange) larvae over 48 hours. The ribbon represents +/-SEM. Zeitgeber time is defined from lights ON=0. **j**. Representative daytime locomotor activity tracking of wild type (top 2 rows) and maternal-zygotic *linc-*mipepdel1.*8kb/del1.8kb;linc-wrbdel11bp/del11bp (linc-mipep;linc-wrb*, bottom 2 rows) 6 dpf fish over a 1 minute bin. Blue and orange dots represent start and stop locations, respectively.

### *linc-mipep;linc-wrb* encode for a micropeptide that regulates zebrafish behavior

To distinguish whether *linc-mipep and linc-wrb* function as regulatory DNA, noncoding RNA, or protein coding genes, we used CRISPR-Cas9 gene editing to generate stable deletion mutants that either target the full sequence, the translation start site, the putative coding region, or the conserved untranslated/non-coding region (Fig. 1f-h). Examining the behavioral profile of these mutants identified a consistent and specific increase in locomotor activity during the daytime in all mutants affecting the ORF for both *linc-mipep and linc-wrb* (Extended Data Fig. 2d-g). In contrast, deleting the ultra-conserved element in the untranslated region in *linc-mipep*, which could encode a conserved lincRNA sequence, did not result in any detectable morphological or behavioral phenotypes (Extended Data Fig. 2a,h). Is the coding part of these genes necessary? Start codon mutations in *linc-mipep* (zygotic or maternal-zygotic *linc-mipep*^*ATG-del6*^) resulted in similar daytime hyperactivity phenotype as frameshift mutations (*linc-mipep*^*del8*^) or deletion of most of the *linc-mipep* region (*linc-mipep*^*del1.78kb*^) (Extended Data Fig. 2d-f, i). These results indicate that the observed phenotypes are the result of protein coding function of *linc-mipep* rather than a non-coding transcript or a regulatory DNA sequence function. Double *linc-mipep*^*del1.78kb*^; *linc-wrb* ^*del11*^ homozygous mutants display even higher daytime locomotor hyperactivity levels compared to *linc-mipep; linc-wrb* heterozygous or wild type fish (p = 0.0036, one-way ANOVA), with no significant changes in nighttime activity (Extended Data Fig. 2j), a phenotype that is maintained if we remove the maternal contribution in maternal-zygotic (MZ) *linc-mipep*^*del1.78kb*^; *linc-wrb* ^*del11*^ animals (Fig. 1i,j). Next, is the coding part of these genes sufficient to drive behavior? To determine that the behavioral phenotypes observed in mutants result from the loss-of-coding function, we generated transgenic zebrafish that ubiquitously express the coding sequence (CDS) of *linc-mipep* (Fig. 2a). The sORF encoded in *linc-mipep* was able to rescue the hyperactivity phenotypes in *linc-mipep* heterozygous and homozygous mutants (Fig. 2b-d) without significant changes in nighttime activity (magnified, Fig. 2c).

**Fig. 2:**
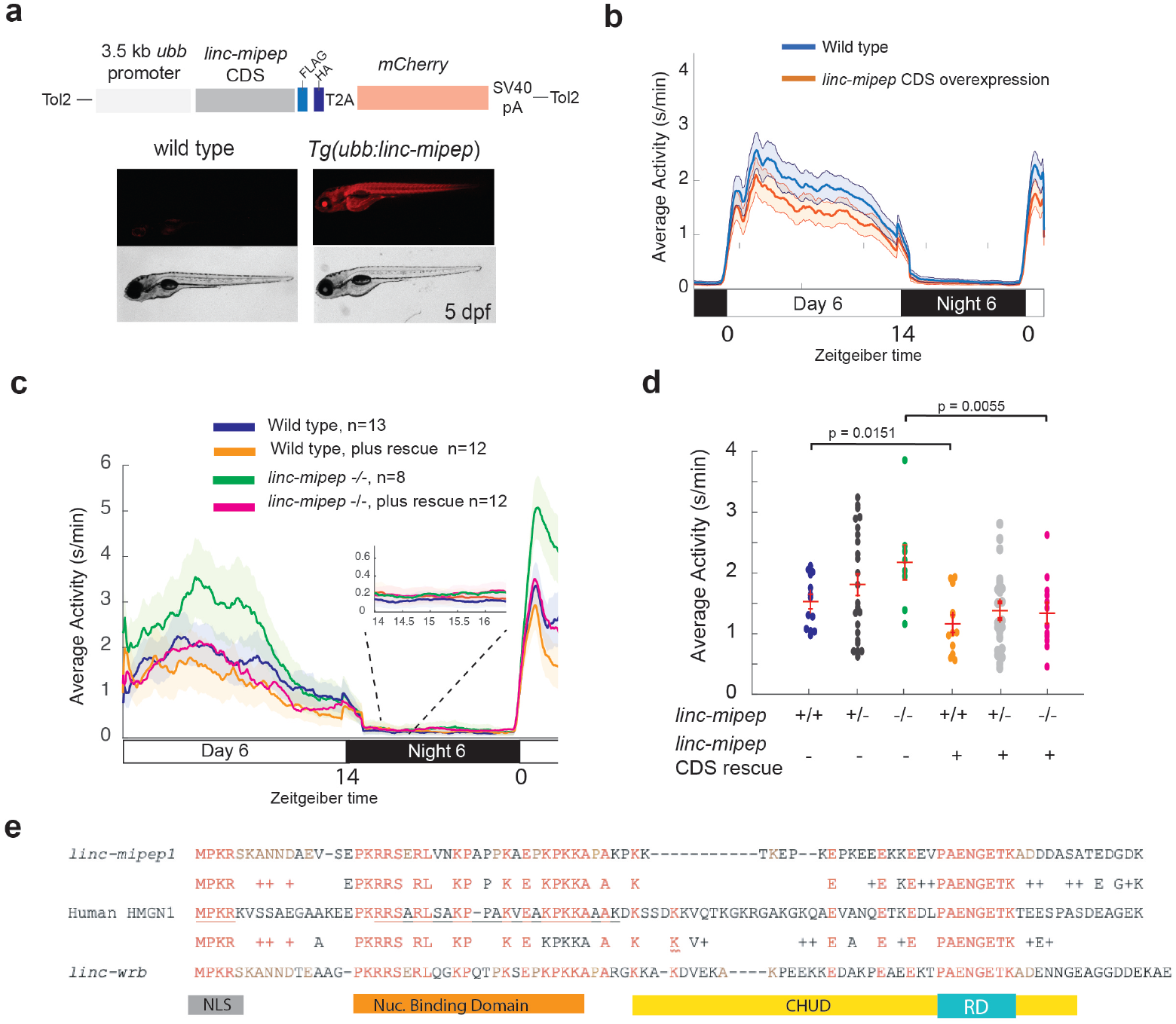
*linc-mipep* and *linc-wrb* encode proteins with homology to human HMGN1. **a**. Top, diagram of transgenic *linc-mipep* overexpression construct. Transgenic lines were established via Tol2-mediated integration of 3.5kb *ubiquitin B (ubb)* promotor driving the *linc-mipep* coding sequence with a FLAG and HA tag at the C-terminus, followed by a T2A self-cleaving peptide, *mCherry* reporter, and SV40 polyA tail. Bottom, fluorescent and brightfield images of 5 dpf zebrafish siblings, without overexpression (wild type, *mCherry-*negative, left) or with *linc-mipep* overexpression (*mCherry-*positive, right). **b**. Activity plot of wild type (mCherry-negative, blue) or *linc-mipep* overexpression (Tg(ubb:linc-mipep) mCherry-positive, orange) siblings at 6 dpf. n=48 per genotype. **c**. Locomotor activity of *linc-mipep* mutants, with or without transgenic *linc-mipep* overexpression (*Tg(ubb:linc-mipep CDS-T2A-mCherry)*, “rescue”), sibling-matched larvae over 24 hours. Inset, no effect on nighttime activity. **d**. Average waking activity of 6 dpf *linc-mipep* mutant, heterozygous, or wild type larvae, with (denoted with +) or without *linc-mipep* transgenic rescue (-)-,. Each dot represents a single fish, and crossbars plot the mean ± SEM. P values from a one-way ANOVA per genotype, Tukey’s post hoc test. **e**. Amino acid sequences of *linc-mipep* (top), *linc-wrb* (bottom), and human *Hmgn1* (middle). Conserved amino acids are denoted in brown (if conserved between 2 sequences) or red (if conserved across the 3 sequences). Conserved functional domains for Hmgn1 are denoted (NLS, nuclear localization signal; Nuclear Binding Domain; RD, Regulatory Domain; and CHUD, Chromatin Unwinding Domain).

Protein BLAST of both *linc-mipep* and *linc-wrb* ORFs identified conserved sequences across teleosts with homology to non-histone chromosomal protein HMG-14, or High Mobility Group N1 (HMGN1), and the related HMG-17/HMGN2 protein (Supplementary Table 3) ^18^. However, the ultra-conserved proximal 3’UTR elements allowed us to identify homologous predicted lincRNAs, unannotated genes, pseudogenes, and HMGN1 genes across vertebrate species spanning over 450 million years (Fig. 2e; Extended Data 3a-c, f; Supplementary Table 3)^19^. Though we did not identify any *linc-mipep* or *linc-wrb* protein-coding homolog in invertebrates (Supplementary Table 3), we did identify a syntenic ORF in the invertebrate lancelet (or amphioxus) genome, which may have been coopted to give rise to the HMGN gene and pseudogene families in vertebrates (Extended Data Fig. 3d). We further identified an unannotated ORF in the basal agnathan (jawless vertebrate) lamprey, syntenic to human HMGN1, that encodes for an ancestral protein more similar to human HMGN2 (Extended Data Fig 3e). Together, these findings suggest that *linc-mipep* and *linc-wrb* arose from the basal HMGN gene in agnathan lineages. Finally, to visualize the protein encoded by *linc-mipep* and *linc-wrb*, we developed custom antibodies. We observed that the protein product of both transcripts are enriched in non-dividing wild-type nuclei (Extended Data Fig. 2k,l) and absent in nuclei of *linc-mipep;linc-wrb* loss-of-function mutants (Extended Data Fig. 2m). Together, these results indicate that *linc-mipep and linc-wrb* encode for vertebrate-specific nuclear localized micropeptides with homology to non-histone chromosomal proteins (HMGN1) that have a dosage effect to regulate locomotor activity and behavior in zebrafish.

### *linc-mipep; linc-wrb* mutants have dysregulation of NMDA receptor-mediated signaling and immediate early gene induction

To gain insight into pathways regulated by *linc-mipep* and *linc-wrb*, we analyzed the behavioral fingerprints of each mutant compared to zebrafish larvae treated with 550 psychoactive drugs that affect different pathways. We used hierarchical clustering ^15^, to identify drugs that elicit a similar behavioral fingerprint to the mutant behavioral profile (i.e. drugs that phenocopy). Since both *linc-mipep* and *linc-wrb* have similar hyperactivity, we focused on *linc-mipep* to allow for drug analyses of mutant and wild type (WT) larvae with matched genetic backgrounds. We found that *linc-mipep* mutant behaviors most resembled those of WT fish treated with NMDA antagonists (Fig. 3a), suggesting that NMDA signaling may be reduced in *linc-mipep* mutants. The *linc-mipep* mutant phenotype also resembled that of WT fish treated with glucocorticoid receptor activators, representing 5/18 top correlating drug profiles (Fig. 3a, Extended Data Table 2).

**Fig. 3:**
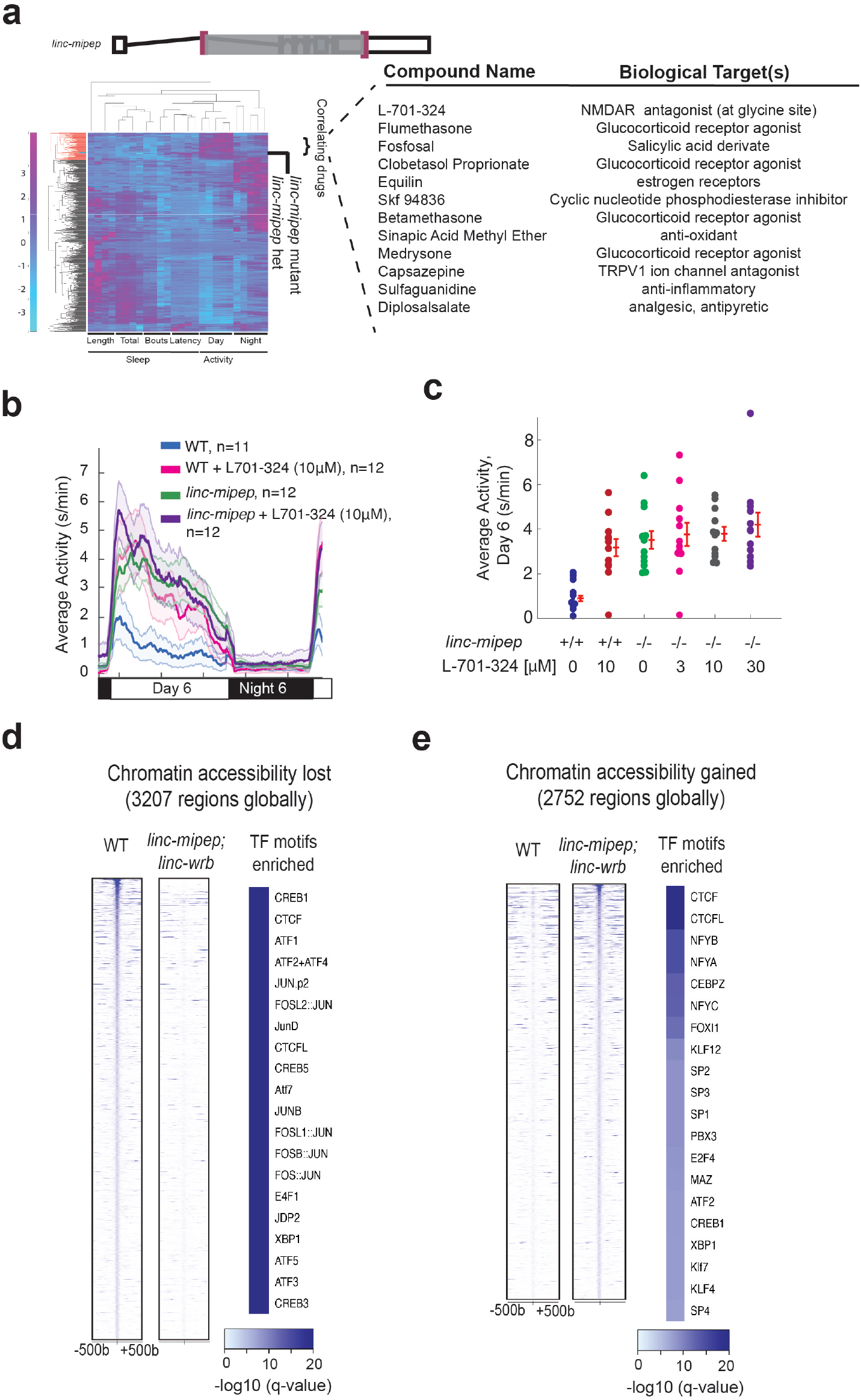
*linc-mipep* mutants have dysregulation of NMDA receptor-mediated signaling and immediate early gene induction. **a**. Left, hierarchical clustering of the *linc-mipep* ^*del-1.8kb*^ behavioral fingerprints (right), compared with the fingerprints of wild-type zebrafish larvae exposed to 550 psychoactive agents from 4–6 dpf (Rihel et al., 2010). The Z score, defined as the average value (in standard deviations) relative to the behavioral profiles of WT exposed to DMSO, is represented by each rectangle in the clustergram (magenta, higher than DMSO; cyan, lower than DMSO). The *linc-mipep* ^*del-1.8kb*^ fingerprint correlates with agents that induce daytime activity (‘‘Correlating Drugs’’). Right, compounds ranked according to correlation with the *linc-mipep*^*del-1.8kb*^ fingerprint, with biological target(s) noted in last column. **b**. Locomotor average activity of wild-type larvae treated with DMSO (WT, blue) or with 10mM NMDA receptor antagonist L-701,324 (magenta), and *linc-mipep* ^*del-1.8kb/del-1.8kb*^ larvae treated with DMSO (*linc-mipep*, green) or with 10mM L-701,324 (purple); sibling-matched larvae over 24 hours. **c**. Average activity (day 6) of WT larvae treated with DMSO or 10mM L-701-324, compared to *linc-mipep*^*del-1.8kb/del-1.8kb*^ larvae treated with DMSO or 3mM, 10mM, or 30mM L-701-324. Each dot represents one fish. **d**. Heatmaps showing chromatin accessibility (omni-ATAC-seq) profiles of 3207 regions globally with lower accessibility in *linc-mipep; linc-wrb* mutant brains at 5 dpfcompared to wild type (WT) brains. Heatmaps are centered at the summit of the Omni-ATAC peak with 500bp on both sides and ranked according to higher global accessibility levels in wild type. Transcription factor (TF) motifs enriched in the top 500 of these regions are presented on the right (-log10(q-value) scale at bottom). **e**. Heatmaps showing chromatin accessibility (omni-ATAC-seq) profiles of 2752 regions globally with higher accessibility in *linc-mipep; linc-wrb* mutant brains at 5 dpf compared to wild type (WT) brains. Heatmaps are centered at the summit of the Omni-ATAC peak with 500bp on both sides and ranked according to higher global accessibility levels in wild type. Transcription factor (TF) motifs enriched in the top 500 of these regions are presented on the right (-log10(q-value) scale at bottom).

The identified drugs may alter either common or parallel pathways in *linc-mipep* mutants. To distinguish between these possibilities, we first assessed glucocorticoid receptor agonists (flumethasone and clobetasol), which further exacerbated the daytime locomotor activity of *linc-mipep*^*-/-*^ larvae above the control-treated *linc-mipep* mutant levels (Extended Data Fig. 4a-d), suggesting that *linc-mipep* mutants are sensitized to glucocorticoid signaling. To test the NMDA pathway, we compared the response of WT and *linc-mipep* mutant to L-701-324, an NMDA antagonist at the glycine binding site. L-701-324 elicited a daytime locomotor hyperactivity in WT larvae to a level that was similar to that of *linc-mipep* mutant fish and *linc-mipep* treated with L-701-324 (Fig. 3b,c). Yet, higher doses of L-701-324 did not exacerbate the activity levels in *linc-mipep* mutants (Extended Data Fig. 4a). These non-additive results indicate that NMDA receptor antagonism and *linc-mipep* share a common mechanism for inducing hyperactivity.

### *linc-mipep* and *linc-wrb* regulate chromatin accessibility for transcription factors modifying neural activation

Given that *linc-mipep* and *linc-wrb* have protein domains with homology to nucleosome binding and chromatin unwinding domains of HMGN1 ^20,21^, and given that both NMDA antagonism and glucocorticoid signaling alters immediate early gene expression, we hypothesized that the daytime hyperactivity might be due to altered chromatin accessibility in the mutants. To test the effect of *linc-mipep;linc-wrb* on chromatin accessibility, we performed OMNI-ATAC at 5 dpf comparing WT and double mutant brains. We first observed a broad dysregulation of chromatin accessibility, with 3207 sites losing accessibility in *linc-mipep;linc-wrb* mutant brains, and 2752 regions gaining accessibility in *linc-mipep;linc-wrb* mutant brains compared to WT (Fig. 3d,e) (MACS2, P <10^−8^, presence, P>10^−3^ absence). CTCF/L transcription factor (TF) motifs were enriched in regions that lost as well as gained accessibility, indicating a broad dysregulation of 3D chromatin structure. Enriched TF motifs at sites that lost accessibility were members of the ATF (activating transcription factor)/CREB (cAMP responsive element binding proteins) family, and AP-1 transcription factor components. These TFs regulate the expression of immediate early response genes (IEG) such as *c-fos, c-jun, c-myc*, and *egr1* ^22^. We confirmed reduced *c-fos* transcription in *linc-mipep;linc-wrb* brains by *in situ* hybridization (Fig. 3f). We did not find significant accessibility changes in NMDA or glucocorticoid receptor signaling components, except for minimal loss in accessibility for only one NMDA receptor subunit, *grin2da* (Supplementary Table 4). On the other hand, TFs most enriched in sites that gained accessibility were NFY and KLF/SP family members, which promote stem cell pluripotency and are downregulated during differentiation, suggesting that *linc-mipep; linc-wrb* brain cells may be in a less differentiated state ^23^ (Fig. 3e). Altogether, these results indicate that *linc-mipep; linc-wrb* regulate chromatin accessibility for TF sites involved in neural activation and IEG expression.

### Evolutionarily newer vertebrate brain cell types are more susceptible to loss of *linc-mipep* and *linc-wrb*

Our molecular analyses of wildtype and mutant brains point to gene regulatory networks involved in global transcription rather than neural specific TFs. We hypothesize that the observed hyperactivity may instead be a result of cell-type specific defects most susceptible to loss of *linc-mipep* and *linc-wrb*. To test this hypothesis, we used single cell multiomics (transcriptomic and chromatin accessibility) and determined how single cell states are affected in *linc-mipep* mutant brains compared to sibling matched WT brains at 5 dpf (Fig. 4a). We used Weighted Nearest Neighbors (WNN) ^24^ and identified 43 clusters of cells, all comprised of both WT and *linc-mipep*^*-/-*^ cells (Fig. 4b, Extended Data Fig. 5a, b, Supplementary Table 2). *linc-mipep* was detected in all WT clusters except microglia, with enrichment in Purkinje cells, the inhibitory projection neurons of the cerebellum (Extended Data Fig. 5c-e).

**Fig. 4:**
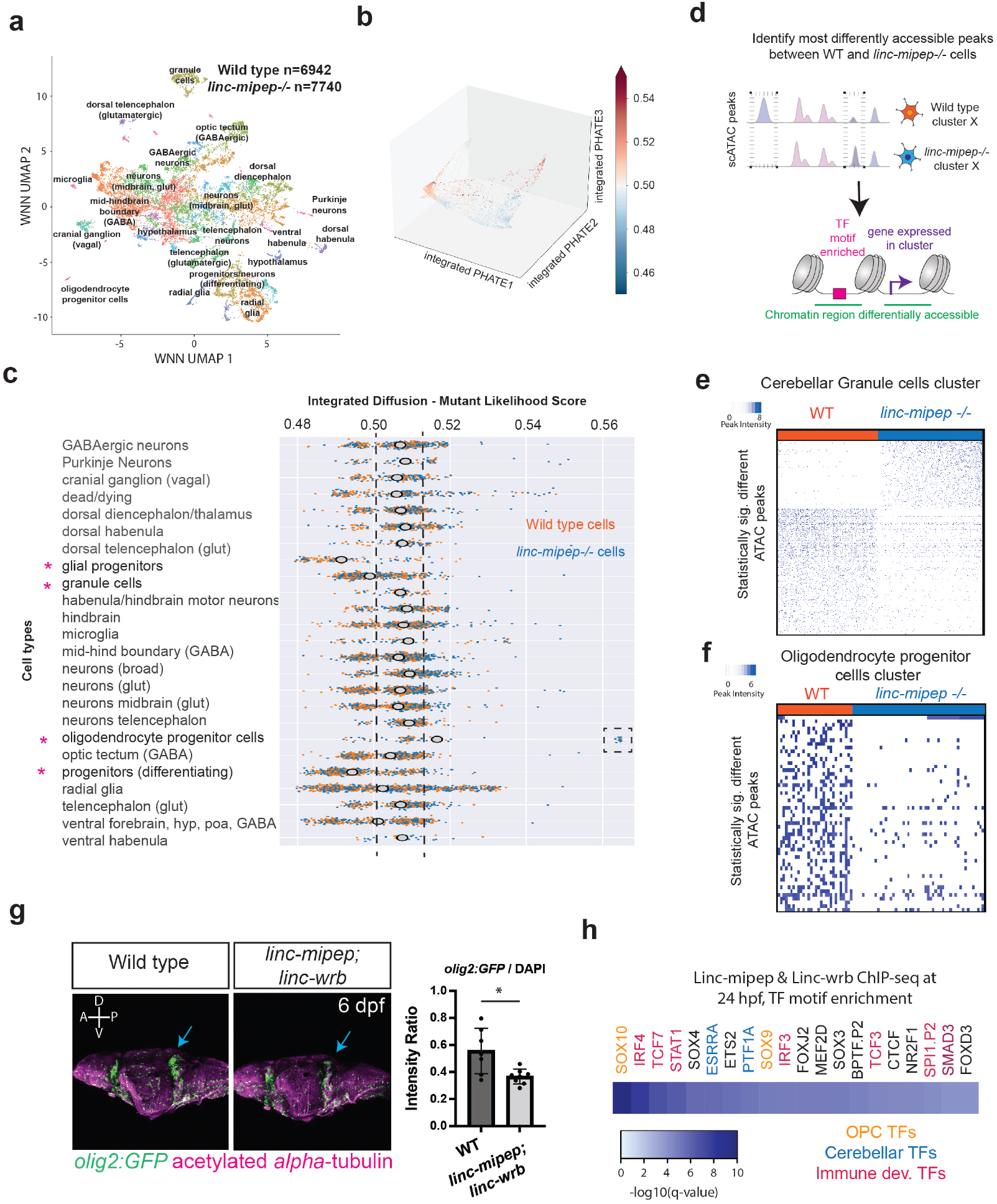
Evolutionarily newer vertebrate cell types are more susceptible to loss of *linc-mipep* and *linc-wrb* proteins. **a**. UMAP representation of WNN analyses of wild type (n=6942 nuclei) and *linc-mipep* ^*del-1.8kb del-1.8kb*^ (n=7740 nuclei) mutant brains at 6 days post-fertilization (dpf). Identified cell types as labeled. **b**. PHATE plot of integrated diffusion analysis of 6 dpf *linc-mipep* ^*del-1.8kb del-1.8kb*^ mutant or wild type (WT) sibling brain nuclei, color-coded by mutant likelihood score as computed by MELD using Integrated Diffusion operator. **c**. Integrated diffusion analysis on identified cell types from 6 dpf wild type (orange) and *linc-mipep* ^*del-1.8kb del-1.8kb*^ (blue) brains. Each dot represents a single cell, with mutant likelihood score across X-axis. Most wild type- or mutant-like groups noted with an asterisk. **d**. Schematic of analysis to identify most differentially accessible peaks between WT and *linc-mipep* ^*del-1.8kb del-1.8kb*^mutant brain nuclei from merged Weighted Nearest Neighbors (WNN) clusters. The most statistically significant changes in chromatin accessibility peaks were identified by the Wilcoxon rank sum and the Kolmogorov-Smirnov (KS) one-tailed tests methods on intensity distributions of each peak in WT and mutant samples, for either wild type or mutant differentially expressed genes per cluster, and for transcription factor (TF) motif overrepresentation by genotype in each cluster. **e**. Statistically significantly different chromatin accessibility peaks between 6 dpf wild type (WT, blue) and *linc-mipep* ^*del-1.8kbdel-1.8kb*^ mutant (red) nuclei in the cerebellar granule cells cluster. Each column is one nucleus. Color scale, peak intensity (blue, more accessible). **f**. Statistically significantly different chromatin accessibility peaks between 6 dpf wild type (WT, blue) and *linc-mipep* ^*del-1.8kb del-1.8kb*^ mutant (red) nuclei in the oligodendrocyte progenitor cells (OPCs) cluster. Color scale, peak intensity (blue, more accessible). **g**. Lateral view confocal images (Z-stack) from *Tg(olig2:GFP)* brains in wild type (left) or *linc-mipep;linc-wrb* double heterozygous mutant (right) backgrounds, stained with GFP (*olig2+*, green) and acetylated alpha-tubulin (magenta). A, anterior; P, posterior; D, dorsal; V, ventral. Quantification of intensity ratio of GFP+/DAPI signal of whole brain, two-tail t-test. * p<0.01. **h**. Enrichment of known sequence motifs of transcription factors (TFs) at gene promoters (+/-1000bp, n=315) from ChIP-sequencing of 24 hpf embryos using custom Linc-mipep and Linc-wrb antibodies. Transcription factors color-coded by known functions in OPCs (orange), cerebellar (blue), and immune (magenta) development.

To identify the most significantly affected cell states between WT and *linc-mipep* brain cells, we used Multiscale PHATE/Integrated Diffusion (Fig 4c,d asterisks; Extended Data Fig. 6a,b)^25,26^. This approach measures the effect of *linc-mipep* on cellular states by calculating the relative likelihood that a cell state would be observed in either WT or mutant. We found that differentiating neuronal progenitor, glial progenitor, and cerebellar granule (excitatory) cell states were more likely observed in WT brain cells, while oligodendrocyte progenitor cell (OPC) states more likely associated with *linc-mipep*^*-/-*^ brain cells (Fig. 4c, asterisks; Extended Data Fig. 6 a-e; Supplementary Table 6). Indeed, we find a subcluster of oligodendrocyte progenitor cells strongly enriched in *linc-mipep*^*-/-*^ samples (Fig. 4c, dashed box). These data indicate that *linc-mipep* and *linc-wrb* preferentially regulate oligodendrocyte and cerebellar cell states during development – evolutionarily newer vertebrate brain cell types ^27^.

We next searched for the most significant changes in chromatin accessibility and differentially expressed genes in each identified cluster between WT and mutant brains (Fig. 4d, Extended Data Fig. 6d-f; Supplementary Table 5). While all cell types are maintained in both conditions, we found a strong misregulation of chromatin accessibility and gene expression within multiple cell types (Supplementary Table 5; Supplementary Data Fig. 1). Chromatin accessibility was strongly conditioned by the presence of *linc-mipep* across clusters. TF motifs in regions of differential chromatin accessibility per cluster indicate that *linc-mipep* regulates accessibility for key neurodevelopmental transcription factor families, including Sox, Stat, and Zic family members in radial glial cells (clusters 3 and 7), among others; Esrra/b in midbrain glutamatergic neurons (cluster 13); and Egr and NeuroD members across various cell states (Extended Data Fig. 7a).

In cerebellar granule cells, we found 989 regions where chromatin accessibility was strongly dependent on *linc-mipep* function (645 regions with decreased accessibility, and 344 regions with increased accessibility) (Fig. 4e). Purkinje neurons also showed significant differences in chromatin accessibility (Extended Data Fig. 6f). We specifically assessed *linc-mipep* granule cells and found that they lost accessibility to Bhlhe22 (an inhibitor of Neurod1-responsive genes), Hic1 (expressed in mature granule cells), Neurod2 (required for survival of granule cells) ^28^ and Nfia TF binding motifs; and gained accessibility at sites for Gfi1b and Nfatc1, both of which induce and respond to cytokines (Extended Data Fig. 7b). Consistent with these results, Pol II ChIP-seq in 5dpf brains found that transcription factors and genes involved in cerebellar development, including *zic2a, ascl1b, foxo6b, adnpa, atx3*, and *rora/b*, showed reduced RNA Polymerase II binding in mutant brains (Supplementary Table 7).

OPCs showed a broad loss of chromatin accessibility in the absence of *linc-mipep* (Fig. 4f). *Linc-mipep* OPCs lost accessibility to E2f7, Elf1 (induced in differentiating oligodendrocytes), Fev, and Hinfp TF sites, while they gained accessibility at Sox10 sites (Extended Data Fig. 7c). Consistent with these results, we found a significant reduction (P=0.0048) in *olig2+* oligodendrocyte progenitor cells across the brain and a more significant reduction in the midbrain and the cerebellum of *Tg(olig2:eGFP); linc-mipep*^*+/-*^; *linc-wrb*^*+/-*^ compared to control larvae (Fig. 4g, arrows).

Finally, ChIP-seq analysis for *linc-mipep* and *linc-wrb* during embryonic development (24 hpf), when different progenitors are being specified, revealed that bound sites were enriched for TF motifs reminiscent of those described in Extended Data Fig. 7a, including for critical regulators of oligodendrocytes (Sox10 and Sox9, specifically required in mouse cerebellar oligodendrocytes)^29–31^ and cerebellar development (Esrra/Nr3b11 and Ptf1a) ^32,33^ (Supplementary Table 8). Together, these data indicate that *linc-mipep* and *linc-wrb* proteins broadly regulate the chromatin state of neural cell types, most impacting OPCs and cerebellar granule cell gene expression networks and cell states in a basal vertebrate.

## Discussion

Here, we establish that *linc-mipep* (or *lnc-rps25*) and *linc-wrb*, previously identified as long non-coding RNAs, are in fact micropeptides with homology to the vertebrate-specific non-histone chromosomal protein HMGN1. These genes regulate oligodendrocyte and cerebellar development and behavior in a dose-dependent manner. Our datasets allow for deep probing of cell type-specific effects on loss of these genes in a tractable basal vertebrate system.

LincRNAs represent a prevalent and functionally diverse class of non-coding transcripts that likely emerged anew from previously untranscribed DNA sequences, either by duplication from other ncRNAs or from transformation of coding regions^12^. While it is possible that *linc-mipep, linc-wrb*, and HMGN1 arose from an originally non-coding transcript, possibly in invertebrates, here we identify a basal-most vertebrate sequence in lamprey for an ancestral HMGN protein lacking the key C-terminal regulatory domain of human HMGN1. This ancestral protein is likely derived from an unannotated ORF in the invertebrate *Amphioxus* (lancelet) encoding for an APEX1-like gene in the HMGN1 syntenic region. Neural crest cells, myelinating cells (both oligodendrocytes in the CNS and neural crest-derived Schwann cells in the peripheral nervous system), and cerebellar cells (including granule cells) are considered to be among these jawed vertebrate-specific innovations ^27,34^. We hypothesize that *linc-mipep, linc-wrb*, and HMGN1 co-evolved with the gene regulatory networks that establish these cell types in development, in line with previous findings ^35–40^, as we find that these evolutionarily newer brain cell types are most affected by loss of *linc-mipep* and *linc-wrb* in zebrafish. We find that mutations in *linc-mipep* and *linc-wrb* don’t change cell fate. Rather, they alter the chromatin state, and therefore cell state, with misregulation of chromatin accessibility across hundreds of sites in the genome and across multiple transcription factors during neural development, with oligodendrocytes and cerebellar cells being most susceptible to these changes.

Both oligodendrocyte progenitor cells and cerebellar cells are typically associated with post-natal growth in humans. The cerebellum is a folded hindbrain structure important for coordinating body movements and higher-order cognitive functions. Our results are consistent with recent findings in Trisomy 21 (Down syndrome) pathology, in which HMGN1 is overexpressed, that developing and adult Down syndrome brains have dysregulated expression of genes associated with oligodendrocyte development and myelination in addition to alterations in the cerebellar cortex ^41–43^. Oligodendrocytes are thus also emerging as an important cell type required for normal neurodevelopment and dysregulated in neurodevelopmental disorders ^44^. Our behavioral mutant analyses highlight the dose-dependent roles of *linc-mipep* and *linc-wrb;* evolutionarily conserved functions between *linc-mipep, linc-wrb*, and human HMGN1 in neurodevelopment ^21,35,45^; and the importance of understanding the ancestral and conserved roles of key neurodevelopmental genes in basal vertebrate systems. Altogether, these studies emphasize the importance of non-neuronal and non-cerebral cortex cell types in neurodevelopmental disorders,^46^ in which the vertebrate-specific *Hmgn1* and related proteins may play a unifying role by regulating chromatin accessibility for key transcription factors.

Overall, this study highlights the power of using a high-throughput, genetically tractable vertebrate model to systematically screen for micropeptide function within putative lincRNAs, behavioral phenotypes, signaling pathways, and cell type susceptibilities in early vertebrate development. How novel protein-coding genes may be born from non-coding genomic elements remains an elusive question ^47^. Several short open reading frames encoding for functional, evolutionarily conserved peptides now have been discovered within putative non-coding RNAs ^13^, and some of these genes may have emerged along vertebrate lineages (for example, *libra/*NREP ^48^). Our analyses support that many more unannotated or undescribed proteins may similarly play critical roles in vertebrate neurodevelopment and behavior ^49^. We propose that revisiting sORFs identified within putative long non-coding RNAs in basal vertebrates may provide insight into gene innovation and evolution. This framework will enable genetic studies in a basal system to understand the evolutionary origins of human developmental disorders and diseases in a vertebrate cell type-specific manner.

## Data Availability

The datasets generated and analyzed in this study will be made available through the Gene Expression Omnibus (GEO) database. The datasets, plasmids, antibodies, and fish lines generated during and/or analysed during the current study are available from the corresponding authors on request.

## Code Availability

Code generated and used in this study is available through GitHub (or equivalent) repositories. Links with code are provided in the methods section.

## Acknowledgements

We thank Dr. Shawna Hiley for editorial and scientific input; Dr. Kaya Bilguvar, Christopher Castaldi, and Dr. Guilin Wang from the Yale Center for Genome Analysis for sequencing support; Dr. Kaelyn Sumigray for sharing Leica confocal; Dr. Mayssa Mokalled for sharing animal transgenic lines; Dr.

Marcus Ghosh for code used in the F0 behavioural data analysis and for teaching F.K. the approach; and Dr. Sumru Bayin, Dr. Sarah Ackerman, and members of the Giraldez and Rihel labs for critical feedback. This work is supported by a fellowship from the Hartwell Foundation (V.A.T.), Wellcome Trust Investigator Award 217150/Z/19/Z (J.R.), Simons Foundation grant and NIH grants R01 HD100035 and MH118554 (A.J.G). We acknowledge the Zebrafish Information Network (ZFIN).^62^

## Materials and Methods

### Zebrafish Husbandry and Care

Fish lines were maintained in accordance with the AAALAC research guidelines, under a protocol approved by the Yale University Institutional Animal Care and Use Committee. We have complied with all relevant ethical regulations under this protocol. Zebrafish husbandry and manipulation were performed as described, and all experiments were carried out at 28°C. For all larval experiments, zebrafish embryos were raised at 28.5 °C in petri dishes at densities of 70 embryos/dish on a 14hr:10hr light:dark cycle in a DigiTherm® 38-liter Heating/Cooling Incubator with circadian lighting (Tritech Research). Dishes of embryos were cleaned once per day with blue water (fish system water with 1mg/L methylene blue, pH 7.0) until they were placed in behavior boxes, to ensure identical growing conditions. Normal development was assessed, and larvae exhibiting abnormal developmental features (no inflated swim bladder, curved) were not used.

### Ribo-seq profiles

Sequences and code for updated ribosome profiling plots available at https://www.giraldezlab.org/data/ribosome_profiling/ and https://github.com/vejnar/notebooks/blob/main/ribosome_profiling/ribo_orf_plot.ipynb.

### CRISPR F0 and mutant generation

CRISPR mutant generation was done following Vejnar, Moreno-Mateos, et al. (2016). Briefly, CRISPRScan (crisprscan.org) was queried to identify appropriate target sequences ^50^. Primers were ordered and amplified with universal primer 5’-AAAAGCACCGACTCGGTGCCACTTTTTCAAGTTGATAACGGACTAGCCTTATTTTAACTTGCTA TTTCTAGCTCTAAAAC-3’. sgRNAs were *in vitro* transcribed using the AmpliScribe T7 Flash kit, using the PCR product (with T7 promoter) as template. *In vitro* transcribed sgRNAs were treated with DNase I and precipitated with sodium acetate and ethanol. For F0 CRISPR experiments and low-efficiency target sequences, synthetic guides were designed using CRISPRscan and ordered through Synthego (Synthego Corportation, Redwood City, CA, USA). *Cas9* mRNA was *in vitro* transcribed from DNA linearized by XbaI (pT3TS-nCas9n) using the mMESSAGE mMACHINE T3 kit (Ambion). *In vitro* transcribed mRNAs were treated with DNase I and purified using the RNeasy Mini kit (Qiagen). EnGen Spy Cas9 NLS protein (NEB, M0646) was used for F0 experiments.

One-cell stage zebrafish embryos were injected with 50 pg of each respective sgRNA and 100 pg of cas9 mRNA. sgRNA and genotyping primers and target sequences are available in Supplementary Table 1.

### Quantitative locomotor activity tracking and statistics for sleep/wake analyses

On 4 dpf, single larvae from heterozygous *linc-mipep* mutant incrosses were placed into individual wells of a clear 96-square well flat plate (Whatman) filled with 650 μL of blue water (fish system water with 1mg/L methylene blue, pH 7.0). Plates were placed in a Zebrabox (ViewPoint Life Sciences), and each well was tracked using ZebraLab (Viewpoint) in quantized mode, and analyzed with custom software as in Rihel et al., 2010. Behavioral data were analyzed for statistical significance using one-way ANOVA followed by Tukey’s post hoc test (α = 0.05), as previously described^15^. For analyses of maternal-zygotic *linc-mipep;linc-wrb* mutants, age- and size-matched wild type adult stocks (AB/TL) or *linc-mipep;linc-wrb* double-homozygous mutants were incrossed, collected simultaneously, and raised in identical conditions prior to quantitative locomotor activity tracking as described above.

### Behavioural fingerprints and Euclidean distances

As previously described ^17^, the raw file generated by the ZebraLab software (ViewPoint Life Sciences) was exported into a series of xls files each containing 1 million rows of data. Each datapoint represented the number of pixels that changed grey value above a sensitivity threshold, set to 18, for one larva at one frame transition, a metric termed Δ pixels. These files, together with a metadata file labelling each well with a genotype, were input to the MATLAB script Vp_Extract.m ^51^, which calculated the following behavioral parameters from the Δ pixels timeseries for both day and night: (1) active bout length (duration of each active bout in seconds); (2) active bout mean (mean of the Δ pixels composing each active bout); (3) active bout standard deviation (mean of the Δ pixels composing each active bout); (4) active bout total (sum of the Δ pixels composing each active bout); (5) active bout minimum (smallest Δ pixels of each bout); (6) active bout maximum (largest Δ pixels of each bout); (7) number of active bouts during the entire day or night; (8) total time active (% of the day or night); (9) inactive bout length (duration of each pause between active bouts in seconds). These measurements were then averaged across both days or both nights to obtain one measure per parameter per larva for the day and night. To build the behavioral fingerprints, we calculated the deviation (Z-score) of each mutant (F0) larva from the mean of their wild-type siblings across all parameters. Plotted in Extended Data Fig. 1b,c for each parameter is the mean ± SEM of the Z-scores. We compared fingerprints between replicates (Extended Data Fig. 1b) or between *linc-mipep* and *linc-wrb* (Extended Data Fig. 1c) using Pearson correlation. The behavioral fingerprint of each larva can be conceptualized as a single datapoint in a multidimensional space where each dimension represents one behavioral parameter. To summarize the intensity of each phenotype across parameters, we measured the Euclidean distance between each larva and the mean fingerprint of its wild type siblings, set at the origin of this space by the Z-score normalization (Extended Data Fig. 1d). Code for this analysis is available on GitHub (https://github.com/francoiskroll/micropeptides_fingerprints). Statistics for Fig. 1c were calculated using Prism (GraphPad).

### Hierarchical clustering

Correlation analysis was done in MATLAB (R2018a; The MathWorks) as previously described ^15^ (Rihel et al., 2010). Behavioral phenotypes of wild-type fish exposed to a panel of 550 psychoactive agents from 4 - 7 dpf were ascertained as previously described (Rihel et al., 2010). To compare the behavioral fingerprints of WT larvae exposed to each drug and the *linc-mipep* mutant behavioral fingerprint, hierarchical clustering analysis was performed as in ^15,52^.

### Sequence alignments and homologies

BLASTp and BLASTn, and UCSC Genome Browser were used to find sequences (especially ultra-conserved 3’UTR sequence) and proteins with sequence homology and/or synteny to human *Hmgn1*. Clustal Omega (through EMBL-EBI) was for multiple sequence alignments.

### Custom antibodies generation

Three custom antibodies were designed (YenZym Antibodies, LLC) against: *Si:ch73-1a9.3 (linc-mipep)*, C-Ahx-DDASATEDGDKKEDGE-cooh; *Si:ch73-281n10.2 (linc-wrb)*, C-Ahx-EDAKPEAEEKTP-amide; and both *Si:ch73-1a9.3* and *Si:ch73-281n10.2*: KRSKANNDAE-Ahx-amide. The last antibody designed to recognize both proteins was non-specific and not further used. Antibody specificity was confirmed by antibody staining in wild type and *linc-mipep; linc-wrb* mutants.

### Antibody staining and imaging

Embryos up to 24 hpf: Embryos were dechorionated and collected into room-temperature 4% PFA in PBS for 1 hour. Embryos were blocked rotating for 1 hour at room temperature in 10% normal goat serum (NGS) (Thermo Fisher Scientific, 50062Z), primary antibody stained for 1 hour at room temperature in 10% NGS, washed 3x 5 minutes in 1xPBS with 0.25% Triton-X (PBST), incubated rotating and protected from light for 1 hour at room temperature, was 3×5 minutes in PBST, and mounted in 0.7-1% low-melt agarose on glass-bottom dishes (MatTek) for imaging. *Larvae*: Larvae (up to 6 dpf) were maintained in a quiet environment. For assessment of olig2+ cells, the *Tg(olig2:egfp)*^*vu12*^ line ^53^ was used in either wild type or *linc-mipep1;linc-wrb* double heterozygous mutant backgrounds. To ensure rapid fixation, larvae were poured through a mesh sieve and immediately submerged into ice-cold 4% PFA (Electron Microscopy Sciences) /1x PBS-0.25% Triton X-100 (PBST)/4% sucrose, in fix, as previously described ^54^. Larvae were fixed overnight at 4°C and washed 3 times for 15 minutes each in PBST. For whole larvae, pigment was bleached with a 1% H_2_O_2_/3% KOH solution (in PBS), washed 3x 15 minutes in PBST, then permeabilized with acetone (pre-cooled to −20°C) at −20°C for 20 minutes, and washed 3 times for 15 minutes with PBST. For dissecting brains, following overnight fixation, larvae were washed 3 × 5 minutes in PBST, then brains were dissected by hand and transferred back into tubes with PBS. Brains were sequentially dehydrated 5 minutes each in 25% MeOH/75% PBS, 50% MeOH/50% PBS, 75% MeOH/50% PBS, and 100% MeOH, and stored at −20°C for at least overnight. Brains were sequentially similarly rehydrated, then permeabilized with 1x Proteinase K (10mg/ml is 1000x stock) in PBST for exactly 10 minutes. Brains were then rinsed 3x with PBST, post-fixed in 4% PFA/PBST for 20 minutes at room temperature, and washed 3 times for 5 minutes in PBST. Samples were mounted in 0.7-1% low-melt agarose on glass-bottom dishes (MatTek) for imaging. Confocal imaging was performed using a Zeiss 980 AiryScan or a Leica SP8 confocal microscope. Images were processed and analyzed using FIJI software and plugins.

Primary antibodies used: custom Linc-mipep (rabbit); custom Linc-wrb (rabbit); anti-GFP (mouse, A11120, Thermo Fisher Scientific, 1:500); acetylated a-tubulin (rabbit, 5335T, Cell Signaling Technology, 1:500). Alexa Fluor 488, 546 or 568 secondary antibodies against rabbit or mouse were used at 1:500 (Invitrogen). DAPI (for nuclear marking) was added at 1:10,000 during secondary antibody staining.

### Overexpression constructs

A gBlock (IDT) was ordered for the *linc-mipep* coding sequence, plus a FLAG and HA tag at the C terminus (5’-gccaccATGCCTAAAAGGAGCAAAGCGAACAATGACGCT GAAGTCTCTGAGCCTAAAAGAAGGTCAGAGAGGTTGGTAAACAAACCTGCACCCCCAAAGG CAGAGCCCAAGCCAAAGAAGGCCCCTGCCAAACCTAAGAAAACAAAGGAACCCAAGGAGC CCAAGGAGGAGGAGAAGAAAGAGGAGGTGCCCGCAGAAAACGGAGAAACAAAAGCTGAC GATGATGCATCGGCAACAGAAGACGGCGACAAGAAAGAAGACGGGGAAGGTTCTGGCTCAg actacaaagacgatgacgacaagtacccatacgatgttccagattacgctTAA-3’). Addgene plasmid #79885 (pMT-ubb-cytoBirA-2a-mCherry, a gift from Tatjana Sauka-Spengler^55^) was digested with BamHI and EcoRV, and the resulting vector was used as the backbone for the construct. InFusion cloning (Takara Bio) was used to amplify the linc-mipep-FLAG-HA coding sequence and ligate with the vector, using primers F: 5’-TTGTTTACAGGGATCgccaccATGCCTAAAAGGAGC-3’ and R: 5’-CTCTCCTGATCCGATagcgtaatctggaacatcgtatggg. Sequence-verified plasmids were midi-prepped and injected into the cell of one-cell stage embryos at 20 ng/ml along with 200 ng/ul of Tol2 transposase capped mRNA.

### In vivo pharmacological drug experiments

At 4 dpf, single larvae from heterozygous *linc-mipep* mutant incrosses were placed into individual wells of a clear 96-square well flat plate (Whatman) filled with 650 μL of blue water. Respective pharmacological agents (from a stock of 5 or 50mM depending on solubility) or corresponding vehicle controls (DMSO or water) were pipetted directly into the water to achieve the desired final concentrations at the start of the experiment (typically evening of 4 dpf). Drug treatments, vehicles, and doses are described in Supplementary Table 3.

### Genotyping

After each behavioral tracking experiment, larvae were anesthetized with an overdose of MS-222 [0.2 to 0.3 mg/mL], transferred into 96-well PCR plates, and incubated in 50 μl of 100 mM NaOH at 95 °C for 20 min. Then, 25 μl of Tris-HCl 1 M pH 7.5 was added to neutralize the mix. 2 μl of these crude DNA extracts were used for genotyping with the corresponding forward and reverse primers (10 μM; Supplementary Table 1) using a standard PCR protocol.

### Brain collection for molecular analyses

Briefly, brains at peak daytime activity levels (Zeitgeber Time 4, i.e. 4 hours after lights on) were dissected from 5 dpf MZ-*linc-mipep;linc-wrb* or wild type zebrafish (for omni-ATAC-seq n=10 per sample, and ChIP-seq n=50 per sample) or 6 dpf zebrafish from one *linc-mipep-/-* heterozygous incross (for single-cell Multiome, n=12 per sample) in ice-cold Neurobasal media supplemented with B-27 (Thermo Fisher Scientific), snap-frozen in a dry ice/methylbutane bath (to preserve nuclear structure), and stored at −80°C until use. Trunks of *linc-mipep* fish were genotyped, then wild type or *linc-mipep-/-* brains pooled together before proceeding with scMulitome.

For ChIP-seq experiments, brains were dissected and homogenized before treatment with 1% PFA (protocol adapted from ^56^) and performed as previously described ^57^ using 4mg of RNA Polymerase II antibody (ab817, Abcam) per sample; 5% input samples were also collected and processed.

Omni-ATAC was performed on frozen brains from 5 dpf zebrafish based on published protocols ^58,59^. Frozen brain tissue was homogenized in cold homogenization buffer (320 mM sucrose, 0.1 mM EDTA, 0.1% NP40, 5 mM CaCl2, 3 mM Mg(Ac)2, 10 mM Tris pH 7.8, 1× protease inhibitors (Roche, cOmplete), and 167 μM β-mercaptoethanol) on ice. The lysate was filtered with a tip strainer (Flowmi® Cell Strainers, porosity 70 μm) into a new Lo-Bind tube. Nuclei were isolated using the gradient iodixanol solution as described^59^. Nuclei solution was mixed with 1 ml of dilution buffer (10 mM Tris-HCl pH 7.4, 10 mM NaCl, 3 mM MgCl2, 0.1% Tween-20) and was then centrifuged at 500 x g for 10 minutes at 4°C. Transposition and library preparation were performed on the purified nuclei as described ^57^.

The supernatant was removed, and the purified nuclei were resuspended in the transposition reaction mixture (25 μl 2×TD Buffer, 2.5 μl Tn5 transposase, 22.5 μl Nuclease-Free water) and incubated for 30 minutes at 37°C. DNA was then purified with the Qiagen MinElute Kit (Qiagen, 28004). Libraries were prepared using NEBNext High-Fidelity 2X PCR Master Mix (NEB, M0541) with the following conditions: 72°C, 5 minutes; 98°C, 30 seconds; 15 cycles of 98°C, 10 seconds; 63°C, 30 seconds; and 72°C, 1 minute. Libraries were purified with Agencourt AMPureXP beads (Beckman Coulter Genomics, A63881) and sequenced with the Illumina NovaSeq 6000 System at the Yale Center for Genome Analysis.

### High-throughput sequencing data management

LabxDB seq ^60^ was used to manage our high-throughput sequencing data and configure our analysis pipeline. Export to the Sequence Read Archive was achieved using the “export_sra.py” script from LabxDB Python.

### Omni-ATAC data processing and analysis

Raw paired-end Omni-ATAC reads were adapter trimmed using Skewer (Jiang et al., 2014) and mapped to the zebrafish GRCz11 genome sequence (Yates et al., 2019) using Bowtie2 (v2.3.4.1) (Langmead and Salzberg, 2012) with parameters ‘-X 2000, --no-unal’. Unpaired and discordant reads were discarded. The alignments were deduplicated using samtools markdup (Li et al., 2009). Reads mapped to the + strand were offset by +4 bp and reads mapped to the – strand were offset by −5 bp (Buenrostro *et al*., 2013). Only fragments with insert size <=100 bp (effective fragments) were used to determine accessible regions. Genome tracks were created using BEDTools (Quinlan and Hall, 2010) and utilities from the UCSC genome browser (Lee et al., 2020). For all the genome tracks in the paper, signal intensity was in RPM (reads per million). Fragment coverage on each nucleotide was normalized to the total number of effective fragments in each sample per million fragments. For genome-wide analysis, only uniquely mapped reads (with alignment quality ≥ 30) were used.

### ChIP-seq data processing and analysis

Raw ChIP-seq reads were adapter trimmed, mapped, and deduplicated using the same method described in the previous section but using the default parameters for Bowtie2 for read mapping. GeneAbacus (all code available at https://github.com/vejnar/geneabacus) was used to create genomic profiles for creating tracks. Fragment coverage on each nucleotide was normalized to the total fragments in each sample per million fragments. For genome-wide analysis, only uniquely mapped reads (with alignment quality ≥ 30) were used.

### Peak calling

Peaks were called using MACS2 (Zhang *et al*., 2008) for Omni-ATAC and ChIP-seq data. For Omni-ATAC, peaks were called with the additional parameters ‘-f BEDPE --nomodel --keep-dup all’ with significance cutoff at *P* = 10^−8^ (high threshold) and *P* = 10^−3^ (low threshold). For ChIP-seq of RNA Polymerase II, narrow peaks were called using MACS2 with the additional parameters ‘-f BEDPE --nomodel --keep-dup all’ with the default significance cut-off (q = 0.05, high threshold) and *P* = 0.05 (low threshold). Peaks that are called at high threshold in one condition but not called at low threshold in the other condition are defined to be specific to the condition. Genes with promoter regions (+/-1kb of transcription start site) that overlap with a peak are defined to be associated with peak.

### Motif enrichment analysis

Motif enrichment was performed using AME in MEME suite (McLeay and Bailey, 2010) with default parameters on all known transcription factor binding motifs from the Motif database on the MEME suite website (http://meme-suite.org/doc/download.html) and HOMER website (http://homer.ucsd.edu/homer/custom.motifs). Motifs for human and mouse transcription factors were used as the motifs for their homologous transcription factors in zebrafish. Homologs between zebrafish and human and mouse were identified using BioMart on Ensembl genome browser (Yates *et al*., 2019). The top 500 or 1000 most significant peaks (all peaks if sample size <500 or 1000) were used for motif enrichment across different conditions.

### Heatmaps and plots

Heatmaps based on Omni-ATAC and ChIP-seq data were created using R 4.1 and the pheatmap package (https://github.com/raivokolde/pheatmap). Motif enrichment and histone acetylation/transcription correlation heatmaps were created using the R package gplots (https://github.com/talgalili/gplots).

### Data analysis of scRNA-seq and scATAC-seq

The raw 10x Genomics Multiome data of scRNA-seq and scATAC-seq were processed using the 10x Genomics cellranger-arc pipeline (v1.0.1) with the genome, GRCz11. The total numbers of sequenced read pairs per sample for RNA and ATAC were between 197,900,000 and 268,400,000. The estimated numbers of cells for WT and mutant were 7,137 and 7,872, respectively. The mean numbers of raw read pairs per cell were (1) 27,742.56 for RNA and 37,593.97 for ATAC in WT and (2) 26,154.78 for RNA and 27,382.86 for ATAC in mutant. The median numbers of genes per cell for WT and mutant were 349 and 365, respectively. ATAC median high-quality fragments per cell for WT and mutant were 10,466 and 8,626, respectively.

For downstream analyses, we used the Weighted Nearest Neighbor (WNN) method in Seurat ^24^. The two experimental conditions of WT and mutant were first analyzed separately. Data filtering was based on visual inspection of data distributions. The number of RNA read counts per cell was filtered between 50 and 3,000 for WT and between 50 and 5,000 for mutant. The number of ATAC read counts per cell was filtered between 500 and 50,000 for WT and between 500 and 80,000 for mutant. The filtering threshold for mitochondrial fractions was 15% for both WT and mutant data. Other parameters were left to default values in Seurat (v4.0.2). The numbers of filtered cells in WT and mutant were 6,942 and 7,740, respectively. The numbers of filtered ATAC peaks in WT and mutant were 164,266 and 167,925, respectively. We then followed the standard Seurat pipelines, with default parameters, for RNA analysis (SCTransform and PCA) and ATAC analysis (TFIDF and SVD) to obtain a WNN graph as a weighted combination of RNA and ATAC data for each of WT and mutant data.

Dimensionality reduction was done by UMAP, clustering by the shared nearest neighbor algorithm, and differential marker identification by Wilcoxon rank sum tests. For analyses of variation in chromatin accessibility and enriched motifs, we used chromVAR (Schep et al., 2017) and all motifs from the JASPAR 2020 database (Fornes et al., 2020). We also performed a merged analysis of the two conditions in a similar way by merging the two datasets using the *merge* function in Seurat. We did not make any correction for batch effects because the two conditions did not show any distinct batch effects on UMAP plots of the merged data. Cell states, or types, were identified by cross-referencing with known markers on ZFIN and 5 dpf datasets from ^61^.

For identification of condition-specific significant ATAC peaks in each cluster, intensity distributions of each peak in WT and mutant were statistically analyzed by the Wilcoxon rank sum and the Kolmogorov-Smirnov (KS) methods using one-tailed tests for each condition. Based on manual inspection of p-value distributions of all peaks, we chose raw p-value thresholds of 0.001 and 0.01 for the Wilcoxon and the KS tests, respectively, to deem peaks to be significant. No p-value correction was performed at this filtering step as a strategy of choice. Those significant peaks were further analyzed to identify enriched motifs as described above. In addition, for those clusters of interest, Clusters 8, 35, 38, 39, and 42, we performed a simulation for the number of significant peaks in each cluster by generating 1,000 random peak intensity datasets by shuffling the intensity values between WT and mutant as many as the number of cells in the cluster in question. This simulation provided empirical null distributions of the number of significant peaks to obtain p-values.

The cells included after filtering from the Seurat analysis were used to perform integrated diffusion and MELD, to keep the analyzed dataset consistent. These new techniques were implemented to analyze the data from a different approach. Integrated diffusion was used to combine multimodal datasets, specifically each cell’s RNA-seq and ATAC-seq data, to create a joint data diffusion operator. The 3D integrated PHATE was computed on this joint data diffusion operator as described previously ^25,26^. To color the plots by likelihood of a cell belonging to the wildtype or mutant sample, this integrated diffusion operator was used for MELD, outputting the likelihood score for each cell belonging to a wildtype or mutant sample.

---

**Extended Data Fig. 1:**
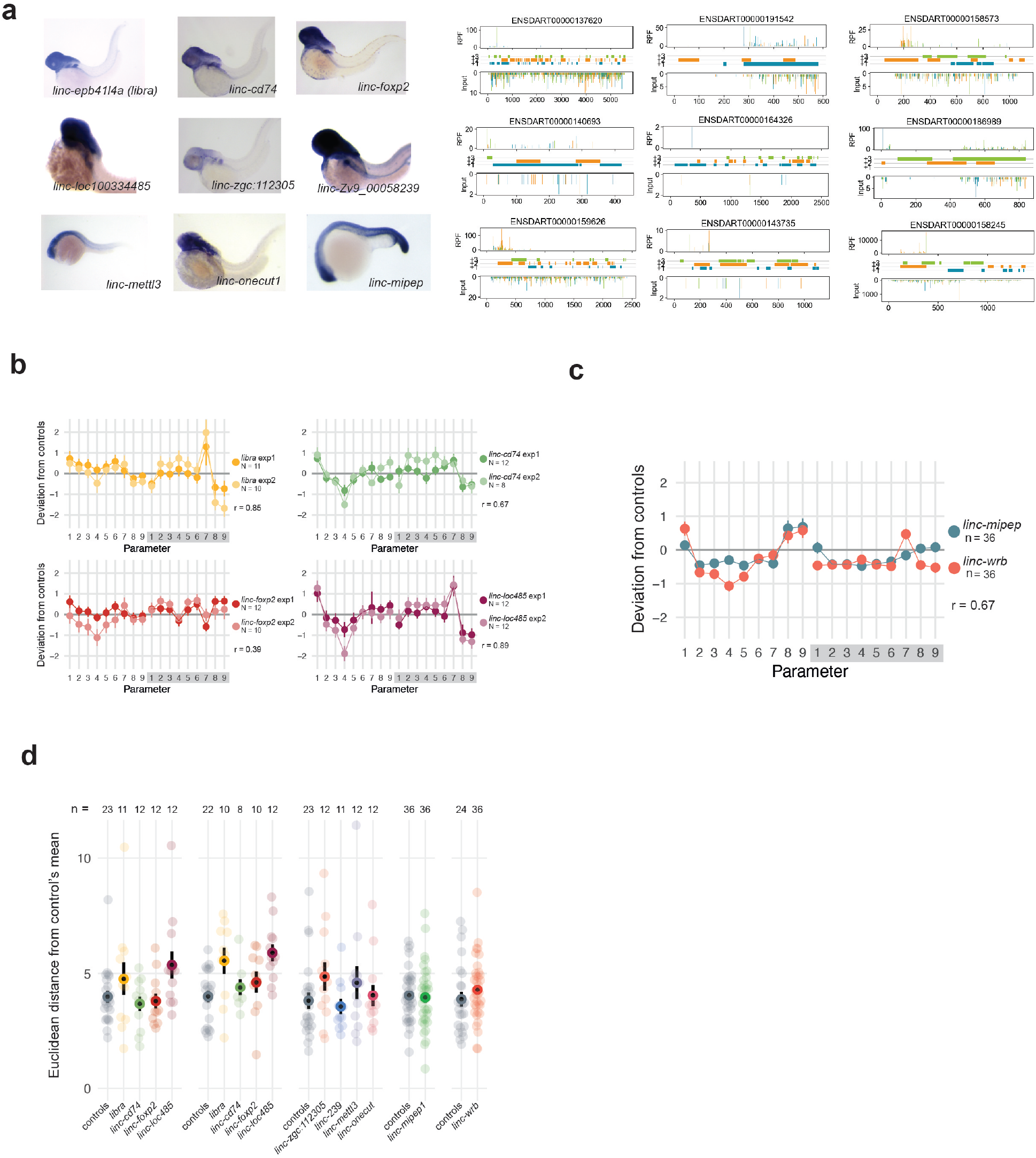
Screening for micropeptide loss-of-function effects on zebrafish baseline behavior. a. Left, *In situ* hybridization screen of zebrafish embryos from 1-4 dpf for transcripts identified as encoding putative micropeptides: *linc-epb41l4a (libra), linc-cd74, linc-foxp2, linc-loc100334485, linc-zgc:112305, linc-Zv9_00058239, linc-mettl3, linc-onecut1*, and *linc-mipep. Right*, corresponding ribosome footprints of transcripts from left panel at 12 hours per fertilization (hpf) across annotated transcript length, with putative coding frames in green (+3), orange (+2), or blue (+1); input on bottom tracks. b. Behavioral fingerprints for independent experiment replicates for *libra, linc-cd74, linc-foxp2*, and *linc-loc100334485* (labeled *linc-loc485*). Deviation (Z-score) of each mutant (F0) larva from the mean of their wild-type siblings across all parameters in day and night. Parameters are as follows: : (1) active bout length (duration of each active bout in seconds); (2) active bout mean (mean of the Δ pixels composing each active bout); (3) active bout standard deviation (mean of the Δ pixels composing each active bout); (4) active bout total (sum of the Δ pixels composing each active bout); (5) active bout minimum (smallest Δ pixels of each bout); (6) active bout maximum (largest Δ pixels of each bout); (7) number of active bouts during the entire day or night; (8) total time active (% of the day or night); (9) inactive bout length (duration of each pause between active bouts in seconds). Exp1, experiment 1. Exp2, experiment 2. r= Pearson’s correlation coefficient between replicate experiments. c. Behavioral fingerprints for *linc-mipep* and *linc-wrb* F0 experiments. Deviation (Z-score) of each mutant (F0) larva from the mean of their wild-type siblings across all parameters, labeled as in c. r= Perason’s correlation coefficient between *linc-mipep* and *linc-wrb* behavioral fingerprints. d. Euclidean distance from control’s mean across 18 behavioral parameters (described in b) for independent F0 experiments targeting putative coding sequence of previously identified lincRNAs. Number of larvae per experiment labeled (n).

**Extended Data Fig. 2:**
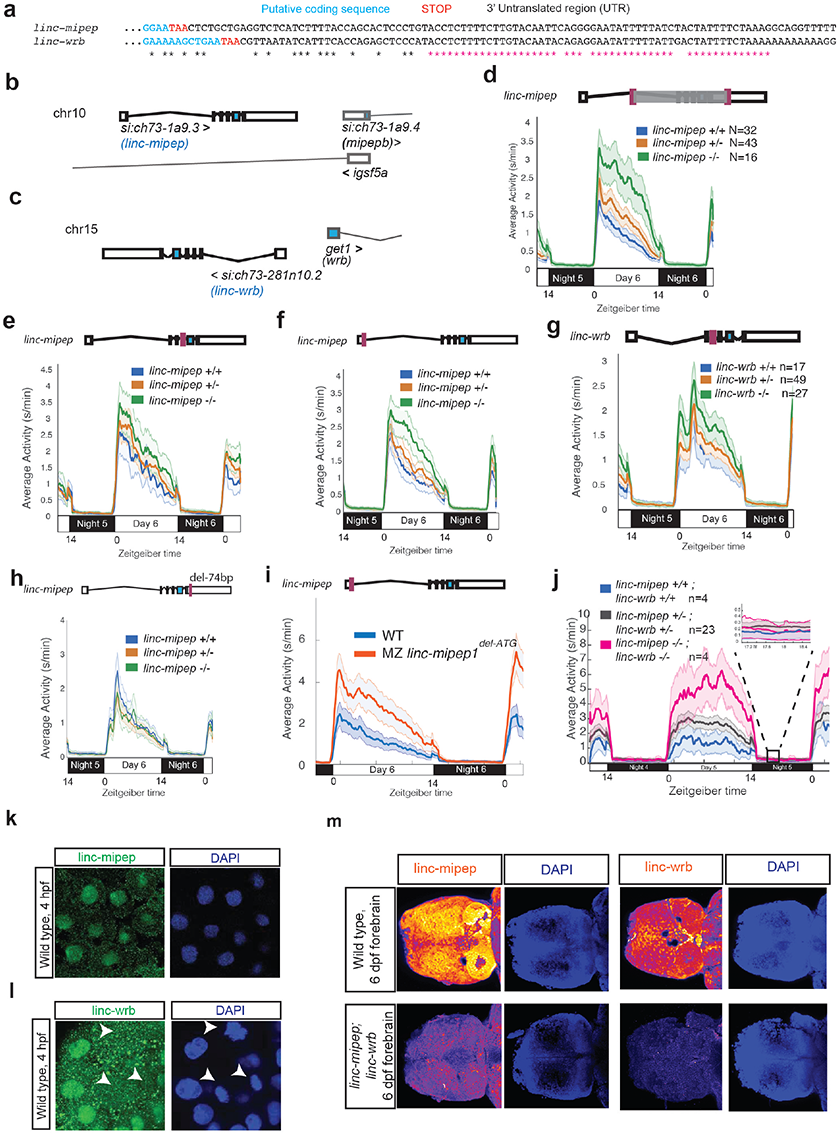
*linc-mipep* and *linc-wrb* loss-of-protein-function mutant larvae are behaviorally hyperactive in a dose-dependent manner. a. A conserved proximal 3’UTR sequence between *linc-mipep* and *linc-wrb* is denoted with magenta asterisks. Black asterisks, conserved nucleotides. Cyan nucleotides, putative coding sequence. Stop codon is denoted in red. 3’ untranslated region (UTR) is in black text. b. Gene location of *linc-mipep*, currently annotated as *si:ch73-1a9.3*, on chromosome 10 (*Danio rerio*). *linc-mipep* contains 6 exons, a 5’UTR and a 3’UTR on the forward strand and is located approximately 3.5kb upstream of *si:ch73-1a9.4 (mipepb). Linc-mipep* lies within intron 1 of *igsf5a* on the reverse (opposite) strand. c. Gene location of *linc-wrb*, currently annotated as *si:ch73-281n10.2*, on chromosome 15 (*Danio rerio*). *Linc-wrb* contains 6 exons, a 5’UTR and a 3’UTR on the reverse strand, and its start site is located approximately 450bp upstream of the *get1* (also known as *wrb*) start site on the forward (opposite) strand. d. Locomotor activity of *linc-mipep*^*del-1.8kb/del-1.8kb*^ (*linc-mipep -/-*, green); *linc-mipep*^*del-1.8kb/+*^ (*linc-mipep +/-*, orange); and wild-type (*linc-mipep +/+*, blue) sibling-matched larvae over 2 nights. The ribbon represents +/-SEM. Zeitgeber time is defined from lights ON=0. Schematic of mutation is above plot, with gray shade indicating the deletion. e. Locomotor activity of *linc-mipep*^*del8bp/del8bp*^ (*linc-mipep -/-*, green); *linc-mipep*^*del8bp /+*^ (*linc-mipep +/-*, orange); and wild-type (*linc-mipep +/+*, blue) sibling-matched larvae over 2 nights. Schematic of mutation is above plot, with mutation location indicated in purple. f. Locomotor activity of *linc-mipep*^*delATG/delATG*^ (*linc-mipep -/-*, green); *linc-mipep*^*delATG /+*^ (*linc-mipep +/-*, orange); and wild-type (*linc-mipep +/+*, blue) sibling-matched larvae over 2 nights. Schematic of mutation is above plot, with mutation location indicated in purple. g. Locomotor activity of *linc-wrb*^*del-11/del-11*^ (*linc-wrb -/-*, green); *linc-wrb*^*del-11/+*^ (*linc-wrb +/-*, orange); and wild-type (*linc-wrb +/+*, blue) sibling-matched larvae over 2 nights. Schematic of mutation is above plot. h. Locomotor activity of *linc-mipep*^*3’UTR-74bpdel /3’UTR-74bpdel*^ (*linc-mipep -/-*, green); *linc-mipep*^*3’UTR-74bpdel /+*^ (*linc-mipep +/-*, orange); and wild-type (*linc-mipep +/+*, blue) sibling-matched larvae over 2 nights. Schematic of mutation is above plot, with mutation location indicated in purple. i. Locomotor activity of wild type (WT, blue) or maternal-zygotic *linc-mipep* ^*delATG/delATG*^ (MZ *linc-mipep* ^*delATG*^, orange) larvae over 24 hours. j. Locomotor activity of *linc-mipep*^*del-1.8kb/del-1.8kb*^ *linc-wrb*^*del-11/del-11*^ (*linc-mipep -/-; linc-wrb -/-*, magenta); *linc-mipep*^*del-1.8kb/+*^ *linc-wrb*^*del-11/+*^ (*linc-mipep +/-; linc-wrb +/-*, black); and wild-type (*linc-mipep +/+; linc-wrb +/+*, blue) sibling-matched larvae over 2 nights. The magnified activity profile on night 5 is shown. k. Confocal images of *linc-mipep* protein (green) and DAPI (nuclei, blue), in 4 hpf zebrafish embryos. l. Confocal images of *linc-wrb* protein (green) and DAPI (nuclei, blue), in 4 hpf Arrows indicate mitotic nuclei, which show no *linc-wrb* antibody staining. m. Confocal images of *linc-mipep* (orange, intensity by depth) and DAPI (nuclei, blue), left, or of *linc-wrb* (orange, intensity by depth) and DAPI (nuclei, blue), right, in 6 dpf zebrafish forebrains (dorsal view), in wild type (WT, top) or *linc-mipep; linc-wrb* double mutants (bottom).

**Extended Data Fig. 3:**
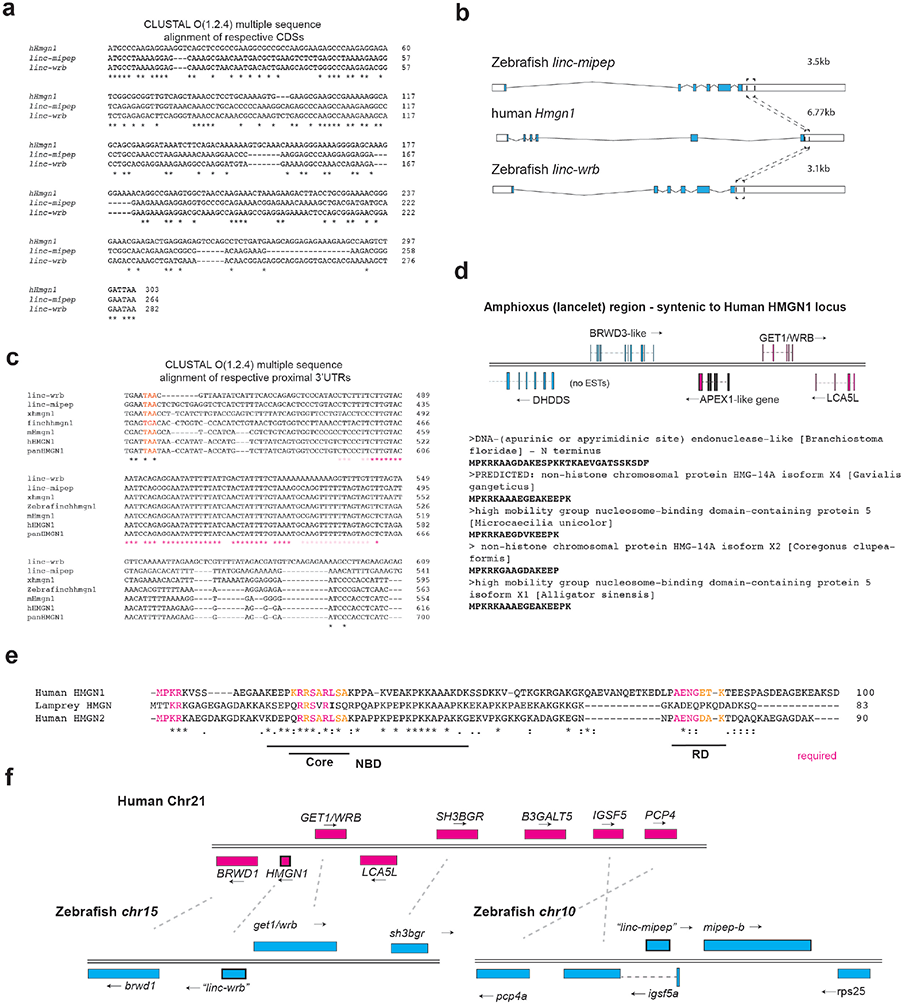
*linc-mipep* and *linc-wrb* encode proteins with homology to human HMGN1. a. Multiple sequence alignment of cDNA sequences of human *Hmgn1* (*hHmgn1*), *linc-mipep*, and *linc-wrb*. Asterisks, nucleotide conservation across species. b. Transcripts for *linc-mipep* (top), *linc-wrb* (bottom), and human *Hmgn1* (middle), normalized for scale. Transcript length denoted on top right of each. A conserved proximal 3’UTR sequence across species is denoted with boxes and dotted lines. c. Multiple sequence alignment of 3’UTR sequences of *linc-mipep*, and *linc-wrb*, and *Hmgn1* across select species. Gray, coding sequence. Red, stop codon. Pink or magenta asterisks, partial or full nucleotide conservation across species. *Linc-wrb* and *linc-mipep*, zebrafish proteins with homology to *HMGN1*. Xhmgn1, *Xenopus tropicalis*. finchhmgn1, zebra finch. mHmgn1, mouse. hHmgn1, human. panHMGN1, chimpanzee (*Pan troglodytes*). d. Identification of a gene syntenic to human *HMGN1* (chromosome 21) in the invertebrate lancelet (Amphioxus) genome. The APEX1-like gene N terminus BLASTs to HMGN genes, shown here for select species. e. Multiple sequence alignment of ancestral sea lamprey putative HMGN1 ORF and human HMGN1 (top) and HMGN2 (bottom). NBD, nucleosome binding domain (with core indicated). RD, regulatory domain. Amino acids functionally required (magenta) or conserved in HMGN1 or HMGN2 lineages (orange) as indicated. f. Syntenic alignment of human chromosome 21 and zebrafish chromosomes 15 and 10. Top: human chromosome 21 (at q22.2, Ensembl GRCh38.p13), showing annotations for *BRWD1, HMGN1, GET1/WRB, SH3BGR, B3GALT5, IGSF5, and PCP4*, in magenta. Bottom: zebrafish chromosome 15 (left) or 10 (right), centered on *linc-wrb;* left, zebrafish chromosome 10, centered at *linc-mipep*, in cyan. Gray dotted lines show synteny between human and zebrafish genes.

**Extended Data Fig. 4:**
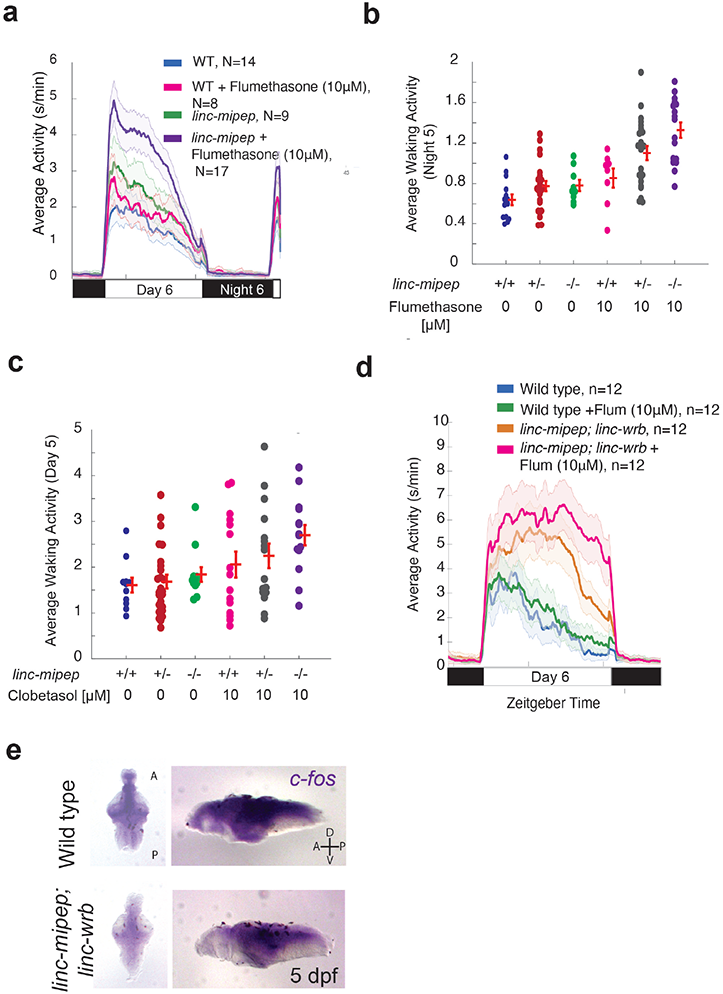
*linc-mipep* mutants have dysregulation of NMDA receptor-mediated signaling, glucocorticoid signaling, and immediate early gene induction. a. Locomotor average activity of wild-type larvae treated with DMSO (WT, blue) or with 10mM glucocorticoid receptor agonist Flumethasone (magenta), and *linc-mipep* ^*del-1.8kb/del-1.8kb*^ larvae treated with DMSO (*linc-mipep*, green) or with 10mM Flumethasone (purple); sibling-matched larvae over 24 hours. b. Average waking activity (Night 5) of progeny of incrosses of *linc-mipep* ^*del-1.8kb/+*^ fish treated with DMSO or 10mM Flumethasone. c. Average waking activity (Day 5) of progeny of incrosses of *linc-mipep* ^*del-1.8kb/+*^ fish treated with DMSO or 10mM Clobetasol (glucocorticoid receptor agonist). d. Average locomotor activity at 6 dpf of wild type larvae treated with DMSO (WT, blue) or with 10mM Flumethasone (Flum) (green), and *linc-mipep;linc-wrb* double mutants treated with DMSO (orange) or 10mM Flumethasone (magenta). e. *In situ* hybridization of *c-fos* expression in 5 dpf wild type (top) and *linc-mipep; linc-wrb* (bottom) larval brains. Dorsal views, left; lateral views, right. A, anterior; P, posterior; D, dorsal; V, ventral.

**Extended Data Fig. 5:**
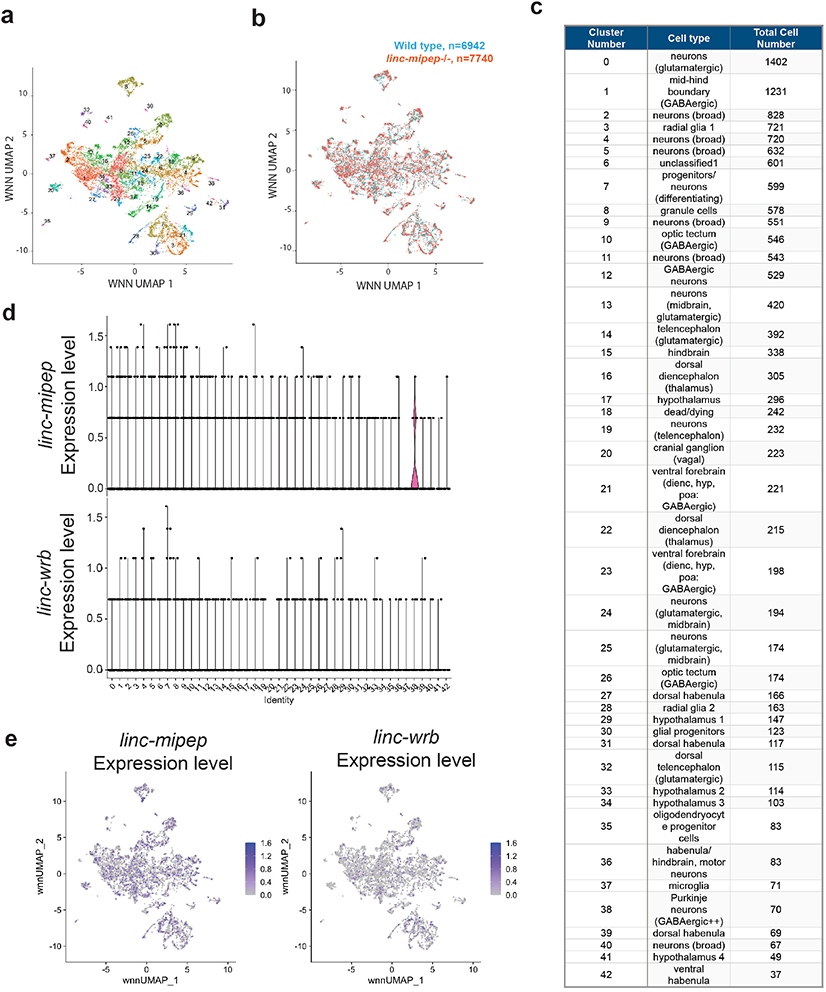
Single cell Multiome analyses in wild type and *linc-mipep* mutant brain nuclei. a. UMAP representation of WNN analyses (transcriptomic and chromatin accessibility) of merged wild type (n=6942 nuclei) and *linc-mipep* ^*del-1.8kb del-1.8kb*^ (n=7740 nuclei) mutant brains at 6 days post-fertilization (dpf), labeled with cell cluster numbers. b. UMAP representation of WNN analyses of merged wild type (n=6942 nuclei, cyan) and *linc-mipep* ^*del-1.8kb del-1.8kb*^ (n=7740 nuclei, orange) mutant brains at 6 days post-fertilization (dpf). c. Table of WNN cluster identification numbers, classified cell type, and total cell number per cluster. d. Violin plots of *linc-mipep* (top) and *linc-wrb* (bottom) expression levels from WNN analysis clusters of wild type brain nuclei (n=6942) at 6 days post-fertilization (dpf). e. UMAP representation of WNN analyses of wild type brain nuclei (n=6942) at 6 days post-fertilization (dpf), color-coded by relative expression levels (purple scale) of *linc-mipep* (left) and *linc-wrb* (right).

**Extended Data Fig. 6:**
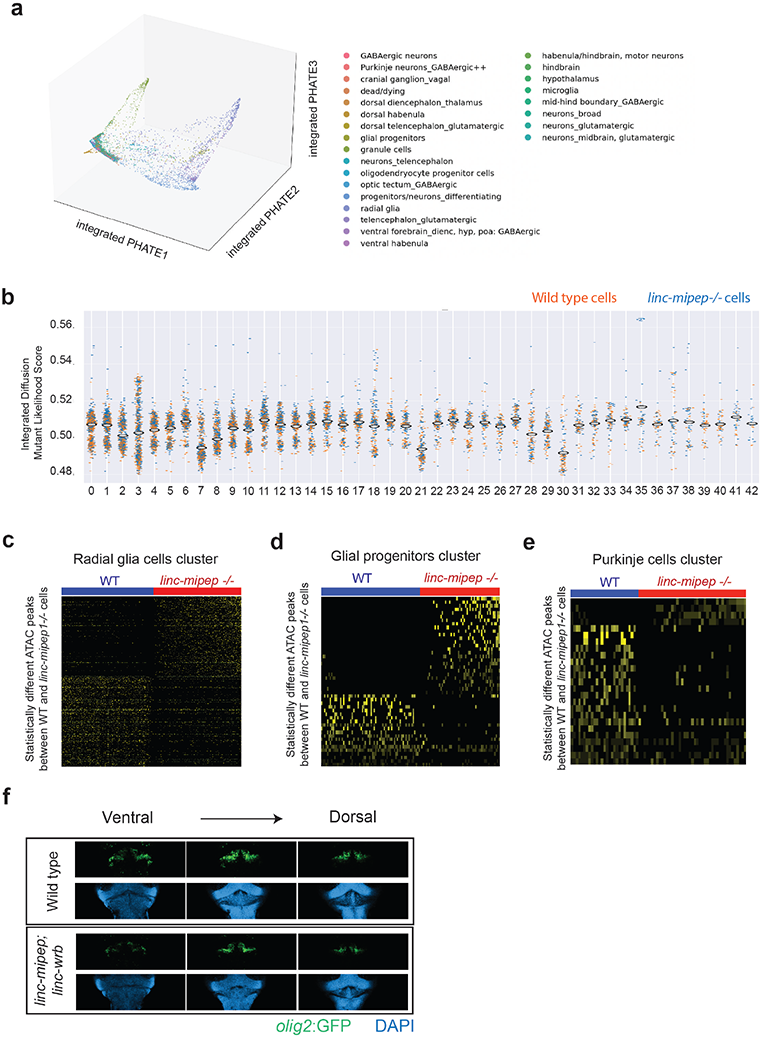
Single cell Multiome analyses reveal cell states altered in *linc-mipep* brain cells. a. PHATE plot of integrated diffusion analysis of 6 dpf wild type and *linc-mipep* ^*del-1.8kb del-1.8kb*^ mutant brain nuclei, color-coded by broad identified cell type. b. Integrated diffusion analysis on identified cell type clusters from 6 dpf wild type (orange) and *linc-mipep* ^*del-1.8kb del-1.8kb*^ (blue) brain nuclei by WNN-identified clusters. Each dot represents a single cell, with mutant likelihood score across Y-axis. c. Statistically significantly different chromatin accessibility peaks between 6 dpf wild type (WT, blue) and *linc-mipep* ^*del-1.8kb del-1.8kb*^ mutant (red) nuclei in the radial glial cells cluster (#3). d. Statistically significantly different chromatin accessibility peaks between 6 dpf wild type (WT, blue) and *linc-mipep* ^*del-1.8kb del-1.8kb*^ mutant (red) nuclei in the glial progenitor cells cluster (#30). e. Statistically significantly different chromatin accessibility peaks between 6 dpf wild type (WT, blue) and *linc-mipep* ^*del-1.8kb del-1.8kb*^ mutant (red) nuclei in the Purkinje cells cluster (#38). f. Dorsal view confocal images (comparable single Z planes) from *Tg(olig2:GFP)* brains zoomed in on the cerebellum in wild type (top) or *linc-mipep;linc-wrb* double heterozygous mutant (bottom) backgrounds, stained with GFP (*olig2+*, green) and DAPI (nuclei, blue), from ventral to dorsal. Anterior to the top.

**Extended Data Fig. 7:**
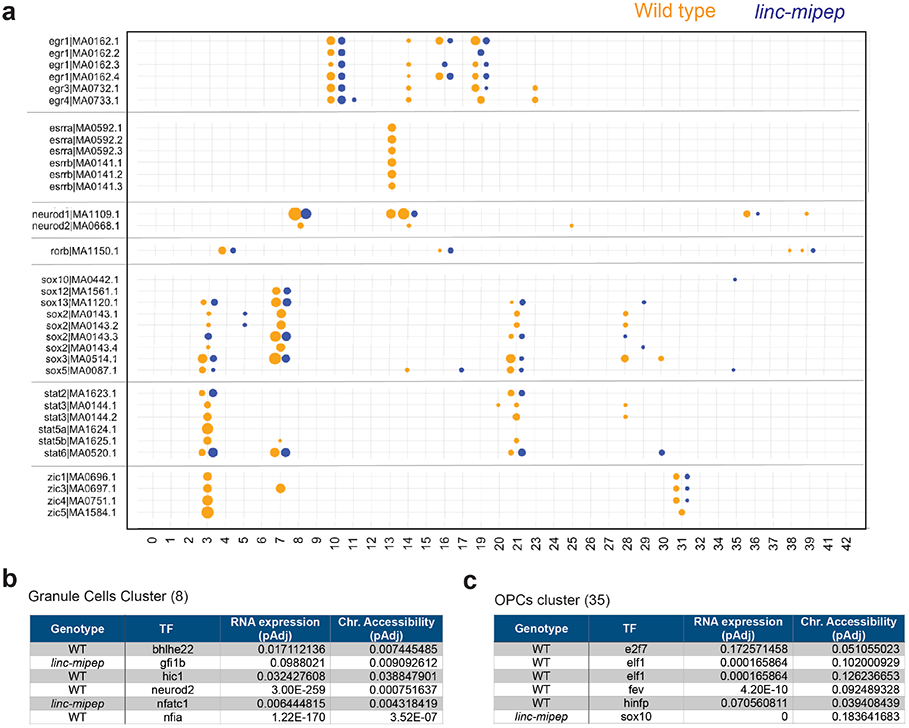
Accessibility for transcription factor motifs most affected in *linc-mipep* brain cells. a. Accessibility for transcription factor motifs, ordered by related family members, that are significantly differentially accessible per WNN cluster (as described in Fig. 4d) in either wild type or *linc-mipep* brain cells. Wild type, orange. *Linc-mipep*, blue. Circle sizes represent adjusted p-values between 2.87e-121 and 0.198 in log scale. b. Accessibility for transcription factor motifs significantly different between wild type or *linc-mipep* granule cells cluster (#8). Difference in RNA expression (adjusted P value) and chromatin accessibility (adjusted P value) for each TF (gene or motif) as shown. c. Accessibility for transcription factor motifs significantly different between wild type or *linc-mipep* OPCs cluster (#35). Difference in RNA expression (adjusted P value) and chromatin accessibility (adjusted P value) for each TF (gene or motif) as shown.

## List of Supplementary Tables

**Supplementary Table 1:** Information on sORFs identified within lincRNAs and targeting/genotyping information (3 sheets)

**Supplementary Table 2:** Protein and proximal 3’UTR BLAST results, and related HMGN1 across species (5 sheets)

**Supplementary Table 3:** Correlating Drugs to *linc-mipep1* heterozygous and homozygous mutants, from hierarchical clustering analysis against >500 FDA-approved small molecues (from Rihel et al., 2010), and concentrations used (1 sheet)

**Supplementary Table 4:** Bulk omni-ATAC-seq on WT or *linc-mipep;linc-wrb* mutant brains at 5 dpf (4 sheets)

**Supplementary Table 5:** Supplementary Table 5 - Single cell Multiome Analyses of WT or linc-mipep mutant brains (sibling-matched) at 5 dpf (8 sheets)

**Supplementary Table 6:** Integrated Diffusion/MELD analyses using WNN clusters and conditional clusters (2 sheets)

**Supplementary Table 7:** RNA Polymerase II ChIP-seq on wild type (WT) or *linc-mipep; linc-wrb* dissected brains at 5 days post-fertilization (dpf) (2 sheets)

**Supplementary Table 8:** Genes identified by ChIP-Seq for Linc-mipep1; Linc-wrb on wild type 24 hpf embryos (1 sheet)

## List of Supplementary Figures

**Supplementary Figure 1:** ATAC peak intensity plots for statistically different peaks between wild type and *linc-mipep* mutant cells.

## References

1. Goudarzi, M., Berg, K., Pieper, L. M. & Schier, A. F. Individual long non-coding RNAs have no overt functions in zebrafish embryogenesis, viability and fertility. eLife 8, e40815 (2019).

2. Bazzini, A. A. et al. Identification of small ORFs in vertebrates using ribosome footprinting and evolutionary conservation. EMBO J. 33, 981–993 (2014).

3. Chen, J. et al. Pervasive functional translation of non-canonical human open reading frames. Science 367, 1140–1146 (2020).

4. Ingolia, N. T., Ghaemmaghami, S., Newman, J. R. S. & Weissman, J. S. Genome-wide analysis in vivo of translation with nucleotide resolution using ribosome profiling. Science 324, 218–223 (2009).

5. Kondo, T. et al. Small peptide regulators of actin-based cell morphogenesis encoded by a polycistronic mRNA. Nat. Cell Biol. 9, 660–665 (2007).

6. Pauli, A. et al. Toddler: An Embryonic Signal That Promotes Cell Movement via Apelin Receptors. Science 343, 1248636 (2014).

7. Couso, J.-P. & Patraquim, P. Classification and function of small open reading frames. Nat. Rev. Mol. Cell Biol. 18, 575–589 (2017).

8. Bi, P. et al. Control of muscle formation by the fusogenic micropeptide myomixer. Science 356, 323–327 (2017).

9. D’Lima, N. G. et al. A human microprotein that interacts with the mRNA decapping complex. Nat. Chem. Biol. 13, 174–180 (2017).

10. Fields, A. P. et al. A Regression-Based Analysis of Ribosome-Profiling Data Reveals a Conserved Complexity to Mammalian Translation. Mol. Cell 60, 816–827 (2015).

11. Derrien, T. et al. The GENCODE v7 catalog of human long noncoding RNAs: analysis of their gene structure, evolution, and expression. Genome Res. 22, 1775–1789 (2012).

12. Ulitsky, I., Shkumatava, A., Jan, C. H., Sive, H. & Bartel, D. P. Conserved function of lincRNAs in vertebrate embryonic development despite rapid sequence evolution. Cell 147, 1537–1550 (2011).

13. Makarewich, C. A. & Olson, E. N. Mining for Micropeptides. Trends Cell Biol. 27, 685–696 (2017).

14. Prober, D. A., Rihel, J., Onah, A. A., Sung, R.-J. & Schier, A. F. Hypocretin/Orexin Overexpression Induces An Insomnia-Like Phenotype in Zebrafish. J. Neurosci. 26, 13400–13410 (2006).

15. Rihel, J. et al. Zebrafish Behavioral Profiling Links Drugs to Biological Targets and Rest/Wake Regulation. Science 327, 348–351 (2010).

16. The Behavioral Repertoire of Larval Zebrafish | Springer Nature Experiments. https://experiments.springernature.com/articles/10.1007/978-1-60761-922-2_12.

17. Kroll, F. et al. A simple and effective F0 knockout method for rapid screening of behaviour and other complex phenotypes. eLife 10, e59683 (2021).

18. Bustin, M. Revised nomenclature for high mobility group (HMG) chromosomal proteins. Trends Biochem. Sci. 26, 152–153 (2001).

19. Kumar, S. & Hedges, S. B. A molecular timescale for vertebrate evolution. Nature 392, 917–920 (1998).

20. Cuddapah, S. et al. Genomic Profiling of HMGN1 Reveals an Association with Chromatin at Regulatory Regions. Mol. Cell. Biol. 31, 700–709 (2011).

21. Deng, T. et al. HMGN1 Modulates Nucleosome Occupancy and DNase I Hypersensitivity at the CpG Island Promoters of Embryonic Stem Cells. Mol. Cell. Biol. 33, 3377–3389 (2013).

22. Sheng, M. & Greenberg, M. E. The regulation and function of c-fos and other immediate early genes in the nervous system. Neuron 4, 477–485 (1990).

23. Bialkowska, A. B., Yang, V. W. & Mallipattu, S. K. Krüppel-like factors in mammalian stem cells and development. Dev. Camb. Engl. 144, 737–754 (2017).

24. Hao, Y. et al. Integrated analysis of multimodal single-cell data. Cell 184, 3573-3587.e29 (2021).

25. Kuchroo, M. et al. Multiscale PHATE identifies multimodal signatures of COVID-19. Nat. Biotechnol. 40, 681–691 (2022).

26. Kuchroo, M., Godavarthi, A., Tong, A., Wolf, G. & Krishnaswamy, S. Multimodal Data Visualization and Denoising with Integrated Diffusion. in 2021 IEEE 31st International Workshop on Machine Learning for Signal Processing (MLSP) 1–6 (2021). doi:10.1109/MLSP52302.2021.9596214.

27. Lamanna, F. et al. Reconstructing the ancestral vertebrate brain using a lamprey neural cell type atlas. 2022.02.28.482278 Preprint at https://doi.org/10.1101/2022.02.28.482278 (2022).

28. Miyata, T., Maeda, T. & Lee, J. E. NeuroD is required for differentiation of the granule cells in the cerebellum and hippocampus. Genes Dev. 13, 1647–1652 (1999).

29. Hashimoto, R. et al. Origins of oligodendrocytes in the cerebellum, whose development is controlled by the transcription factor, Sox9. Mech. Dev. 140, 25–40 (2016).

30. Martik, M. L. & Bronner, M. E. Regulatory Logic Underlying Diversification of the Neural Crest. Trends Genet. 33, 715–727 (2017).

31. Simões-Costa, M., Tan-Cabugao, J., Antoshechkin, I., Sauka-Spengler, T. & Bronner, M. E. Transcriptome analysis reveals novel players in the cranial neural crest gene regulatory network. Genome Res. 24, 281–290 (2014).

32. Cui, H. et al. Behavioral Disturbances in Estrogen-Related Receptor alpha-Null Mice. Cell Rep. 11, 344–350 (2015).

33. Hoshino, M. et al. Ptf1a, a bHLH Transcriptional Gene, Defines GABAergic Neuronal Fates in Cerebellum. Neuron 47, 201–213 (2005).

34. Gans, C. & Northcutt, R. G. Neural Crest and the Origin of Vertebrates: A New Head. Science 220, 268–273 (1983).

35. Deng, T. et al. Interplay between H1 and HMGN epigenetically regulates OLIG1&2 expression and oligodendrocyte differentiation. Nucleic Acids Res. 45, 3031–3045 (2017).

36. González-Romero, R., Eirín-López, J. M. & Ausió, J. Evolution of High Mobility Group Nucleosome-Binding Proteins and Its Implications for Vertebrate Chromatin Specialization. Mol. Biol. Evol. 32, 121–131 (2015).

37. Hock, R., Furusawa, T., Ueda, T. & Bustin, M. HMG chromosomal proteins in development and disease. Trends Cell Biol. 17, 72–79 (2007).

38. Ihewulezi, C. & Saint-Jeannet, J.-P. Function of chromatin modifier Hmgn1 during neural crest and craniofacial development. Genes. N. Y. N 2000 59, e23447 (2021).

39. Zalc, B. The acquisition of myelin: An evolutionary perspective. Brain Res. 1641, 4–10 (2016).

40. Zalc, B., Goujet, D. & Colman, D. The origin of the myelination program in vertebrates. Curr. Biol. CB 18, R511–512 (2008).

41. Baxter, L. L., Moran, T. H., Richtsmeier, J. T., Troncoso, J. & Reeves, R. H. Discovery and genetic localization of Down syndrome cerebellar phenotypes using the Ts65Dn mouse. Hum. Mol. Genet. 9, 195–202 (2000).

42. Mowery, C. T. et al. Trisomy of a Down Syndrome Critical Region Globally Amplifies Transcription via HMGN1 Overexpression. Cell Rep. 25, 1898-1911.e5 (2018).

43. Olmos-Serrano, J. L. et al. Down Syndrome Developmental Brain Transcriptome Reveals Defective Oligodendrocyte Differentiation and Myelination. Neuron 89, 1208–1222 (2016).

44. Jin, X. et al. In vivo Perturb-Seq reveals neuronal and glial abnormalities associated with autism risk genes. Science 370, eaaz6063 (2020).

45. Abuhatzira, L., Shamir, A., Schones, D. E., Schäffer, A. A. & Bustin, M. The Chromatin-binding Protein HMGN1 Regulates the Expression of Methyl CpG-binding Protein 2 (MECP2) and Affects the Behavior of Mice. J. Biol. Chem. 286, 42051–42062 (2011).

46. Sathyanesan, A. et al. Emerging connections between cerebellar development, behavior, and complex brain disorders. Nat. Rev. Neurosci. 20, 298–313 (2019).

47. Weisman, C. M. The Origins and Functions of De Novo Genes: Against All Odds? J. Mol. Evol. (2022) doi:10.1007/s00239-022-10055-3.

48. Bitetti, A. et al. MicroRNA degradation by a conserved target RNA regulates animal behavior. Nat. Struct. Mol. Biol. 25, 244–251 (2018).

49. Barlow, I. L. et al. A genetic screen identifies dreammist as a regulator of sleep. 2020.11.18.388736 Preprint at https://doi.org/10.1101/2020.11.18.388736 (2020).

50. Moreno-Mateos, M. A. et al. CRISPRscan: designing highly efficient sgRNAs for CRISPR-Cas9 targeting in vivo. Nat. Methods 12, 982–988 (2015).

51. Ghosh, M. & Rihel, J. Hierarchical Compression Reveals Sub-Second to Day-Long Structure in Larval Zebrafish Behavior. eNeuro 7, ENEURO.0408-19.2020 (2020).

52. Hoffman, E. J. et al. Estrogens Suppress a Behavioral Phenotype in Zebrafish Mutants of the Autism Risk Gene, CNTNAP2. Neuron 89, 725–733 (2016).

53. Shin, J., Park, H.-C., Topczewska, J. M., Mawdsley, D. J. & Appel, B. Neural cell fate analysis in zebrafish using olig2 BAC transgenics. Methods Cell Sci. Off. J. Soc. Vitro Biol. 25, 7–14 (2003).

54. Randlett, O. et al. Whole-brain activity mapping onto a zebrafish brain atlas. Nat. Methods 12, 1039–1046 (2015).

55. Trinh, L. A. et al. Biotagging of Specific Cell Populations in Zebrafish Reveals Gene Regulatory Logic Encoded in the Nuclear Transcriptome. Cell Rep. 19, 425–440 (2017).

56. Cotney, J. L. & Noonan, J. P. Chromatin Immunoprecipitation with Fixed Animal Tissues and Preparation for High-Throughput Sequencing. Cold Spring Harb. Protoc. 2015, 419 (2015).

57. Miao, L. et al. The landscape of pioneer factor activity reveals the mechanisms of chromatin reprogramming and genome activation. Mol. Cell 82, 986-1002.e9 (2022).

58. Buenrostro, J. D., Giresi, P. G., Zaba, L. C., Chang, H. Y. & Greenleaf, W. J. Transposition of native chromatin for fast and sensitive epigenomic profiling of open chromatin, DNA-binding proteins and nucleosome position. Nat. Methods 10, 1213–1218 (2013).

59. Corces, M. R. et al. An improved ATAC-seq protocol reduces background and enables interrogation of frozen tissues. Nat. Methods 14, 959–962 (2017).

60. Vejnar, C. E. & Giraldez, A. J. LabxDB: versatile databases for genomic sequencing and lab management. Bioinforma. Oxf. Engl. 36, 4530–4531 (2020).

61. Raj, B. et al. Emergence of Neuronal Diversity during Vertebrate Brain Development. Neuron 108, 1058-1074.e6 (2020).

62. Bradford, Y. M. et al. Zebrafish information network, the knowledgebase for Danio rerio research. Genetics 220, iyac016 (2022).

