## SupplementaryNote1 for "*linc-mipep* and *linc-wrb* encode micropeptides that regulate chromatin accessibility in vertebrate-specific neural cells"

**Supplementary Note 1**

The protein-coding potential and function of transcripts previously identified as noncoding remain unknown. There are two nomenclature issues in the literature specifically for *linc-mipep.*

1) *Nomenclature:* Multiple names for the same transcripts can be found in the literature. In Ulitsky et al. (2011) and Bazzini et al. (2014), *linc-mipep* was the name given to the transcript that is now annotated as *si:ch73-19.3* (Ensembl zv9 coordinates chr10:40,425,061-40,428,902), while Wang et al (2017) and Wang et al., (2021) named this transcript *lnc-rps25.* The downstream gene annotated at that time was named *mipepb.* Thus, we have chosen to use *linc-mipep* for this transcript.

2) *Nomenclature of ultra-conserved 3'UTR sequence identified:* In our study, we identify a short ultra-conserved element in the 3'UTR sequences of *linc-mipep* and *linc-wrb* that correspond to a conserved region in the 3'UTR of human Hmgn1. This region is distinct from the conserved 3'UTR region between *linc-mipep* (also called *lnc-rps25*) and *linc-epb41l4a* (also called *libra and* currently annotated as *si:ch73-46j18.5*, ENSDARG00000092467), identified in Ulitsky et al. 2011 and Bitetti et al., 2018. Both studies showed this region is conserved in the 3'UTR sequence of a gene encoding neuronal protein 3.1 (P311), later called NREP, in human, mouse, and chicken (Figure 7B in Ulitsky et al.; Figure 1A in Bitetti et al., 2018). Interestingly, in our study, we do find that part of the ultra-conserved sequence we identify in *linc-mipep* and *linc-wrb* 3'UTRs (corresponding to human *Hmgn1* 3'UTR) is also found in Hmgn1 pseudogene copies, which comprises one of the largest families of retroposed copies in the human (and mouse) genome (see González-Romero, Eirín-López, & Ausió 2014) (as described in Extended Data Fig. 3b,c and Supplementary Table 2). Specifically, HMGN1P14 (and the identified conserved sequence) is located within intron 2/3 of NREP, yet remains a separate gene unrelated to *Hmgn1* and *linc-mipep*.

**References**

Bitetti A, Mallory AC, Golini E, Carrieri C, Carreño Gutiérrez H, Perlas E, Pérez-Rico YA, Tocchini-Valentini GP, Enright AJ, Norton WHJ, Mandillo S, O'Carroll D, Shkumatava A (2018). MicroRNA degradation by a conserved target RNA regulates animal behavior. *Nat Struct Mol Biol* 25 (3): 244–251.

Gao T, Li J, Li N, Gao Y, Yu L, Zhuang S, Zhao Y, Dong X (2020). *lncrps25* play an essential role in motor neuron development through controlling the expression of *olig2* in zebrafish. *J Cell Physiol*. 2020; 235: 3485– 3496.

González-Romero R, Eirín-López JM, Ausió J (2015). Evolution of High Mobility Group Nucleosome-Binding Proteins and Its Implications for Vertebrate Chromatin Specialization. *Molecular Biology and Evolution* 32(1): 121-131.

Ulitsky I, Shkumatava A, Jan CH, Sive H, Bartel DP (2011). Conserved Function of lincRNAs in Vertebrate Embryonic Development despite Rapid Sequence Evolution. *Cell* 147(7): 1537-1550.

Wang L, Ma X, Xu X, Zhang Y (2017). Systematic identification and characterization of cardiac long intergenic noncoding RNAs in zebrafish. *Sci Rep* **7,** 1250

Wang L, Song F, Zhu W, Fu J, Dong Z, Xu P (2021). The stage-specific long non-coding RNAs and mRNAs identification and analysis during early development of common carp, Cyprinus carpio. *Genomics* 113(1): 20-28.
