## SupplementalFig1 for "*linc-mipep* and *linc-wrb* encode micropeptides that regulate chromatin accessibility in vertebrate-specific neural cells"

**Supplementary Figure 1.** ATAC peak intensity plots for statistically different peaks between wild type and *linc-mipep1* cells. Per cluster: top, heat-map of chromatin accessibility by peak (blue scale increasing in accessibility); bottom, P value of statistically different chromatin accessibility peaks between wild type and mutant cells, using Wilcoxon rank sum (red) and the Kolmogorov-Smirnov (blue) methods using one-tailed tests for each condition. Raw p-value thresholds of 0.001 and 0.01 for the Wilcoxon and the KS tests, respectively, were deemed to be significant peaks for further analysis.

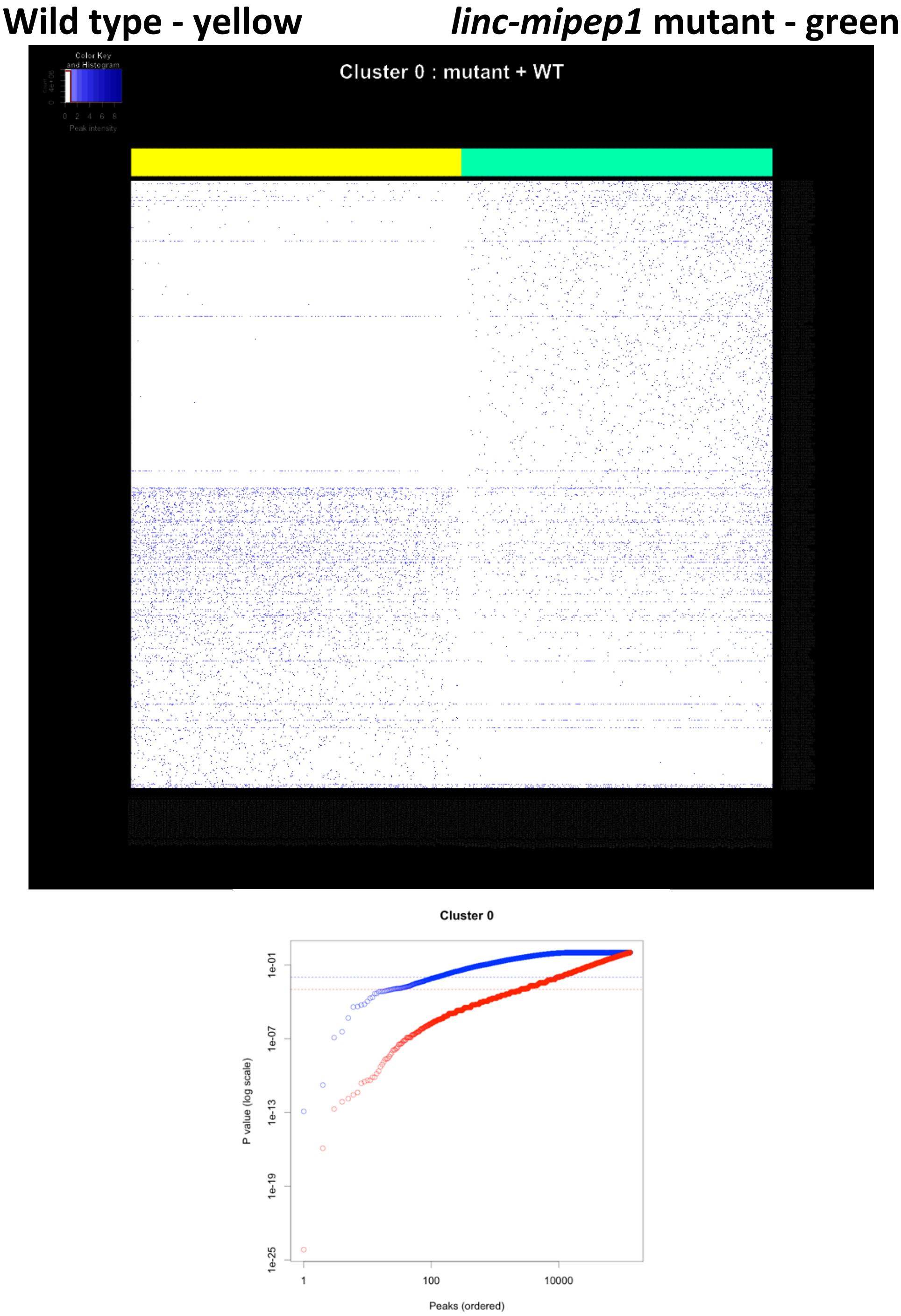

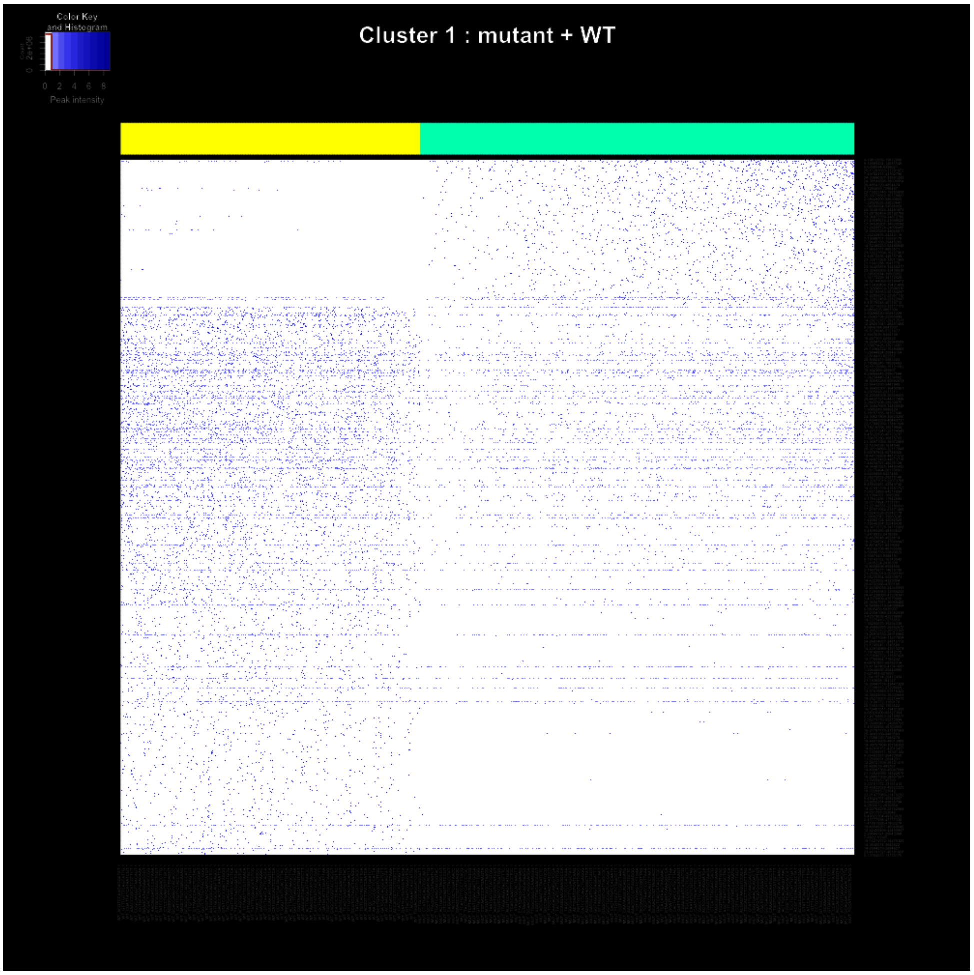

Color Key and Histogram
Peak intensity
0 2 4 6 8
Cluster 1 : mutant + WT


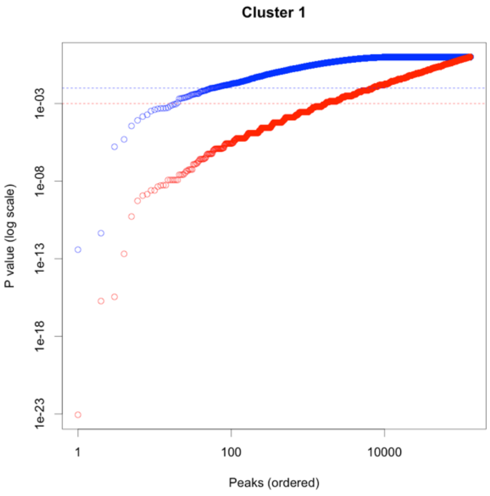

Cluster 1
P value (log scale)
1e-03
1e-08
1e-13
1e-18
1e-23
1
100
10000
Peaks (ordered)

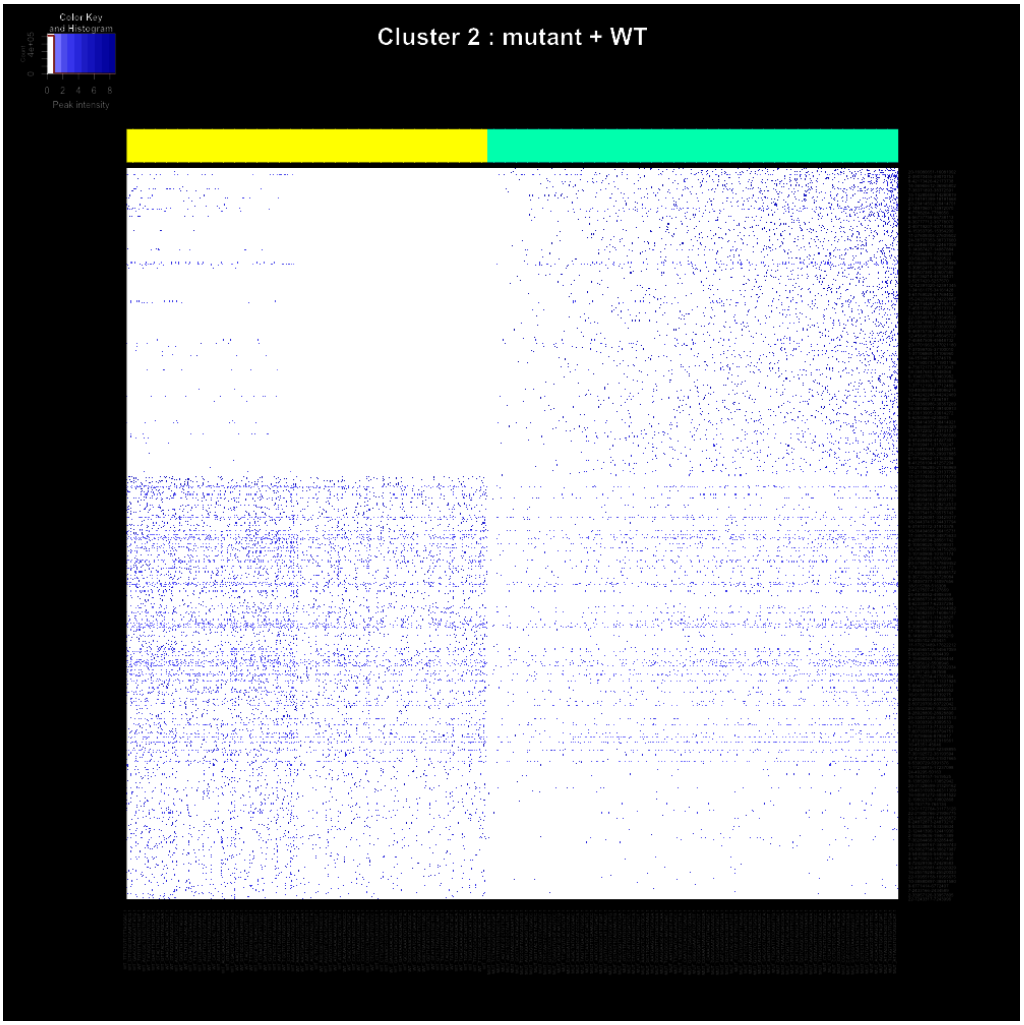
Color Key
and Histogram
0 2 4 6 8
Peak intensity
Cluster 2 : mutant + WT

Cluster 2

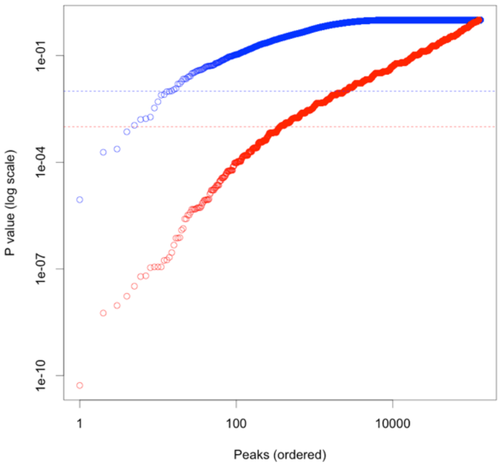
1e-01
1e-04
1e-07
1e-10
P value (log scale)
1
100
10000
Peaks (ordered)

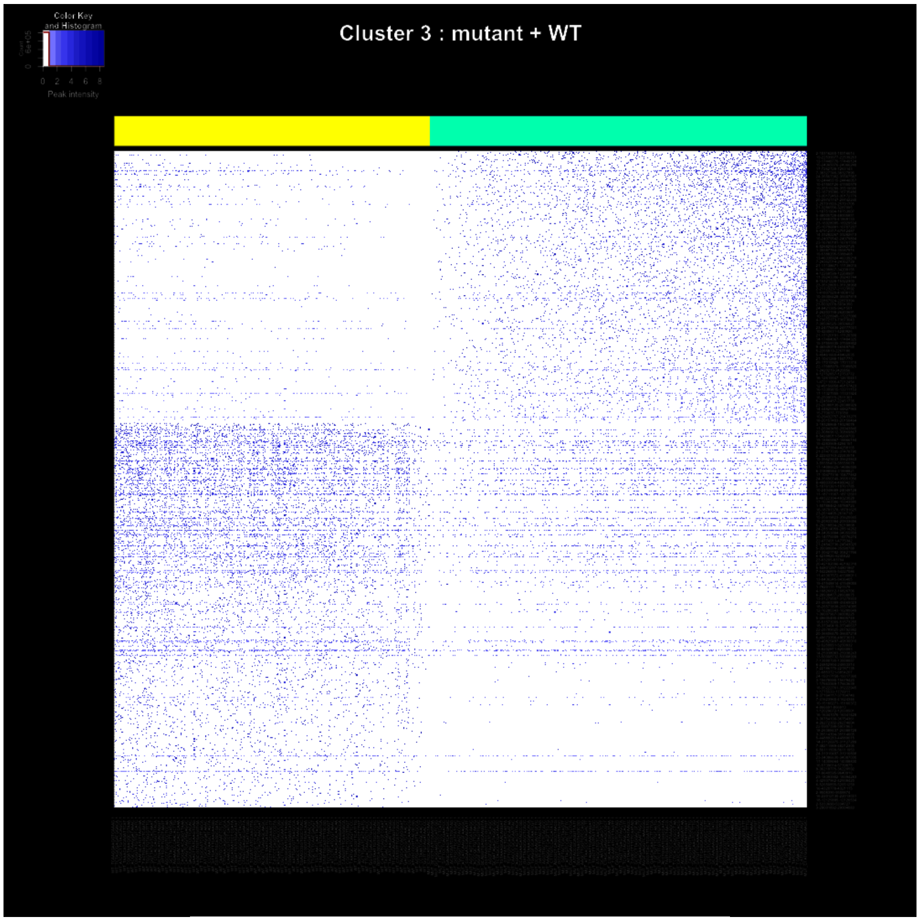
Color Key
and Histogram
Count
0 2 4 6 8
Peak intensity
Cluster 3 : mutant + WT

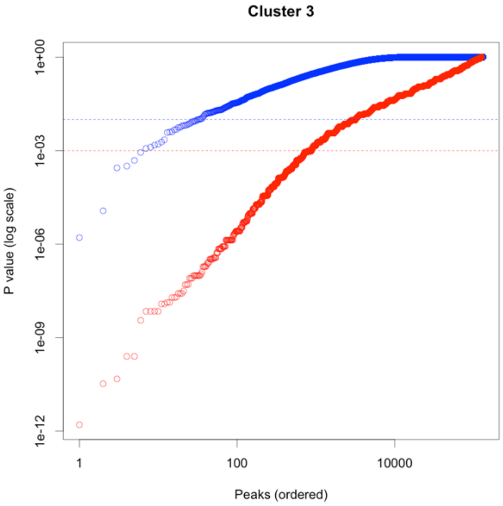
Cluster 3
P value (log scale)
1e+00
1e-03
1e-06
1e-09
1e-12
1
100
10000
Peaks (ordered)

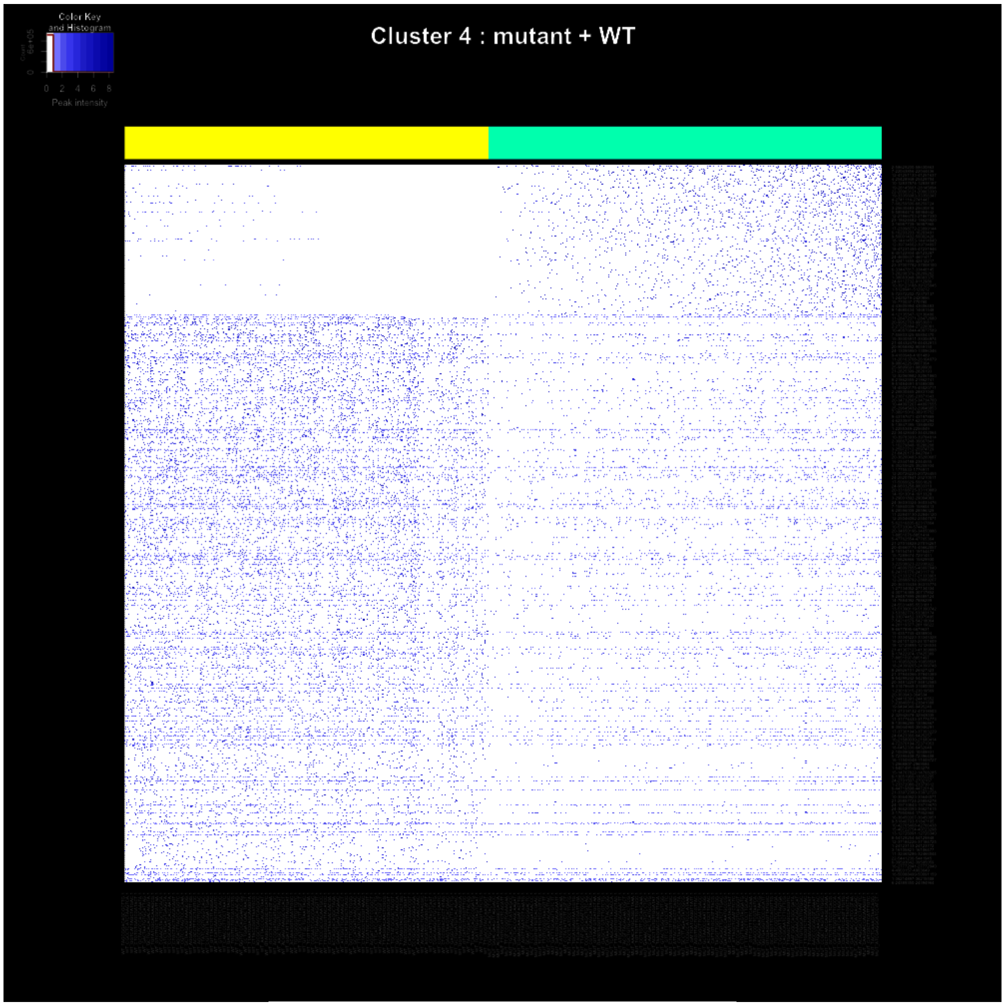
Color Key
and Histogram
Peak intensity
Cluster 4 : mutant + WT

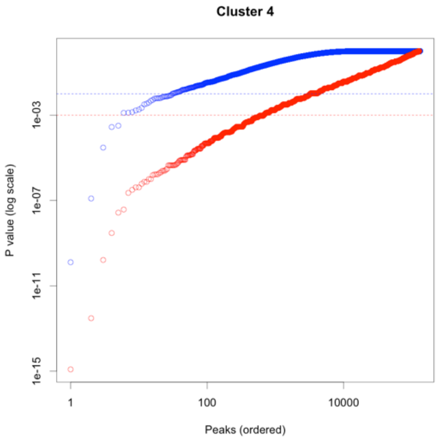
Cluster 4
P value (log scale)
1e-03
1e-07
1e-11
1e-15
1
100
10000
Peaks (ordered)

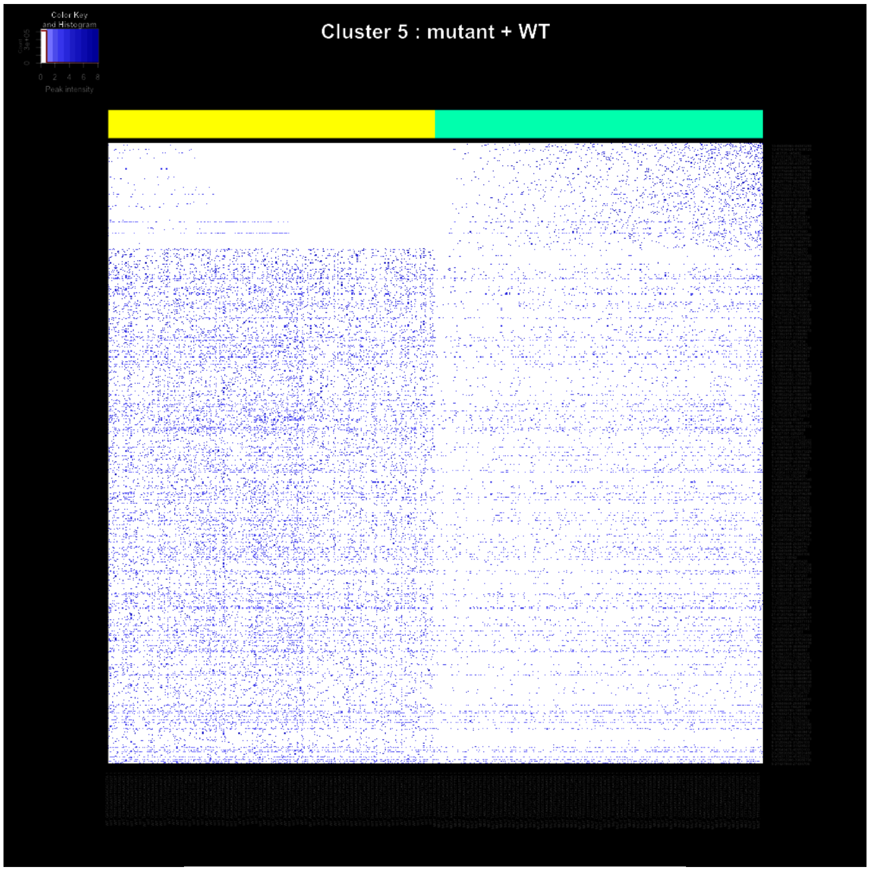
Color Key
and Histogram
Peak intensity
Cluster 5 : mutant + WT

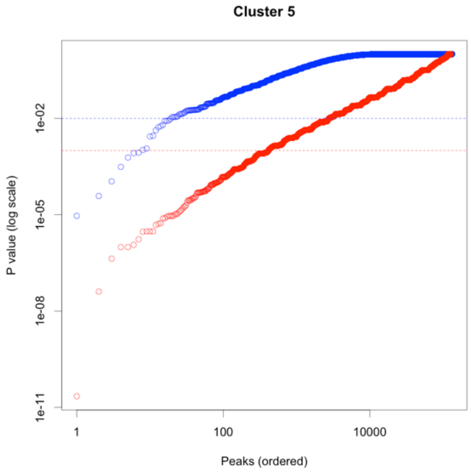
Cluster 5
P value (log scale)
Peaks (ordered)
1e-02
1e-05
1e-08
1e-11
1
100
10000

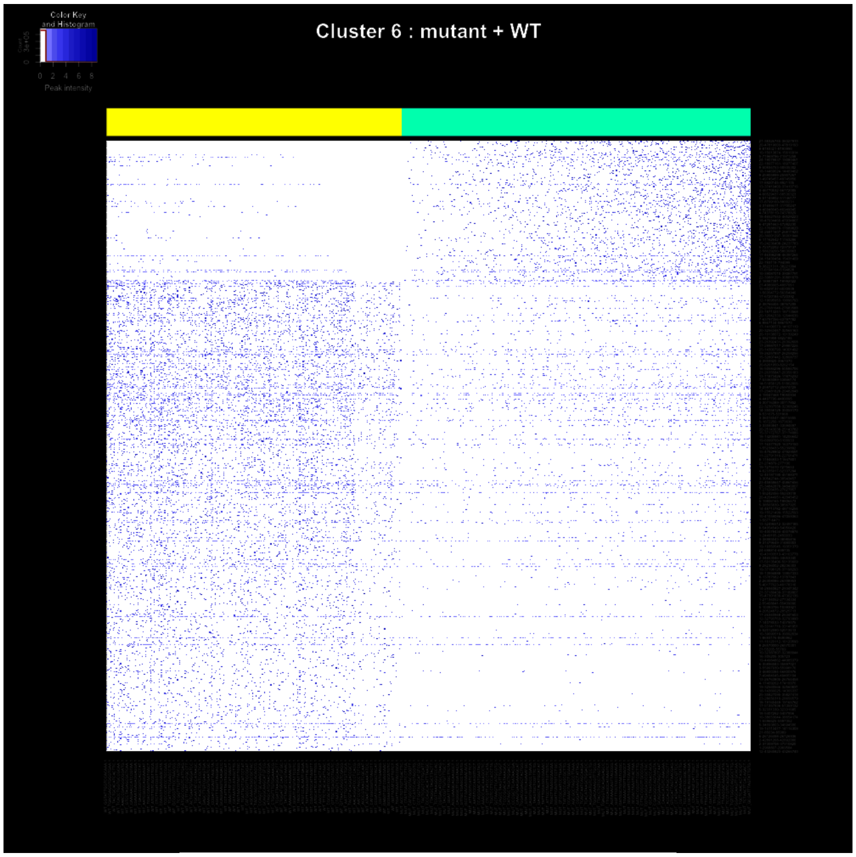
Cluster 6 : mutant + WT
Color Key
and Histogram
Peak intensity

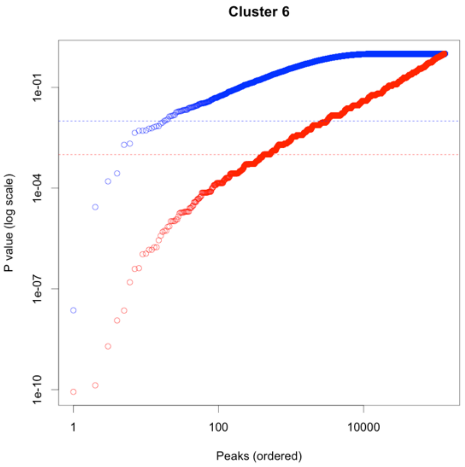
Cluster 6
P value (log scale)
1e-01
1e-04
1e-07
1e-10
1
100
10000
Peaks (ordered)

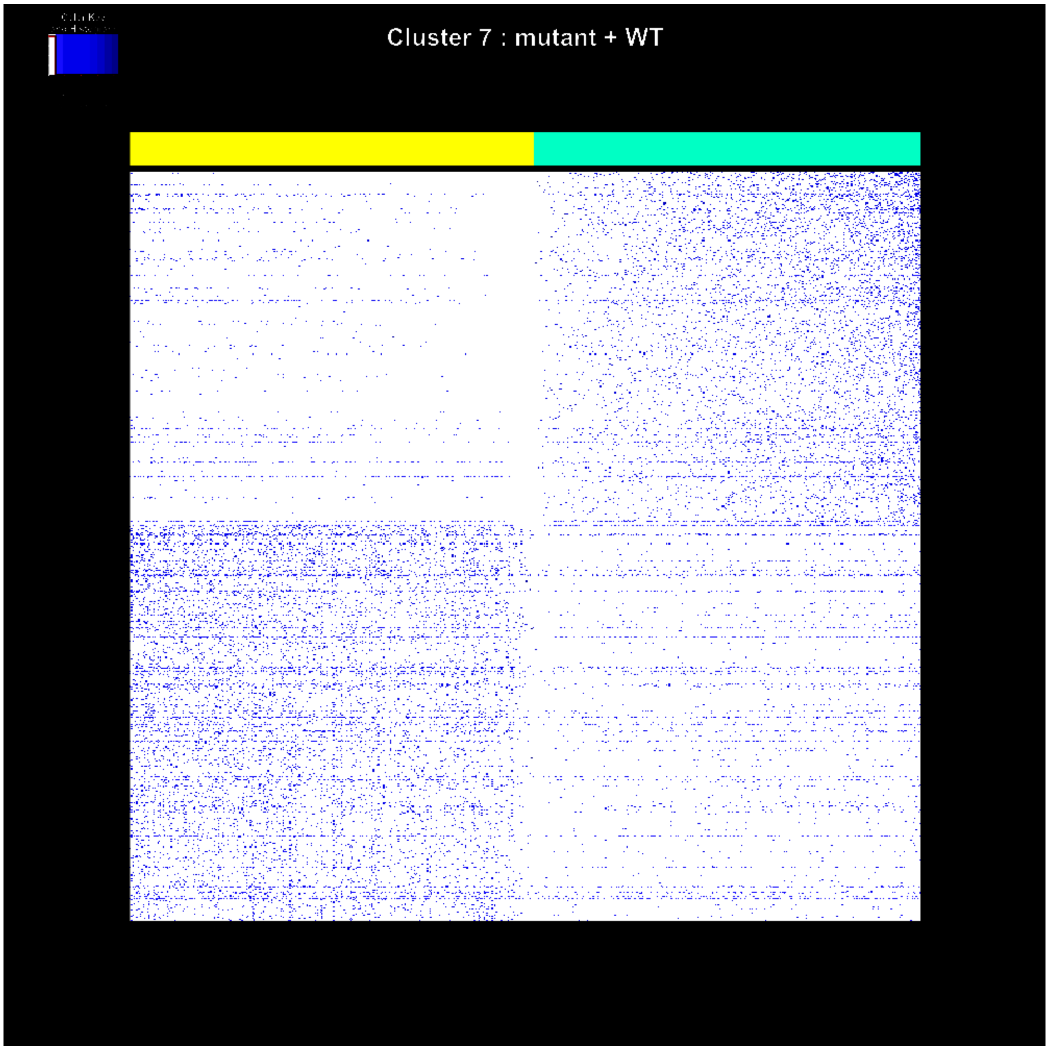
Cluster 7 : mutant + WT

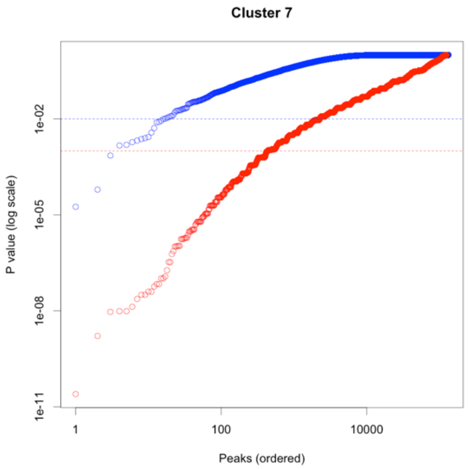
Cluster 7
P value (log scale)
1e-02
1e-05
1e-08
1e-11
1
100
10000
Peaks (ordered)

Cluster 8

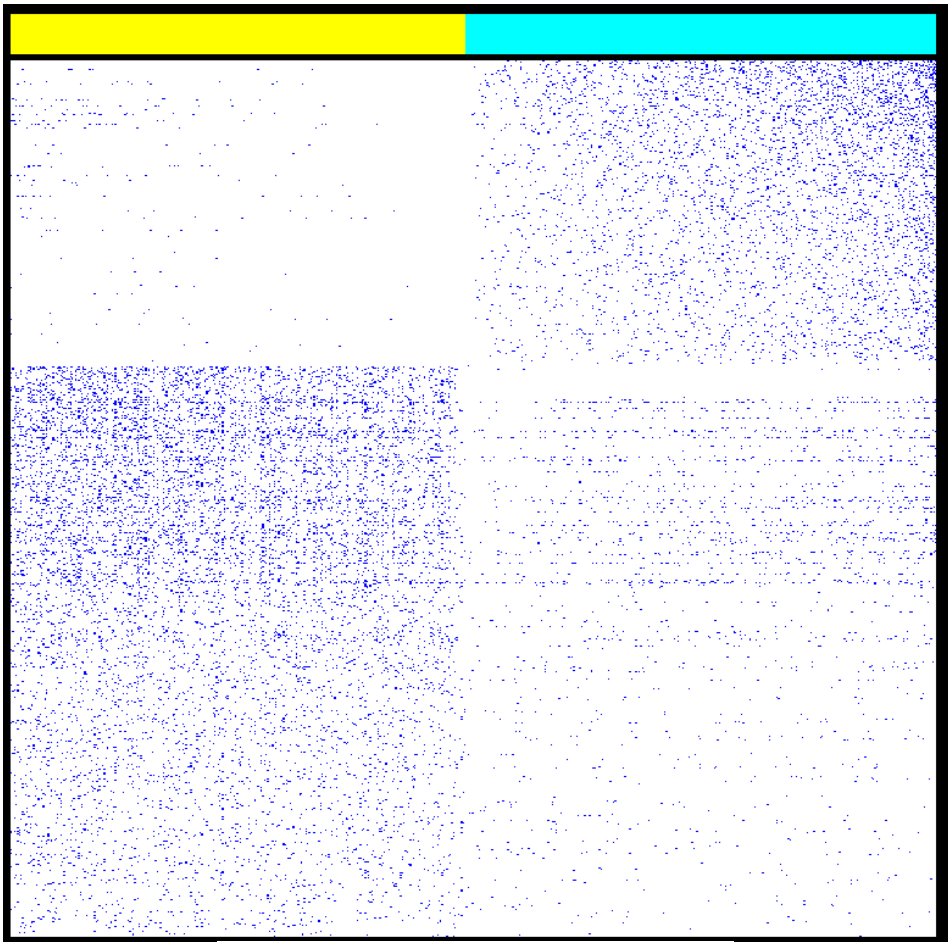

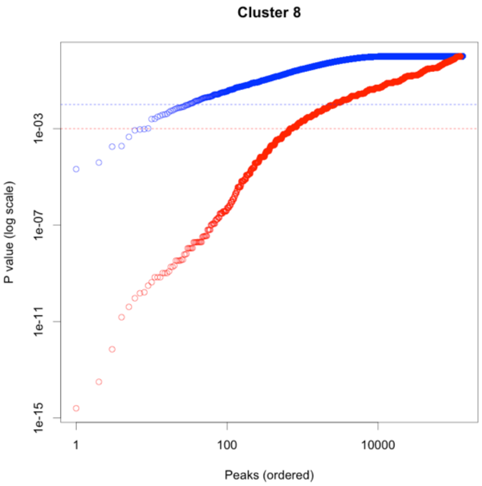
Cluster 8
P value (log scale)
1e-03
1e-07
1e-11
1e-15
1
100
10000
Peaks (ordered)

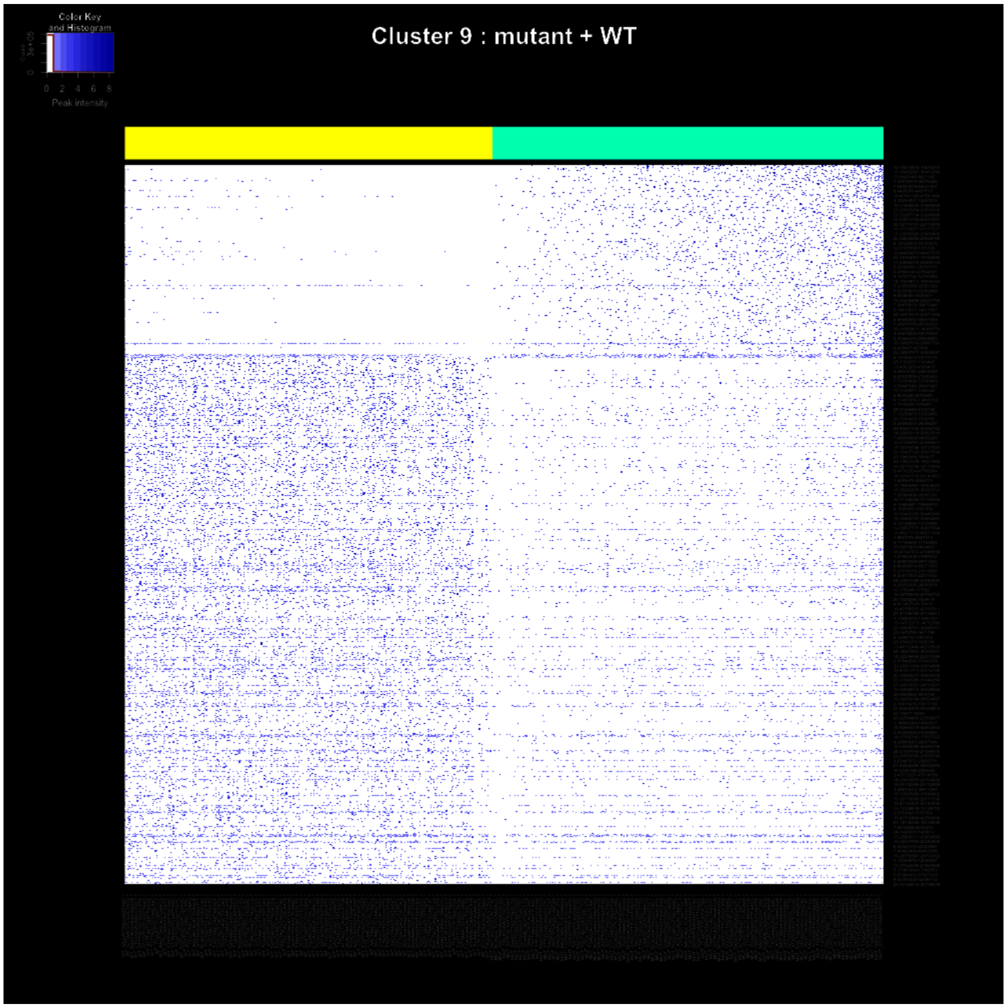
Color Key
and Histogram
Peak intensity
0 2 4 6 8
Cluster 9 : mutant + WT

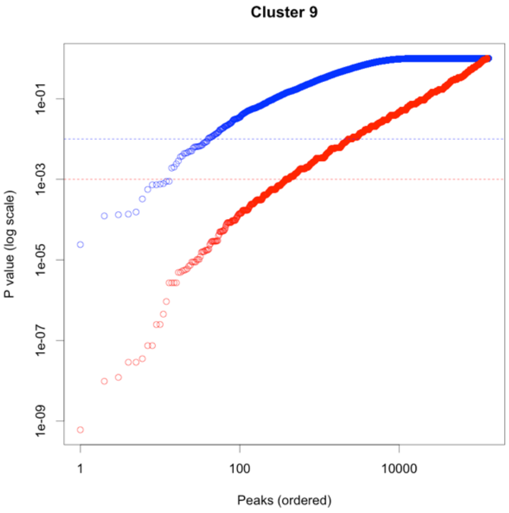
Cluster 9
P value (log scale)
1e-01
1e-03
1e-05
1e-07
1e-09
1
100
10000
Peaks (ordered)

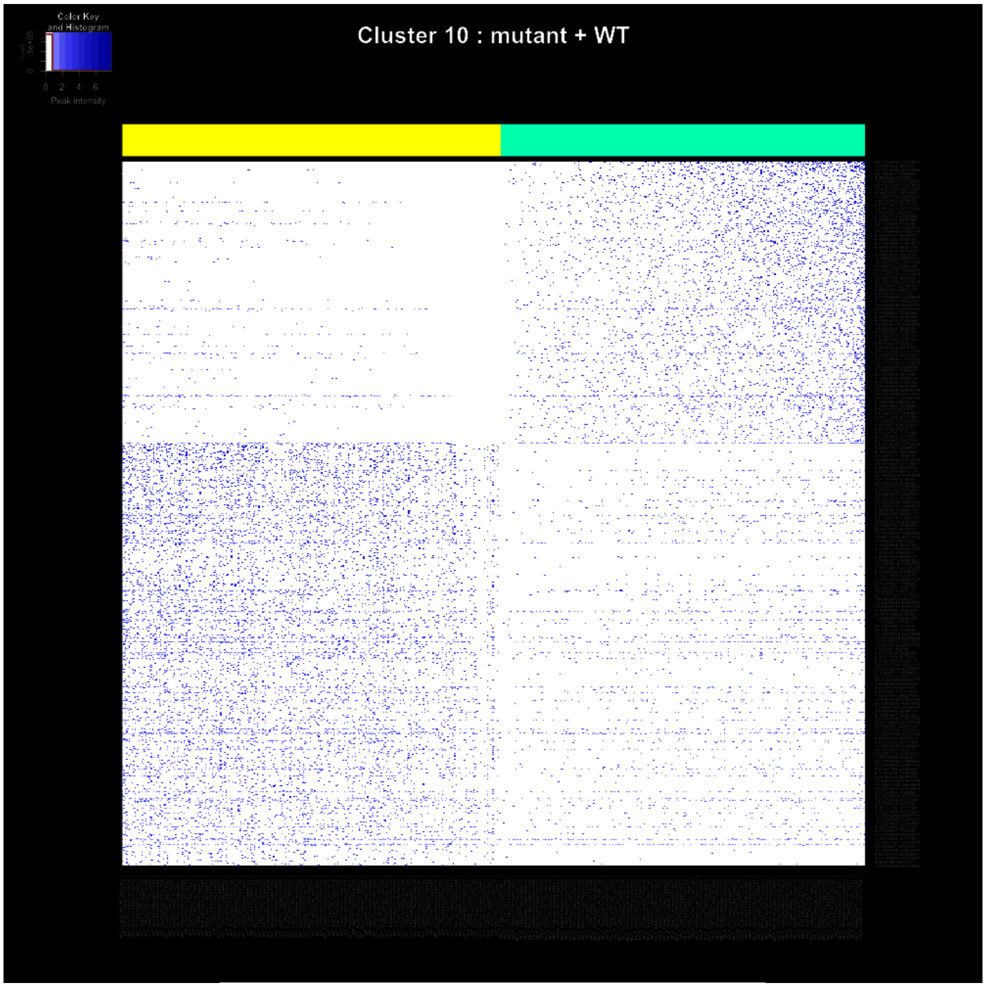
Color Key and Histogram
0 2 4 6
Peak intensity
Cluster 10 : mutant + WT

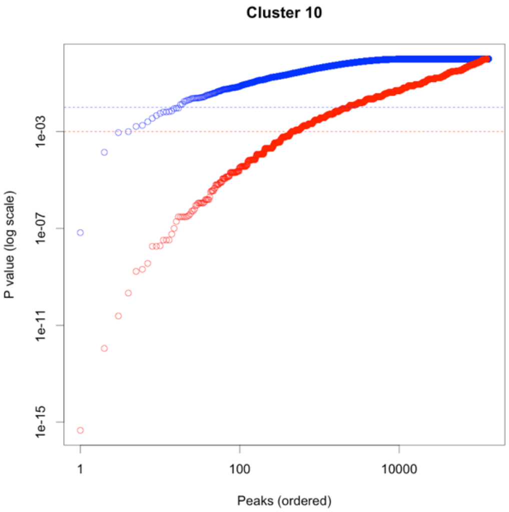
Cluster 10
P value (log scale)
1e-03
1e-07
1e-11
1e-15
1
100
10000
Peaks (ordered)

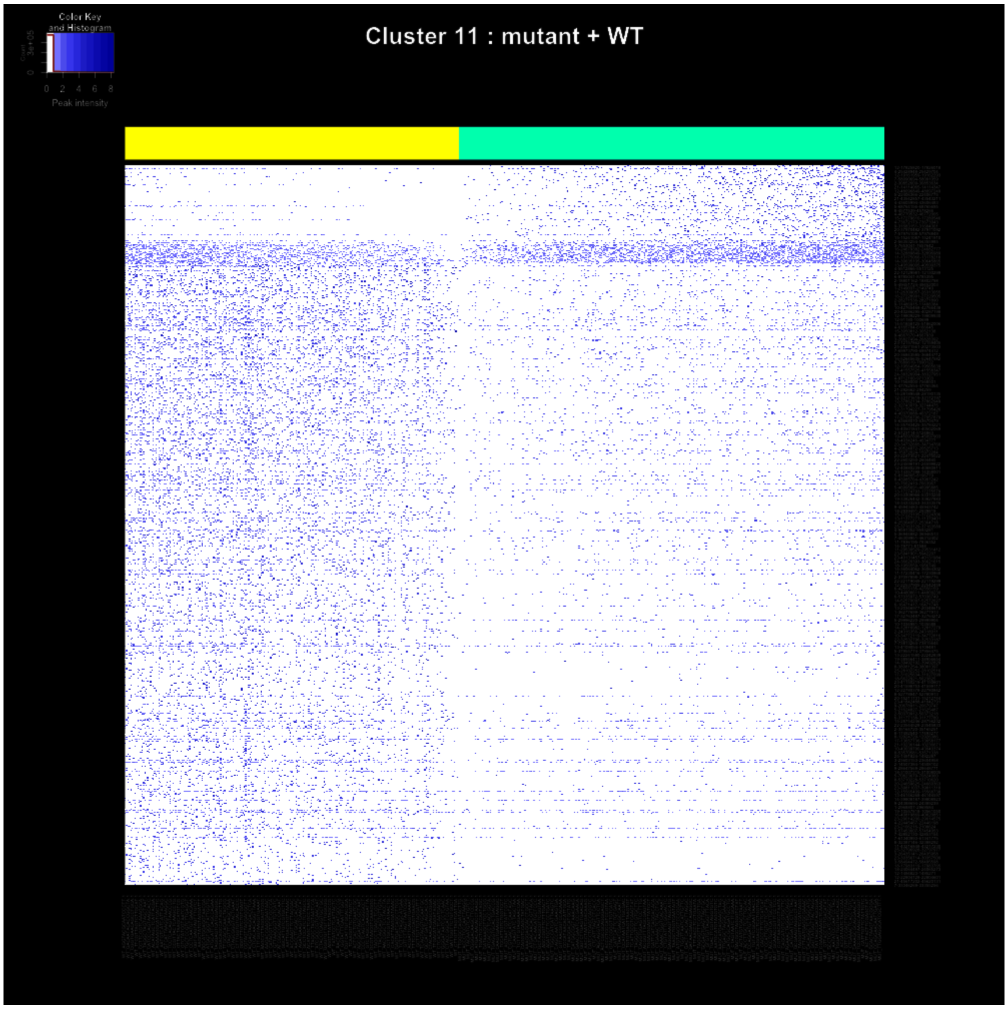
Color Key
and Histogram
0 2 4 6 8
Peak intensity
Cluster 11 : mutant + WT

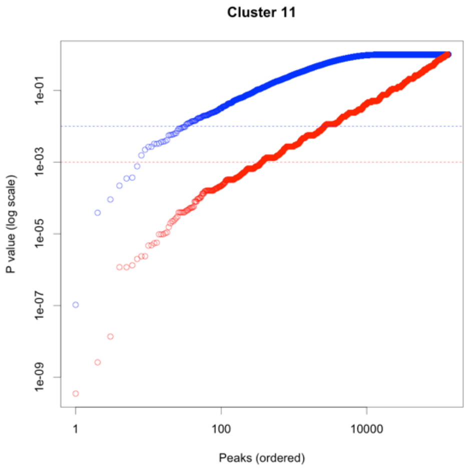
Cluster 11
P value (log scale)
1e-01
1e-03
1e-05
1e-07
1e-09
1
100
10000
Peaks (ordered)

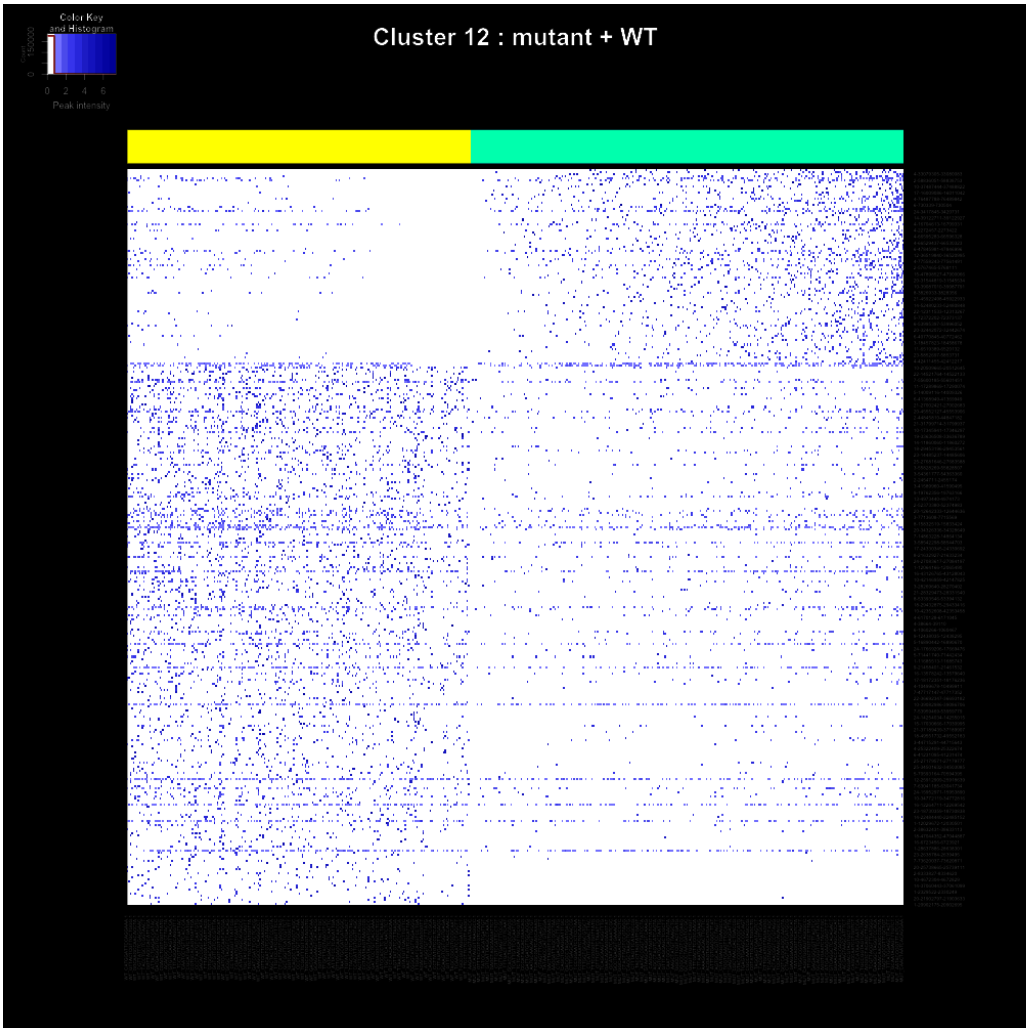
Cluster 12 : mutant + WT
Color Key and Histogram
Count
0
150000
0 2 4 6
Peak intensity

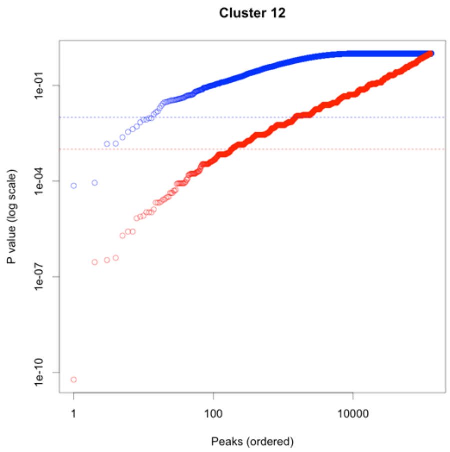
Cluster 12
P value (log scale)
1e-01
1e-04
1e-07
1e-10
1
100
10000
Peaks (ordered)

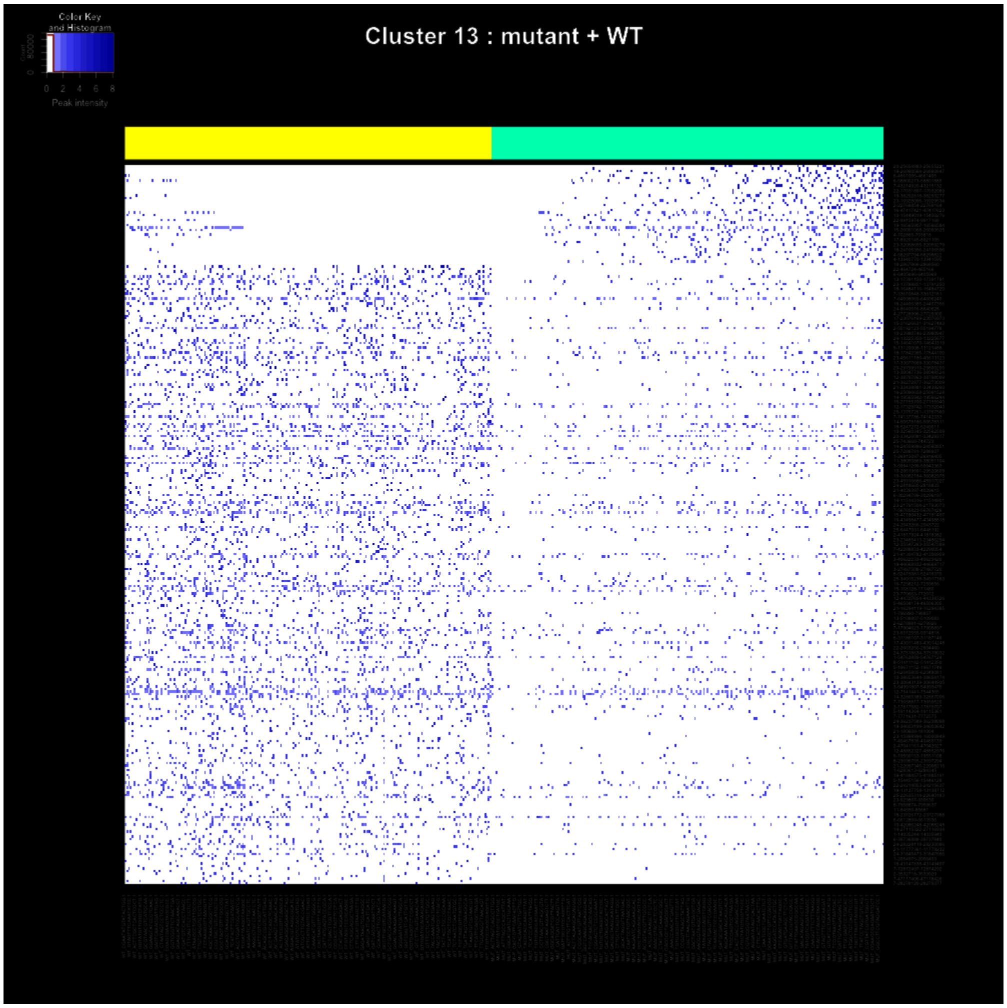
Color Key
and Histogram
Count
Peak intensity
Cluster 13 : mutant + WT

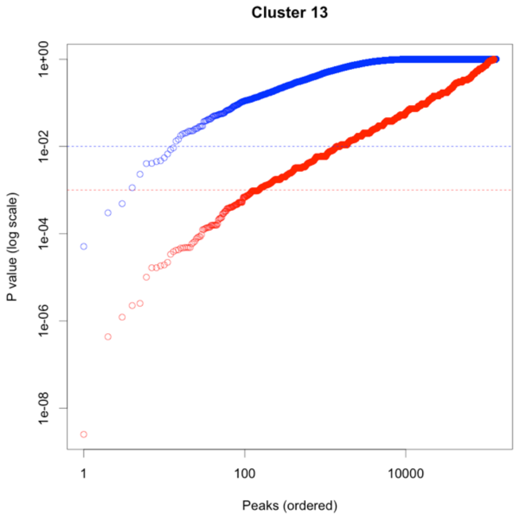
Cluster 13
P value (log scale)
1e+00
1e-02
1e-04
1e-06
1e-08
1
100
10000
Peaks (ordered)

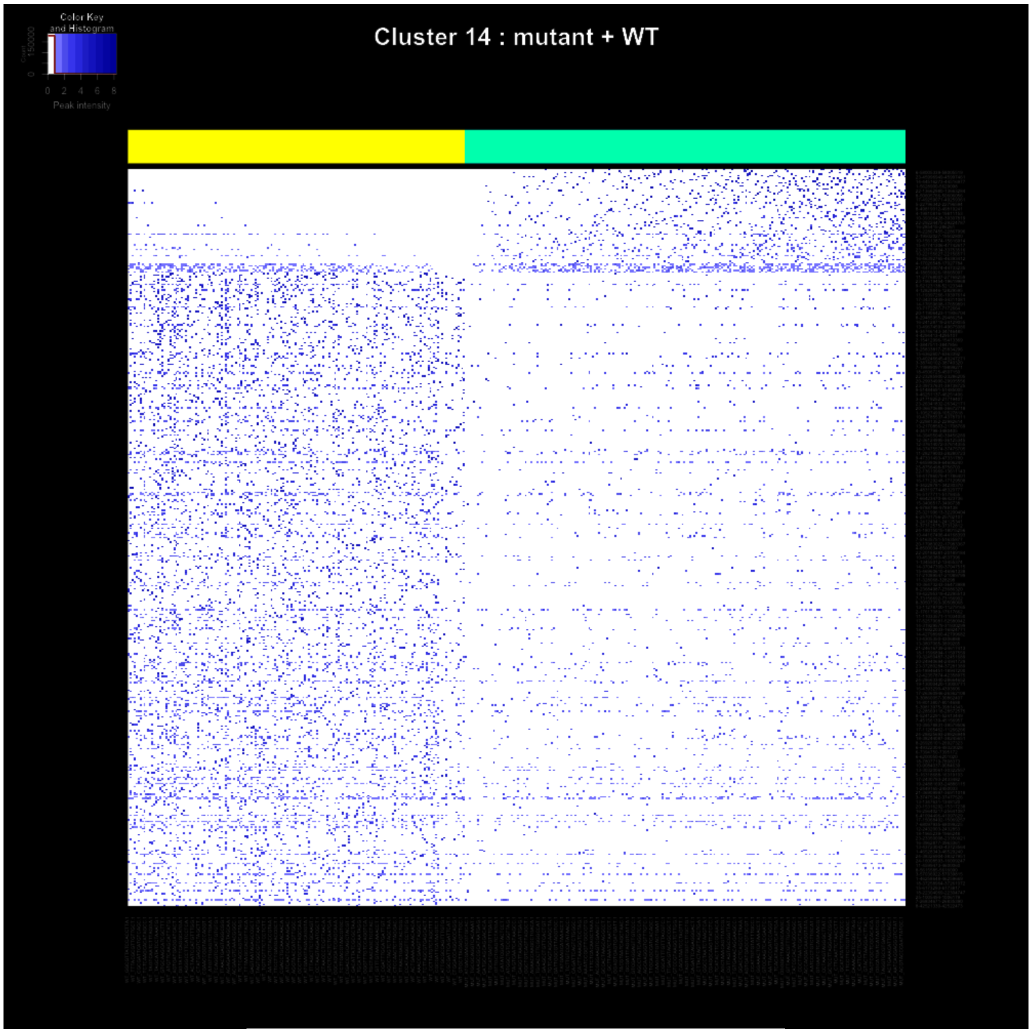
Color Key and Histogram
Count
0
150000
0 2 4 6 8
Peak intensity
Cluster 14 : mutant + WT

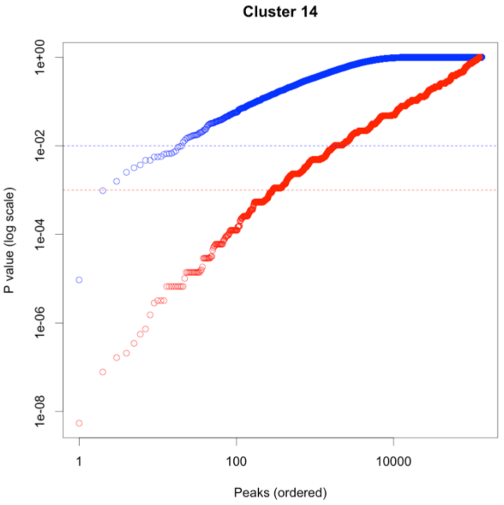
Cluster 14
P value (log scale)
1e+00
1e-02
1e-04
1e-06
1e-08
1
100
10000
Peaks (ordered)

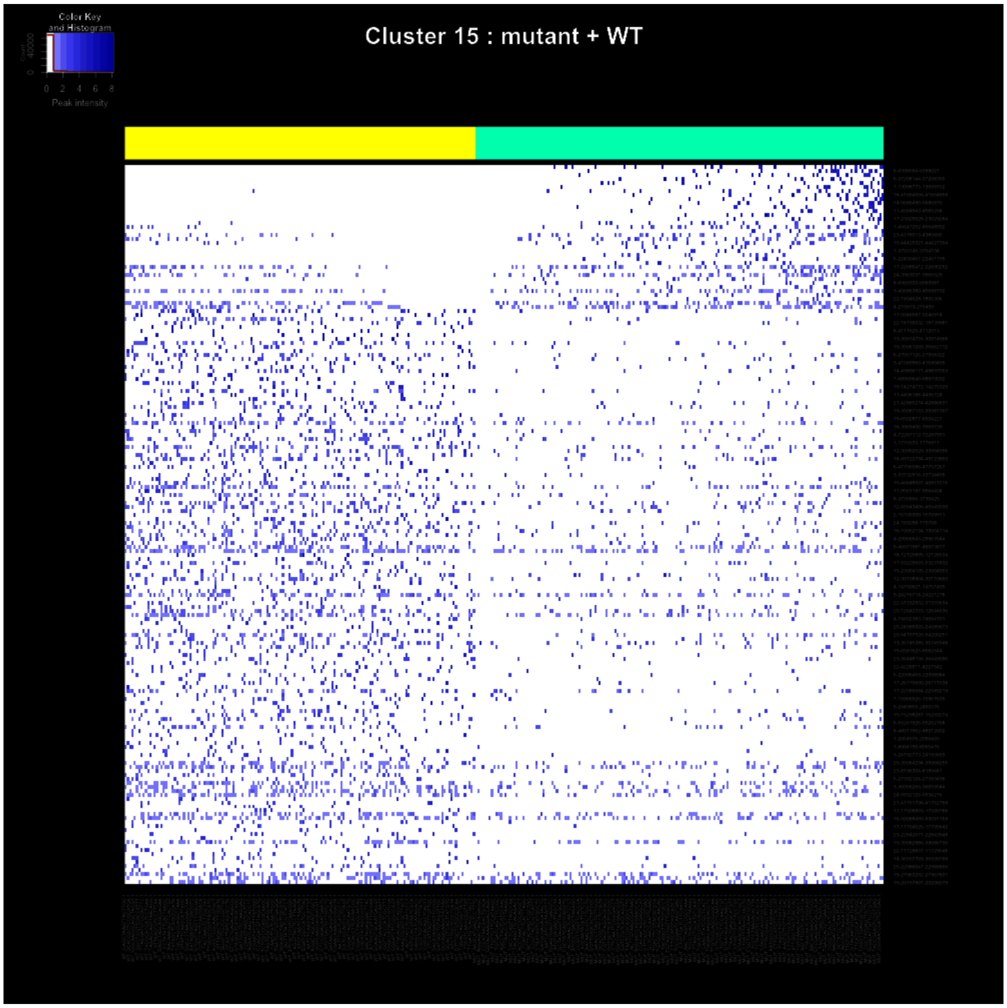

Color Key
and Histogram
Count
0
40000
0 2 4 6 8
Peak intensity
Cluster 15 : mutant + WT


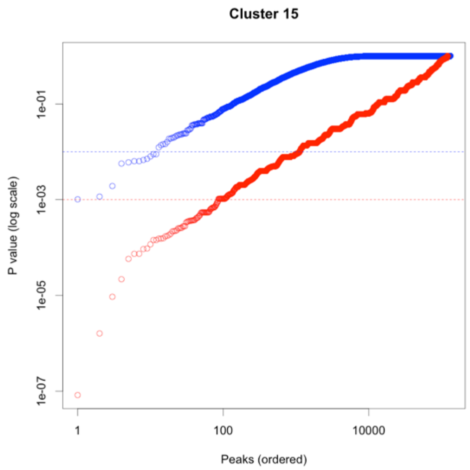

Cluster 15
P value (log scale)
1e-01
1e-03
1e-05
1e-07
1
100
10000
Peaks (ordered)

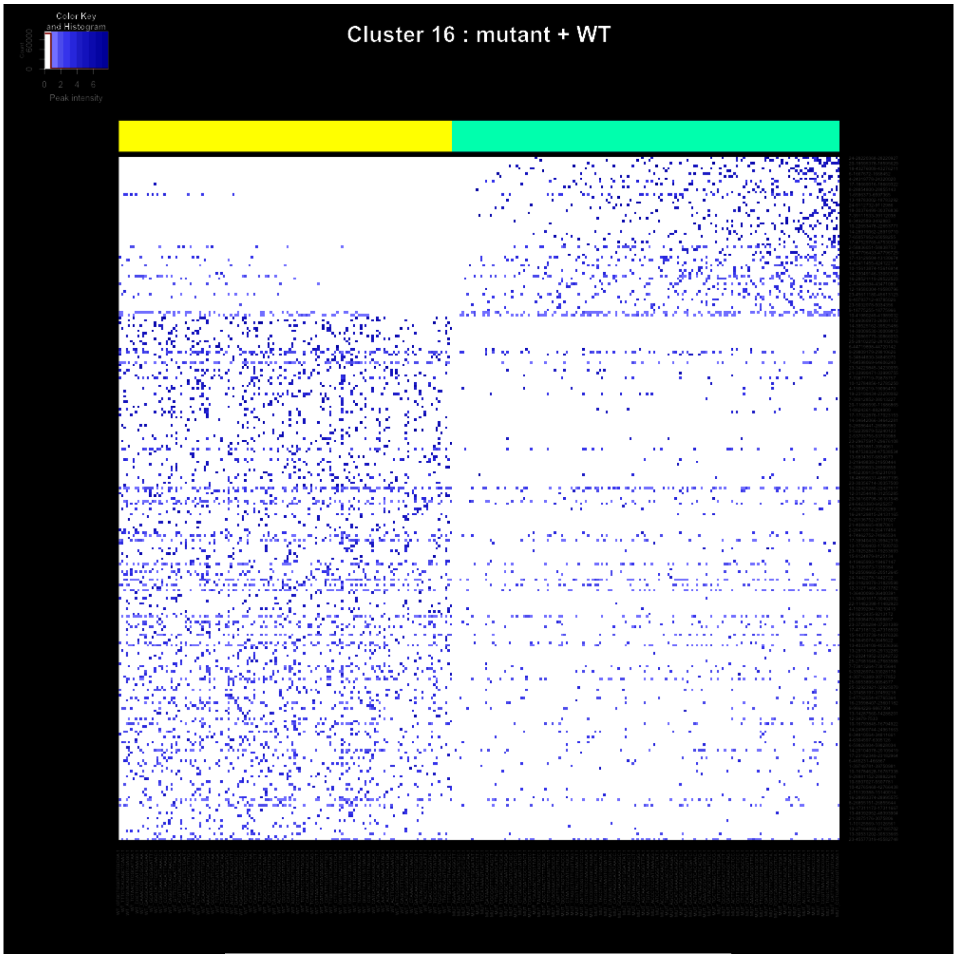
Color Key and Histogram
Count
0 2 4 6
Peak intensity
Cluster 16 : mutant + WT

**Cluster 16**

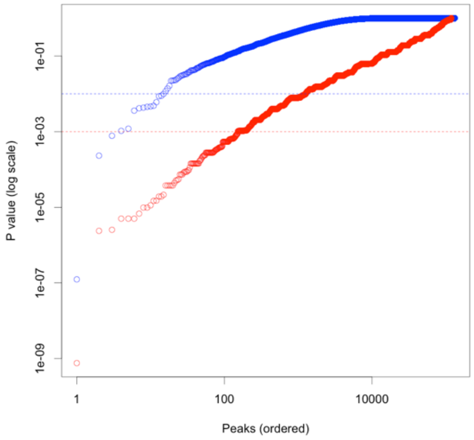
P value (log scale)
1e-01
1e-03
1e-05
1e-07
1e-09
1
100
10000
Peaks (ordered)

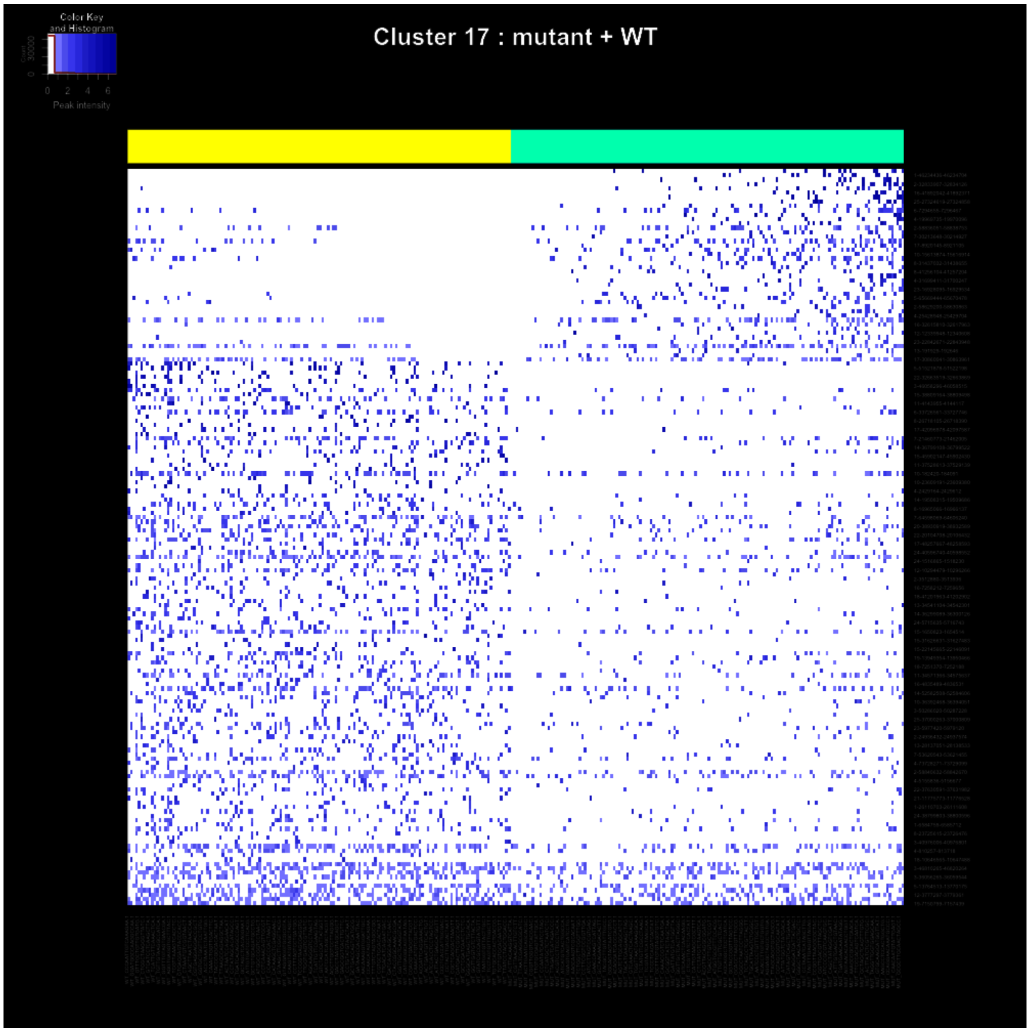
Color Key
and Histogram
Count
0
30000
0
2
4
6
Peak intensity
Cluster 17 : mutant + WT

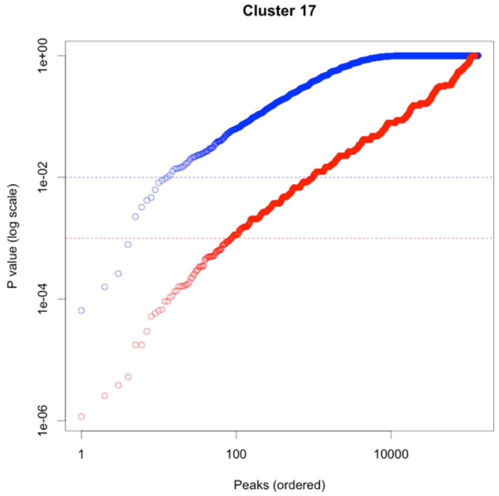
Cluster 17
P value (log scale)
1e+00
1e-02
1e-04
1e-06
1
100
10000
Peaks (ordered)

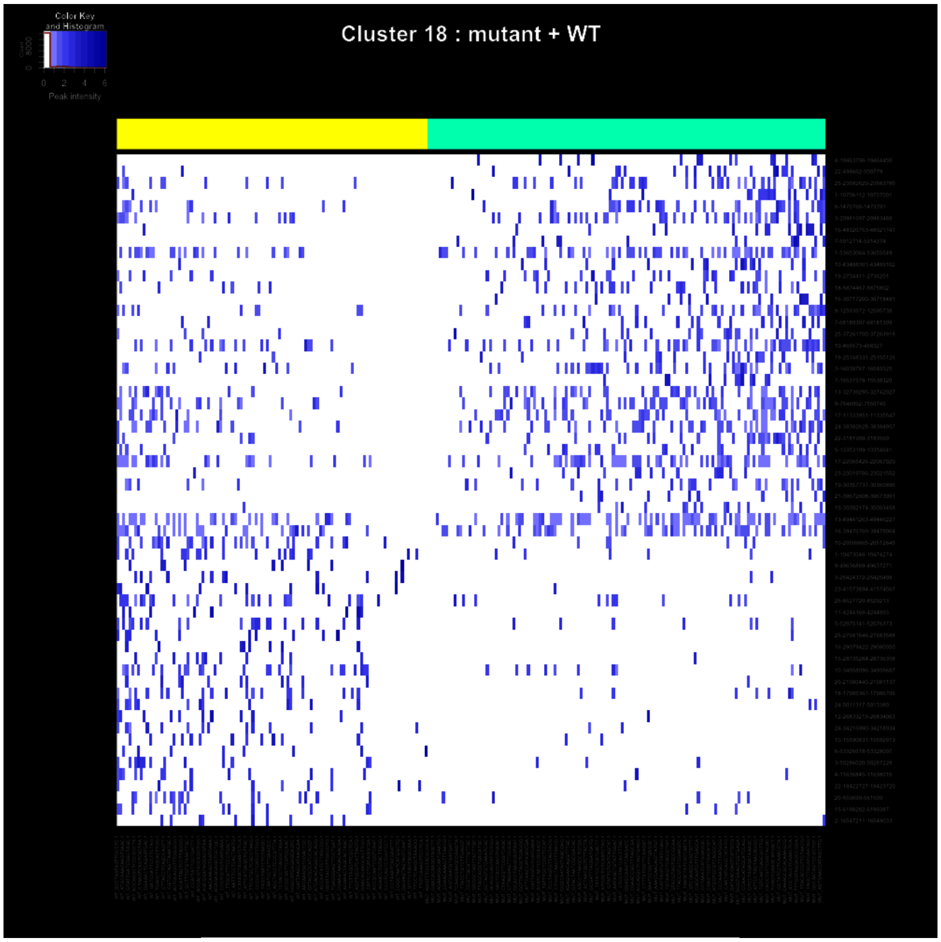

Color Key and Histogram
Peak intensity
Cluster 18 : mutant + WT


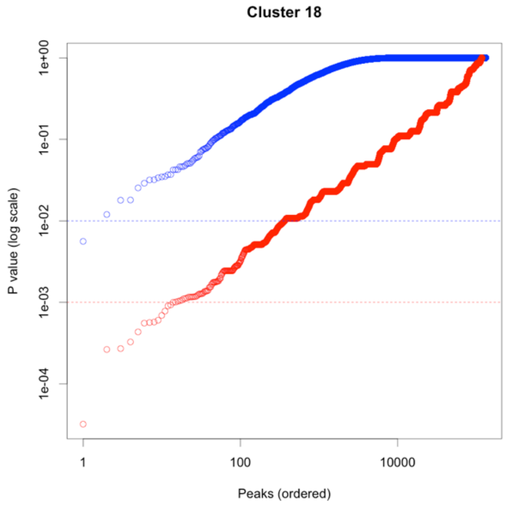

Cluster 18
P value (log scale)
1e+00
1e-01
1e-02
1e-03
1e-04
1
100
10000
Peaks (ordered)

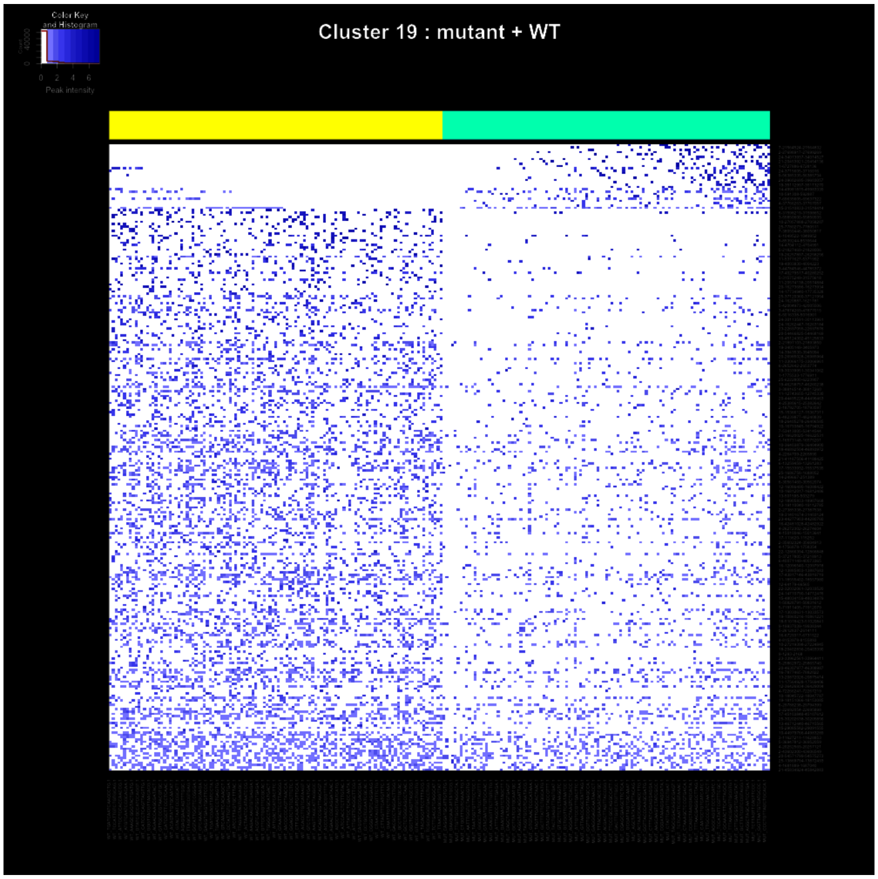
Cluster 19 : mutant + WT
Color Key
and Histogram
0 2 4 6
Peak intensity

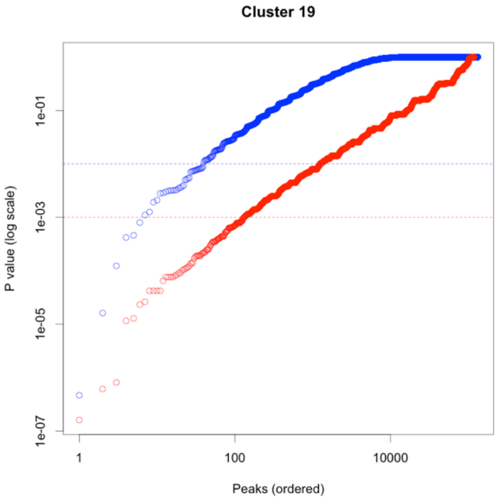
Cluster 19
P value (log scale)
1e-01
1e-03
1e-05
1e-07
1
100
10000
Peaks (ordered)

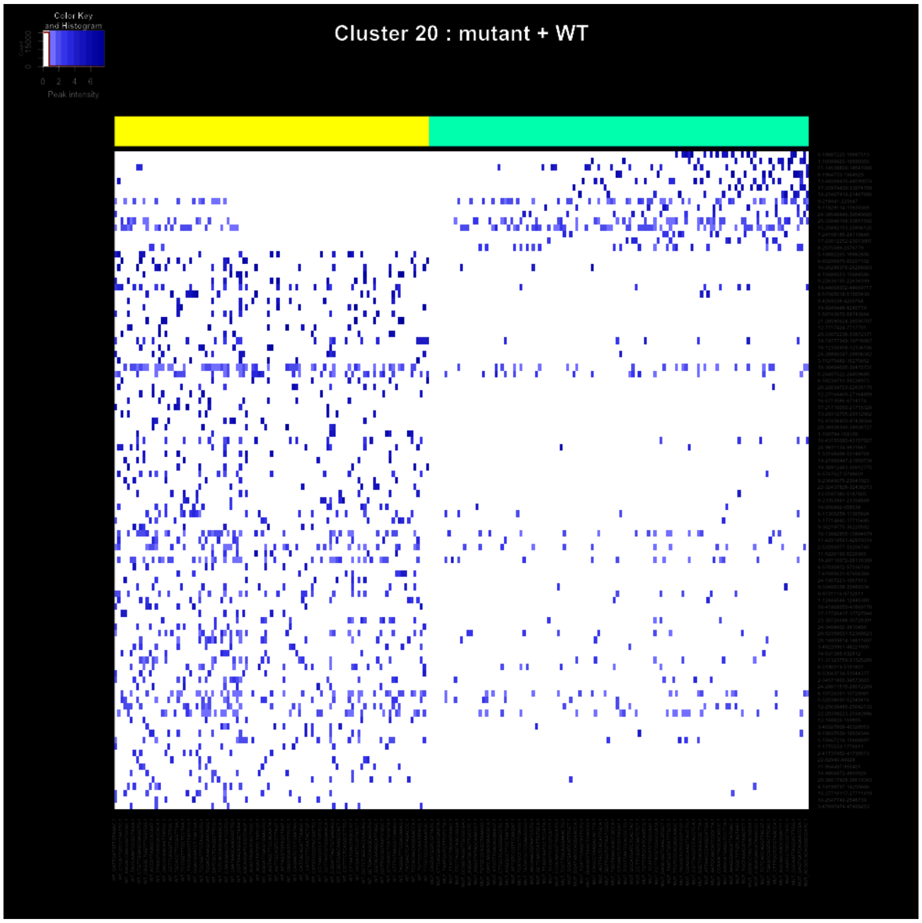
Color Key
and Histogram
Count
0
2
4
6
Peak intensity
Cluster 20 : mutant + WT

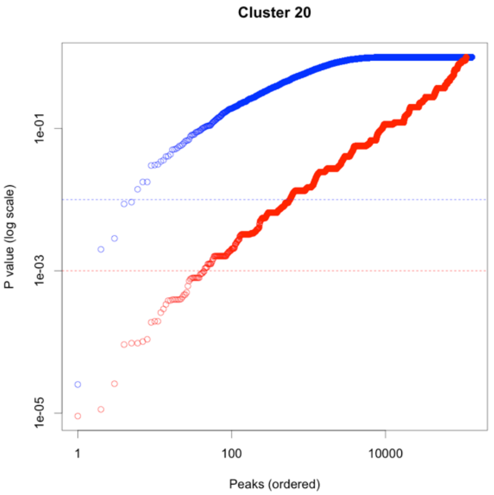
Cluster 20
P value (log scale)
1e-01
1e-03
1e-05
1
100
10000
Peaks (ordered)

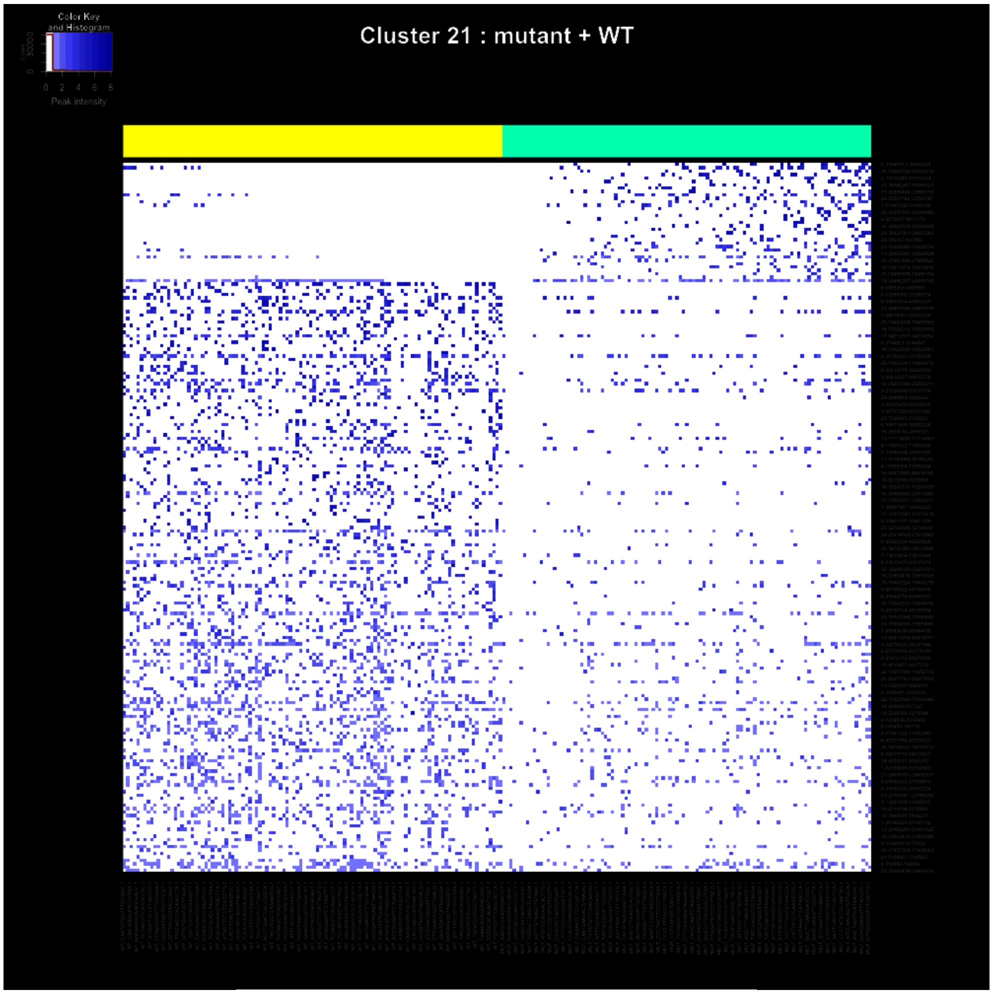
Color Key
and Histogram
0 2 4 6 8
Peak intensity
Cluster 21 : mutant + WT

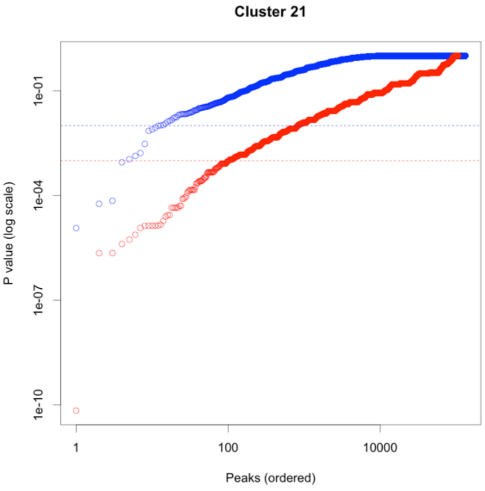
Cluster 21
P value (log scale)
1e-01
1e-04
1e-07
1e-10
1
100
10000
Peaks (ordered)

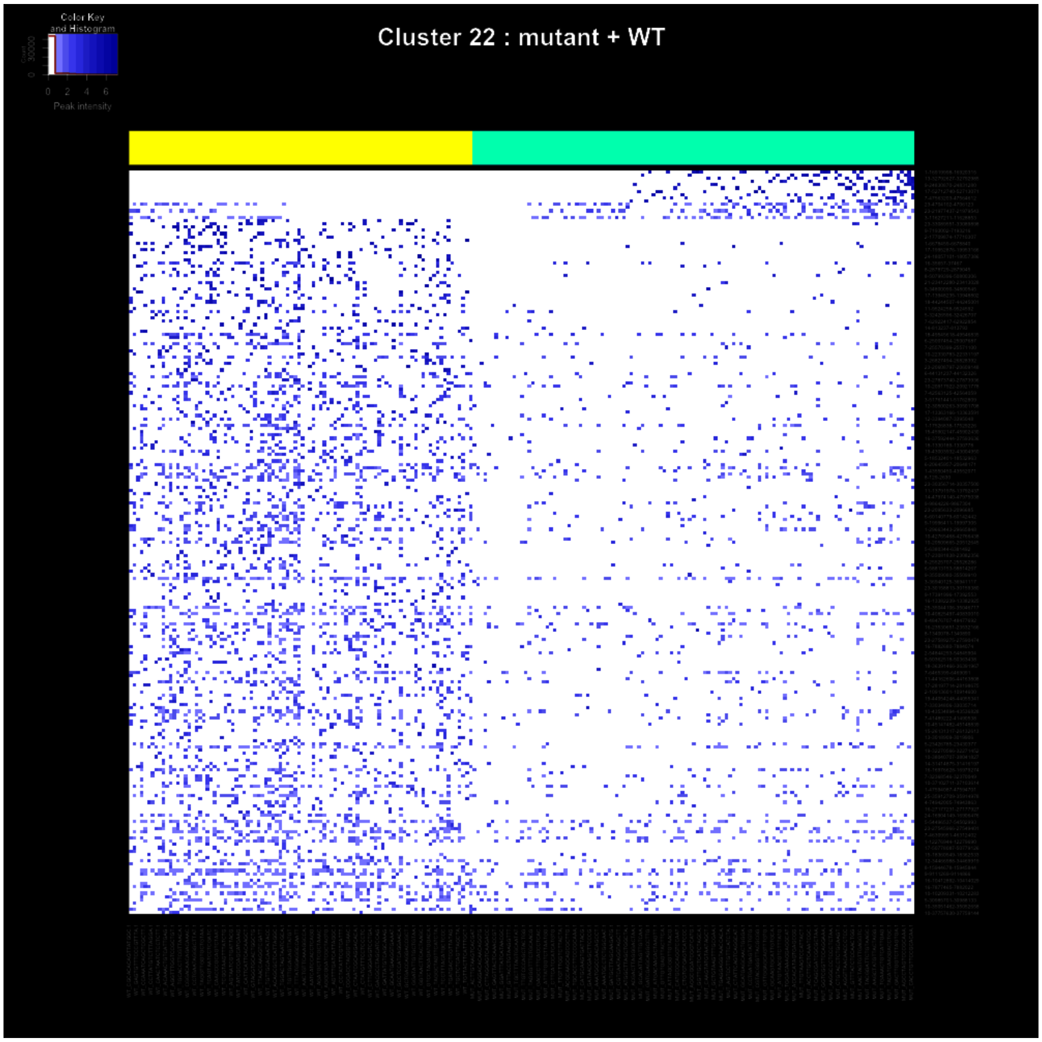
Color Key
and Histogram
Peak intensity
Cluster 22 : mutant + WT

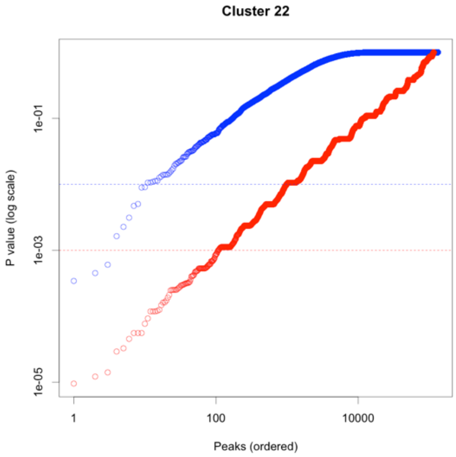
Cluster 22
P value (log scale)
1e-01
1e-03
1e-05
1
100
10000
Peaks (ordered)

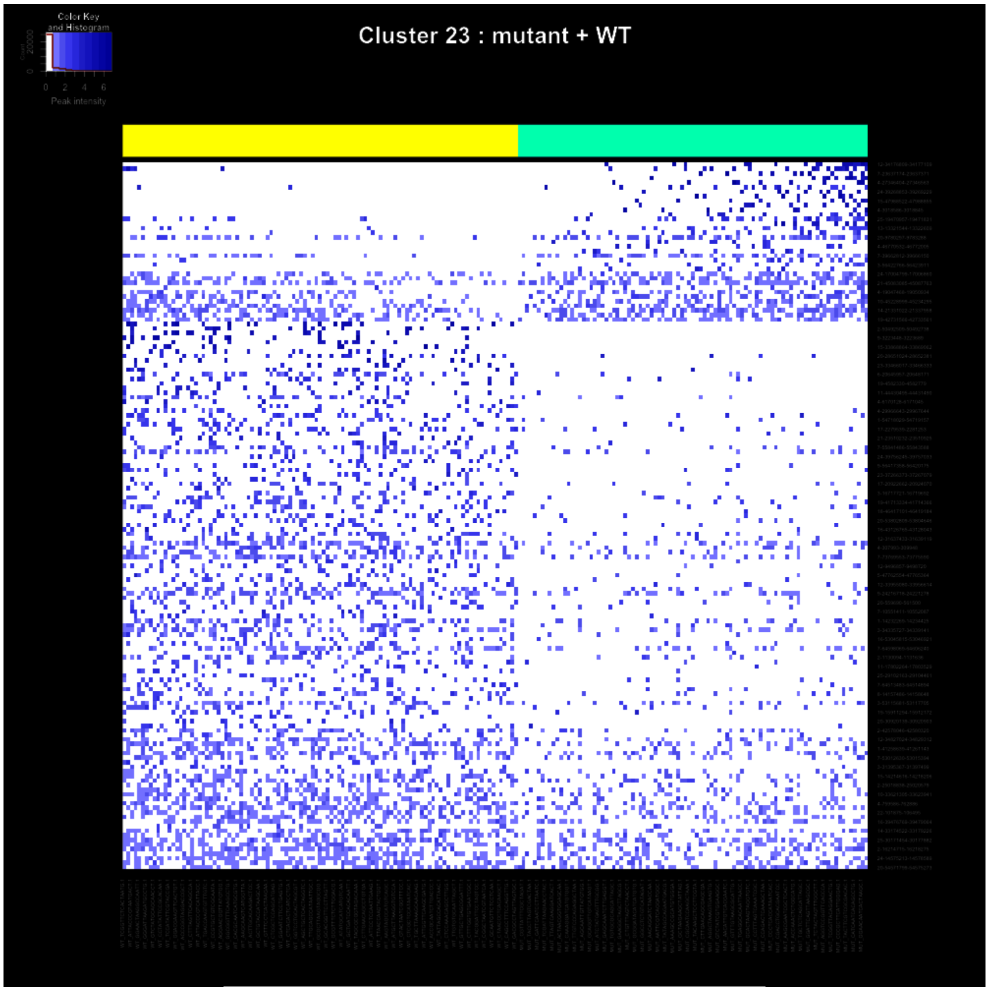
Color Key and Histogram
0 2 4 6
Peak intensity
Cluster 23 : mutant + WT

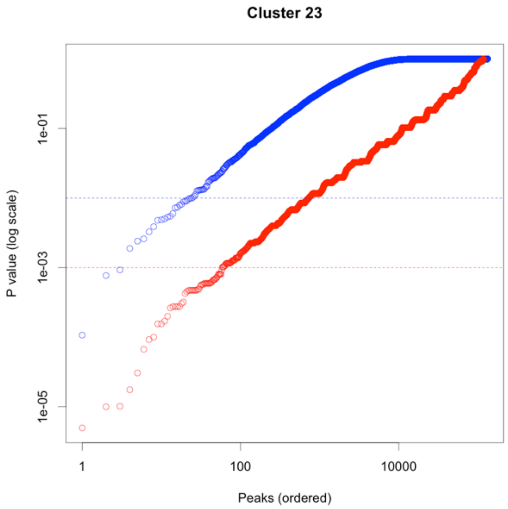
Cluster 23
P value (log scale)
1e-01
1e-03
1e-05
1
100
10000
Peaks (ordered)

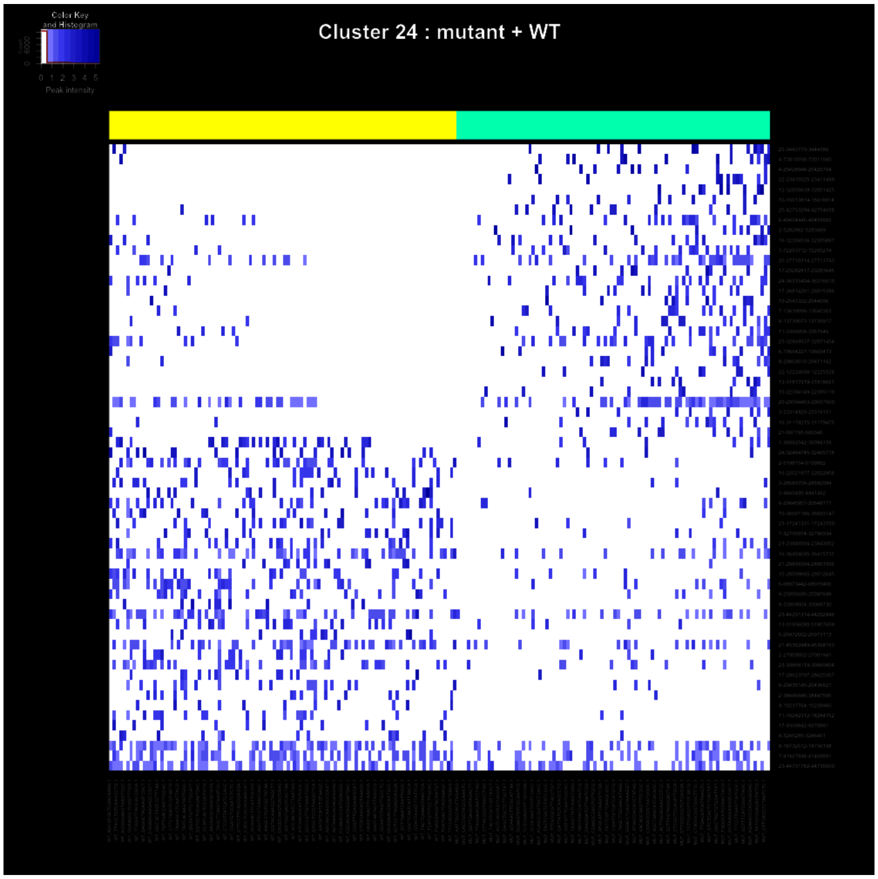
Color Key
and Histogram
0 1 2 3 4 5
Peak intensity
Cluster 24 : mutant + WT

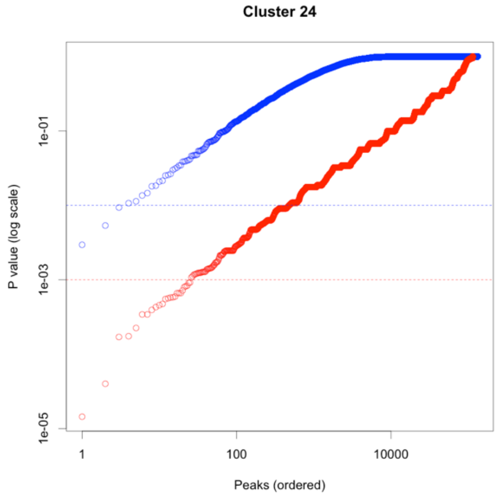
Cluster 24
1e-01
1e-03
1e-05
P value (log scale)
1
100
10000
Peaks (ordered)

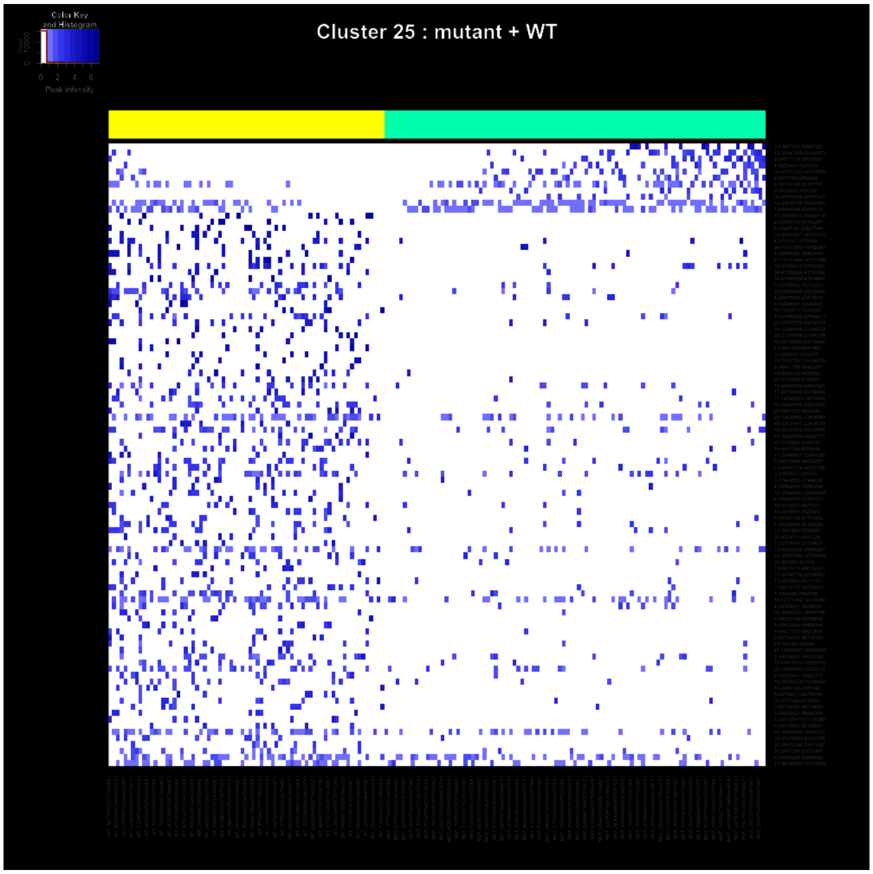
Color Key
and Histogram
0 2 4 6
Peak intensity
Cluster 25 : mutant + WT

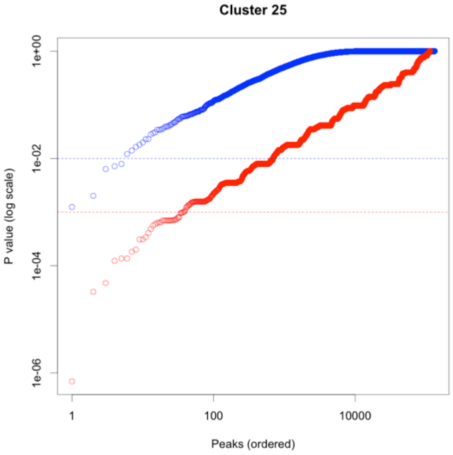
Cluster 25
1e+00
1e-02
1e-04
1e-06
P value (log scale)
1
100
10000
Peaks (ordered)

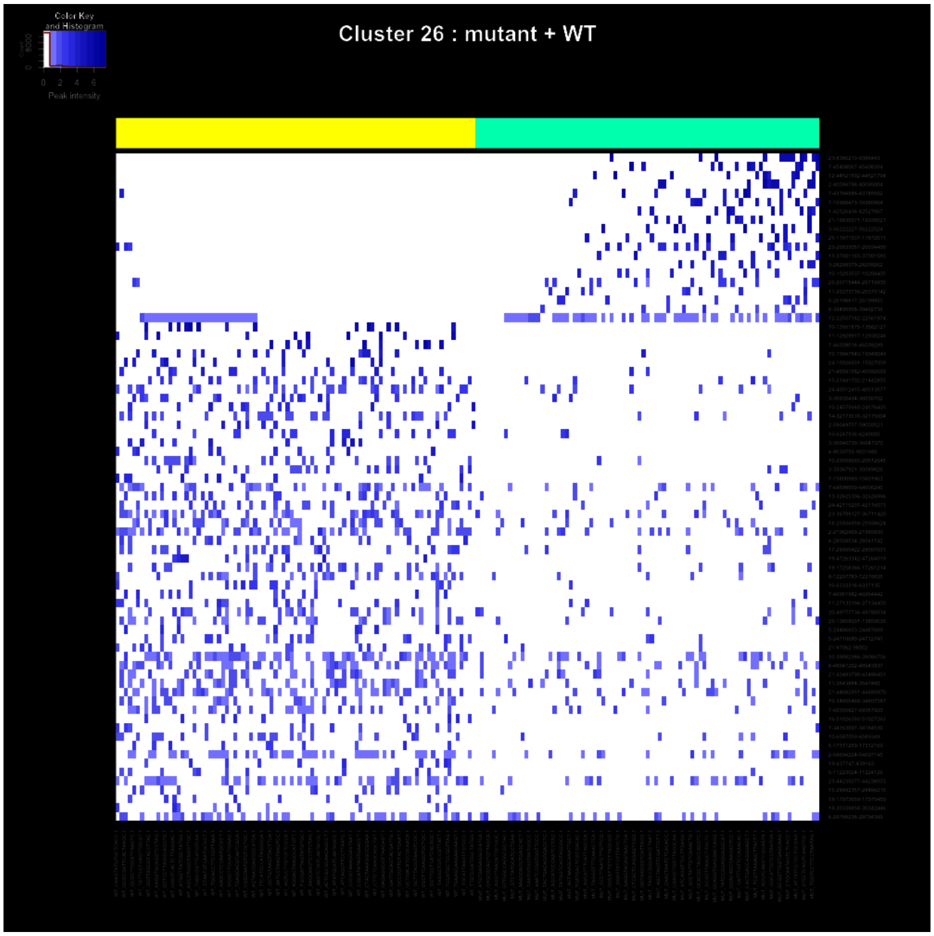
Color Key
and Histogram
0 2 4 6
Peak intensity
Cluster 26 : mutant + WT

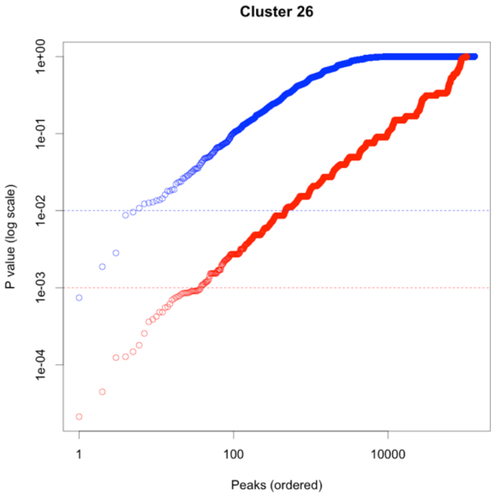
Cluster 26
P value (log scale)
1e+00
1e-01
1e-02
1e-03
1e-04
1
100
10000
Peaks (ordered)

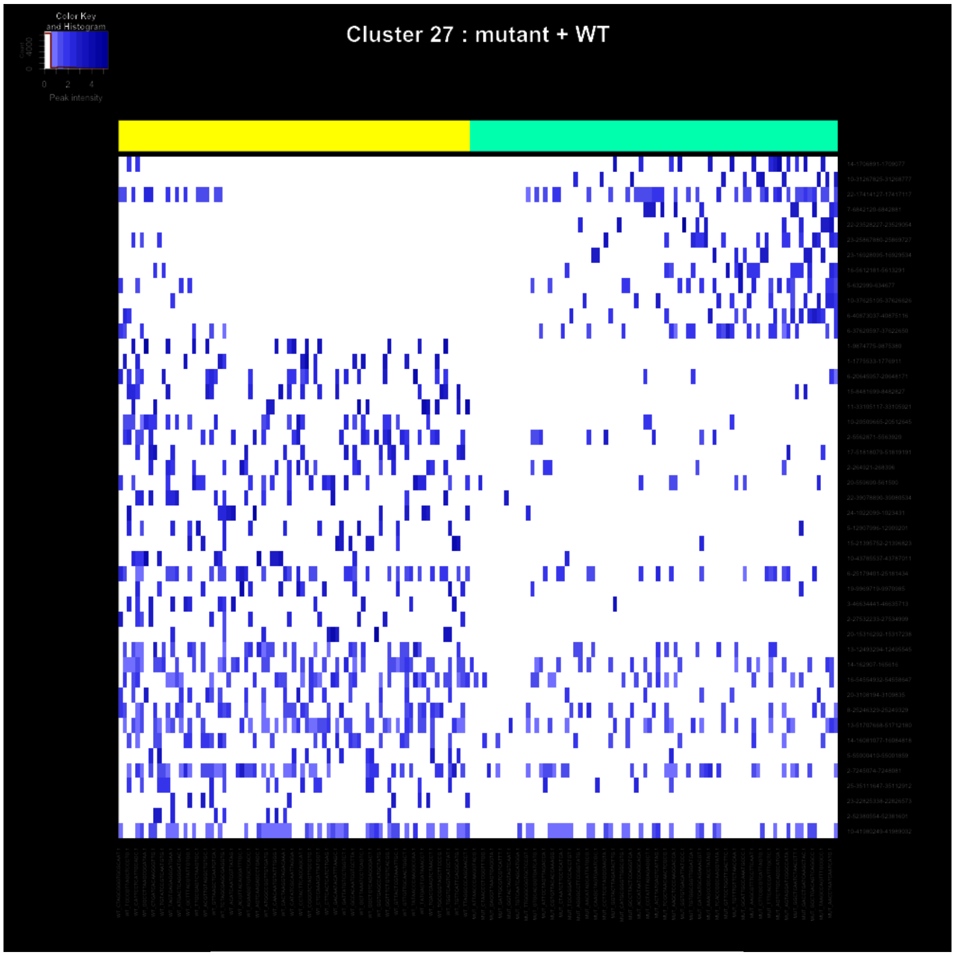
Cluster 27 : mutant + WT
Color Key and Histogram
0
2
4
Peak intensity

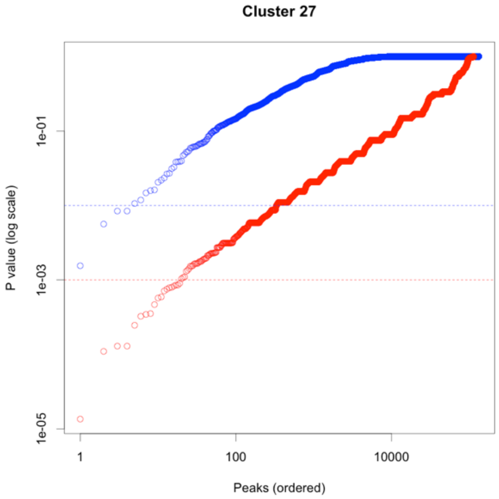
Cluster 27
P value (log scale)
1e-01
1e-03
1e-05
1
100
10000
Peaks (ordered)

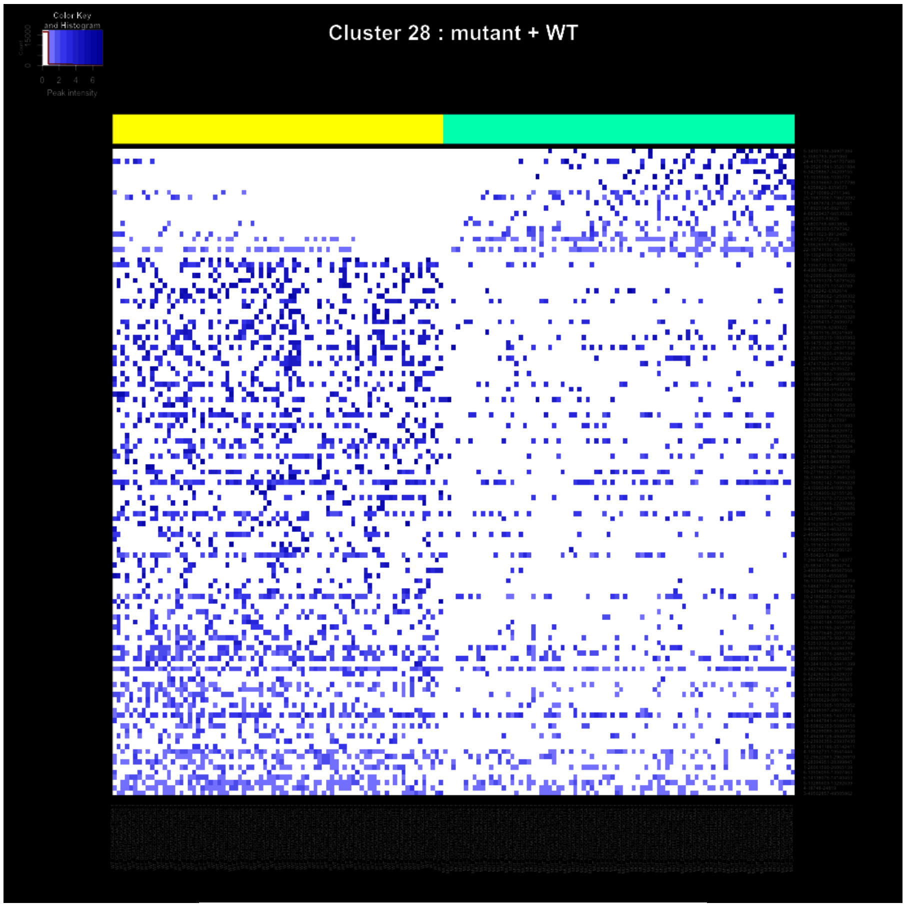
Color Key and Histogram
Peak intensity
0 2 4 6
Cluster 28 : mutant + WT

Cluster 28

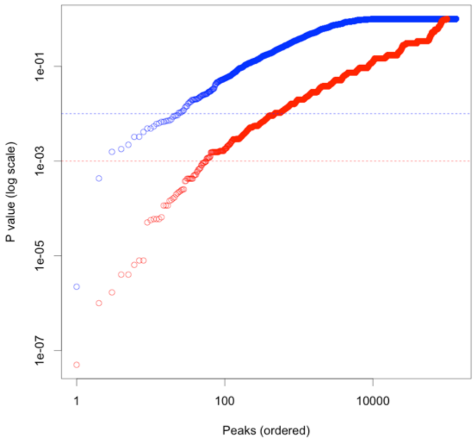
P value (log scale)
1e-01
1e-03
1e-05
1e-07
1
100
10000
Peaks (ordered)

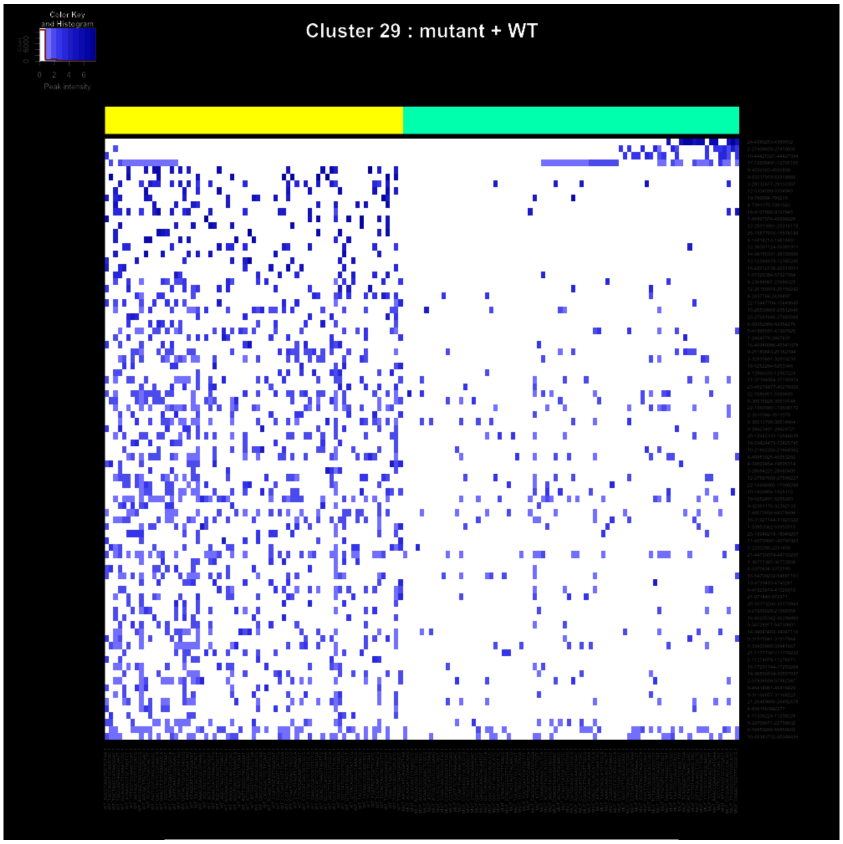
Color Key and Histogram
0 2 4 6
Peak intensity
Cluster 29 : mutant + WT

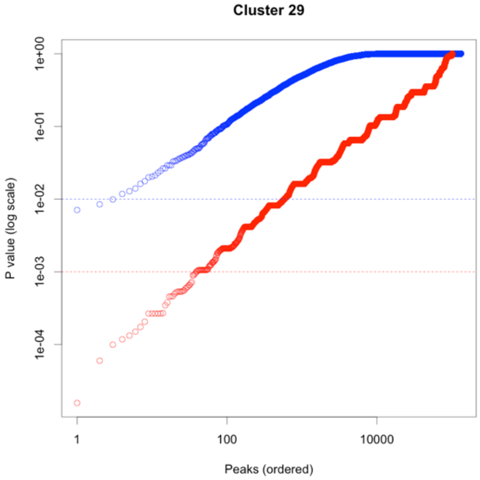
Cluster 29
P value (log scale)
1e+00
1e-01
1e-02
1e-03
1e-04
1
100
10000
Peaks (ordered)

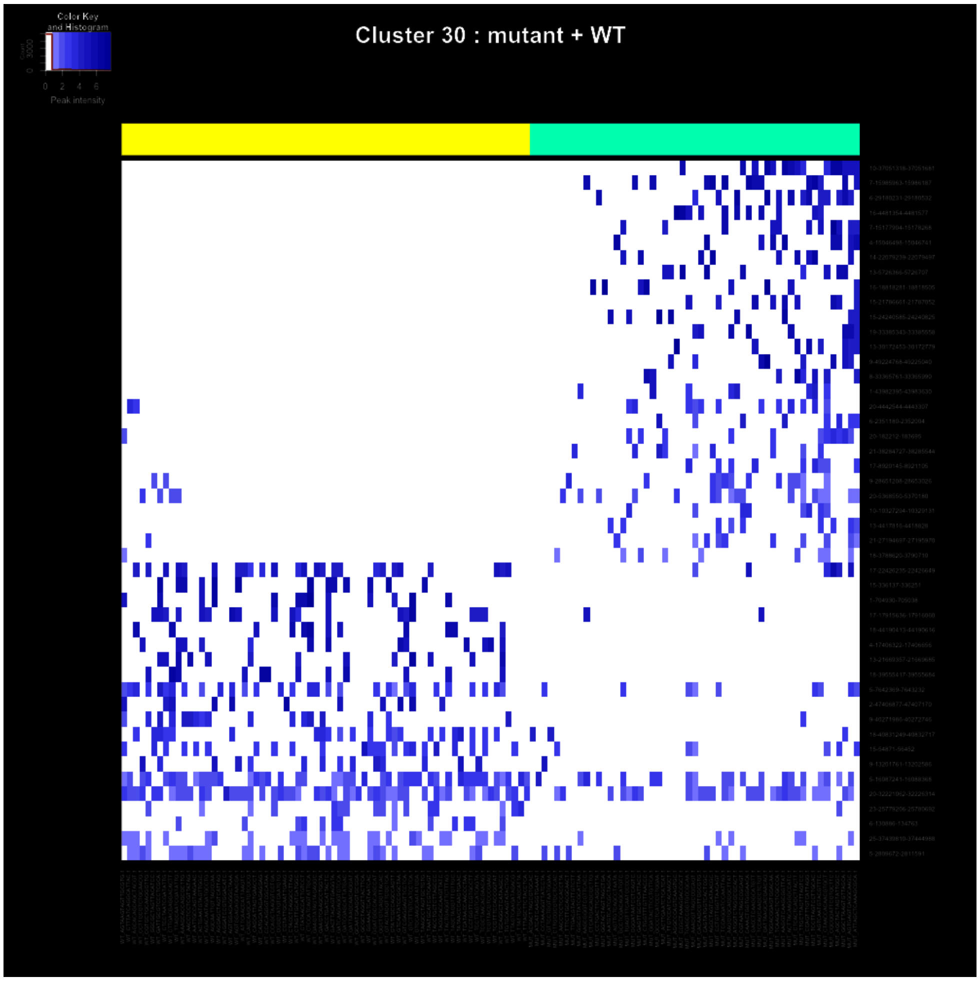
Cluster 30 : mutant + WT
Color Key and Histogram
Count
0
3000
0 2 4 6
Peak intensity

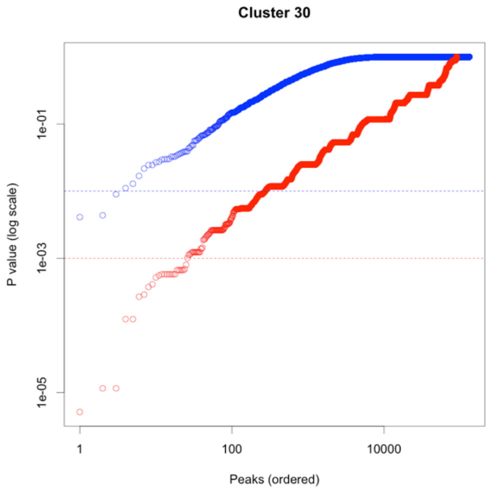
Cluster 30
P value (log scale)
1e-01
1e-03
1e-05
1
100
10000
Peaks (ordered)

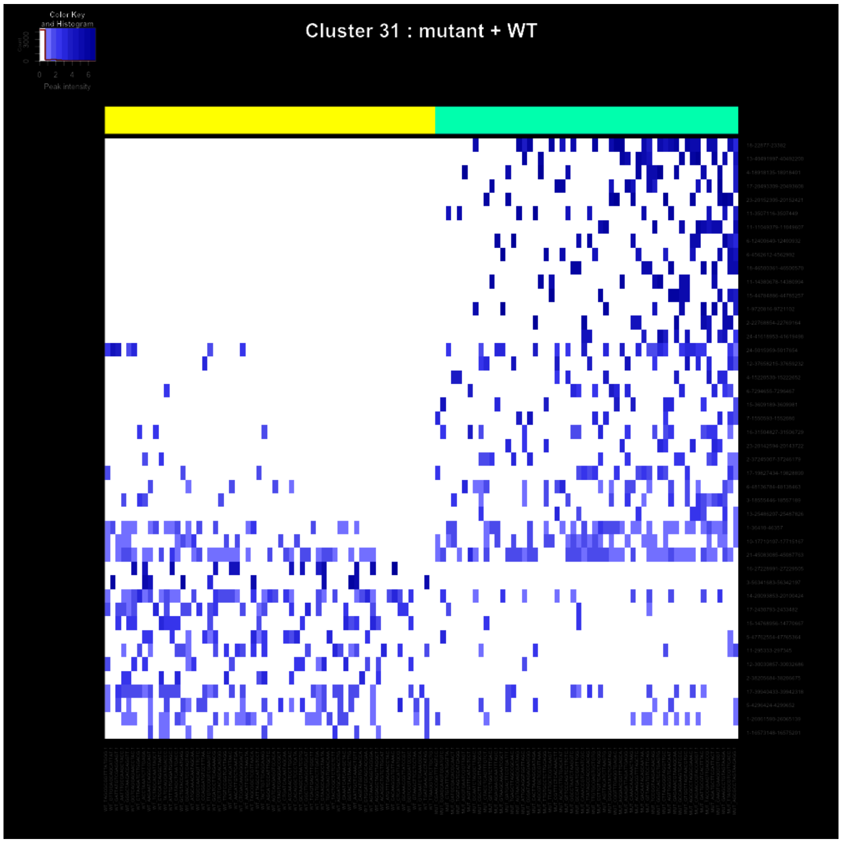
Cluster 31 : mutant + WT
Color Key
and Histogram
Peak intensity

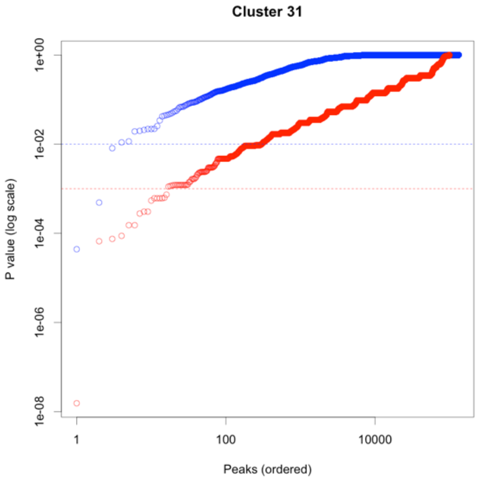
Cluster 31
P value (log scale)
1e+00
1e-02
1e-04
1e-06
1e-08
1
100
10000
Peaks (ordered)

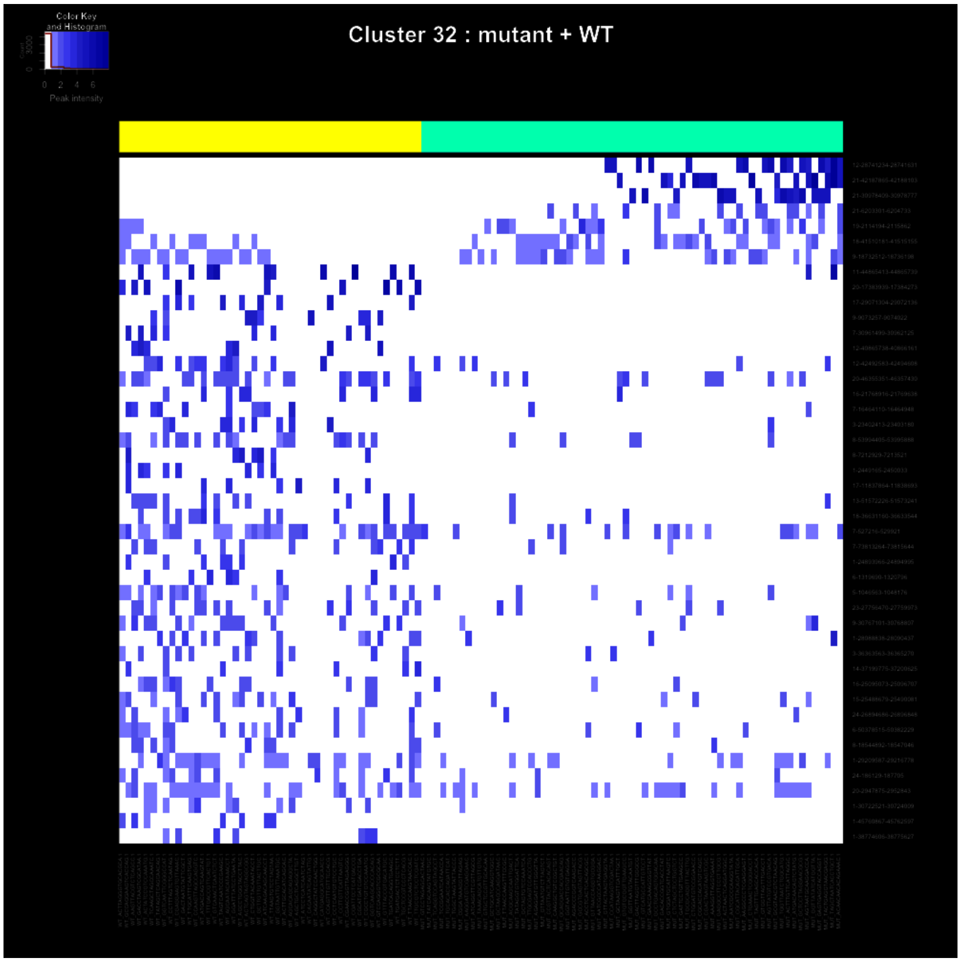

Color Key and Histogram
Peak intensity
Cluster 32 : mutant + WT


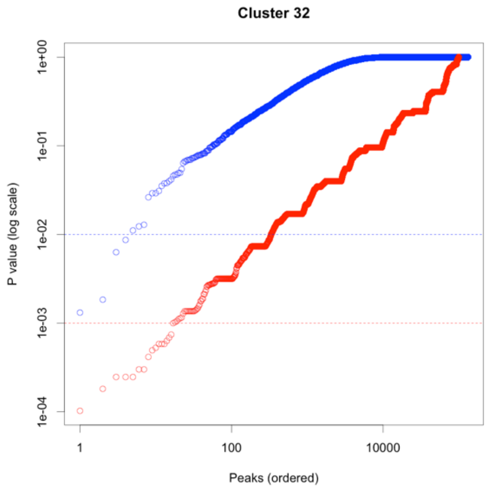

Cluster 32
P value (log scale)
1e+00
1e-01
1e-02
1e-03
1e-04
1
100
10000
Peaks (ordered)

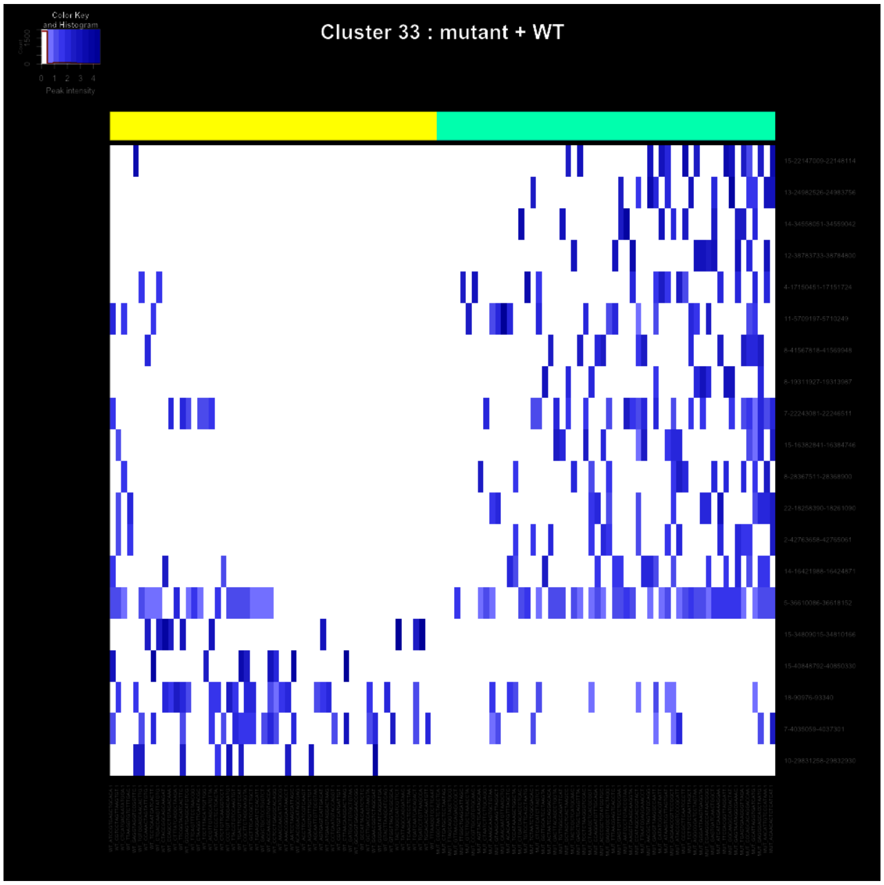
Cluster 33 : mutant + WT
Color Key and Histogram
Peak intensity

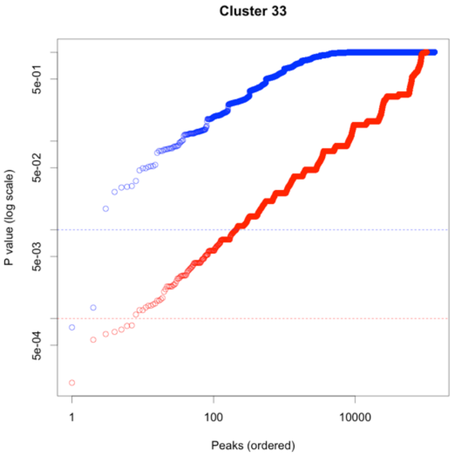
Cluster 33
P value (log scale)
5e-01
5e-02
5e-03
5e-04
1
100
10000
Peaks (ordered)

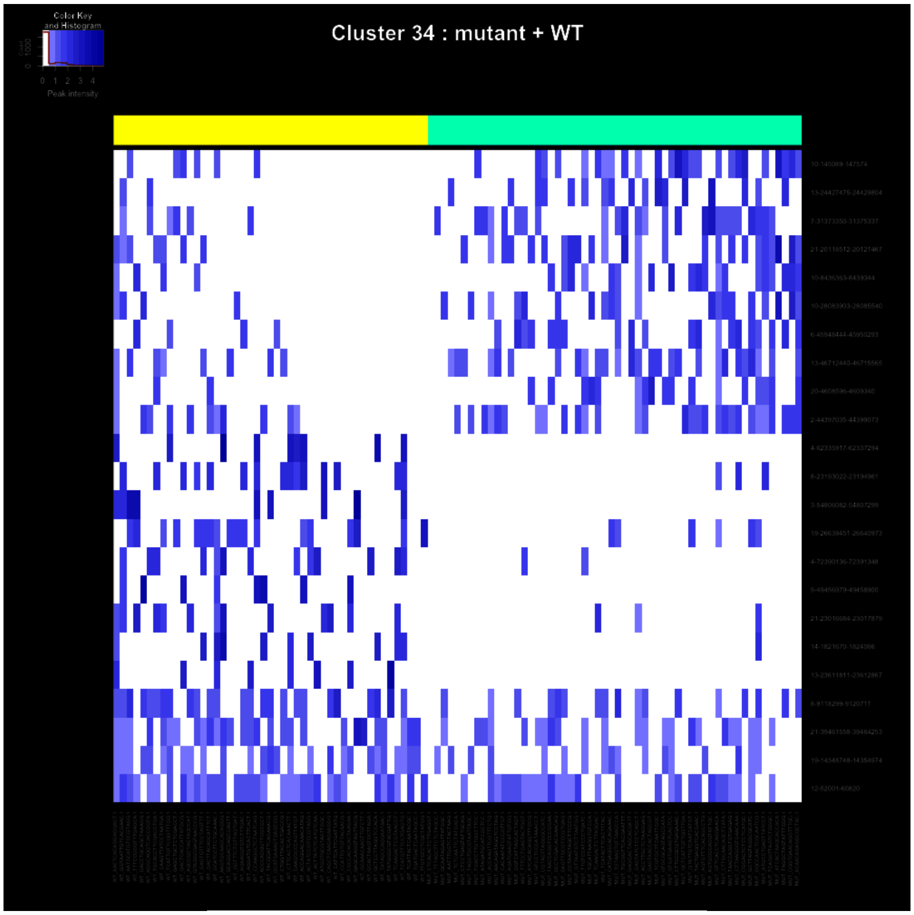
Color Key
and Histogram
Peak intensity
Cluster 34 : mutant + WT

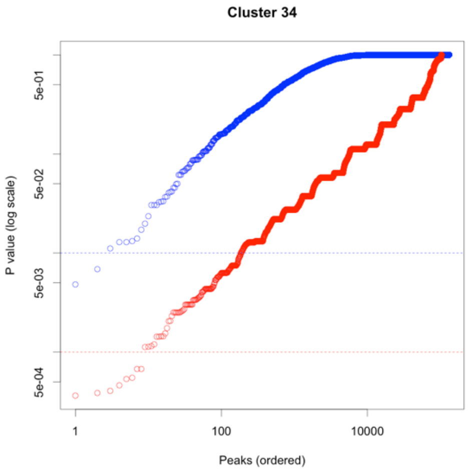
Cluster 34
P value (log scale)
5e-01
5e-02
5e-03
5e-04
1
100
10000
Peaks (ordered)

Cluster 35

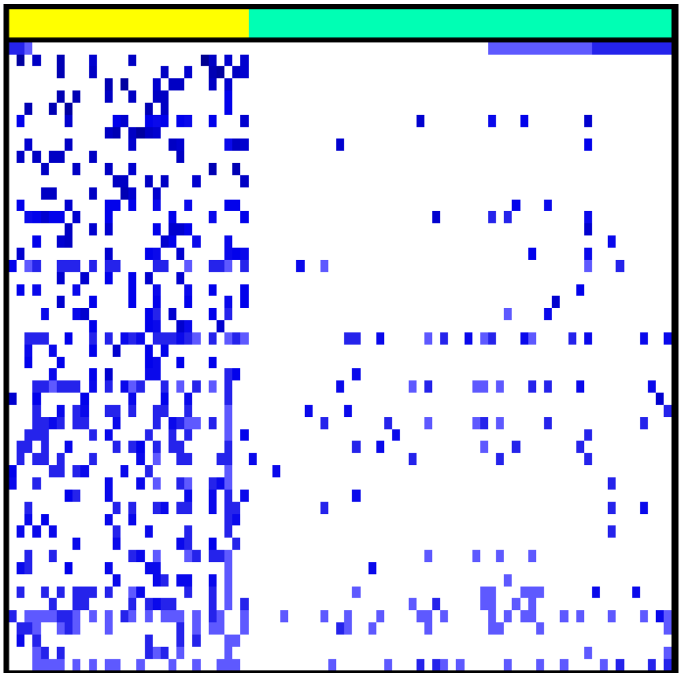

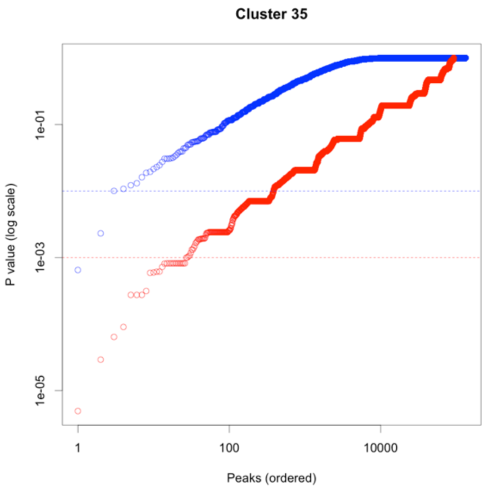

Cluster 35
P value (log scale)
1e-01
1e-03
1e-05
1
100
10000
Peaks (ordered)

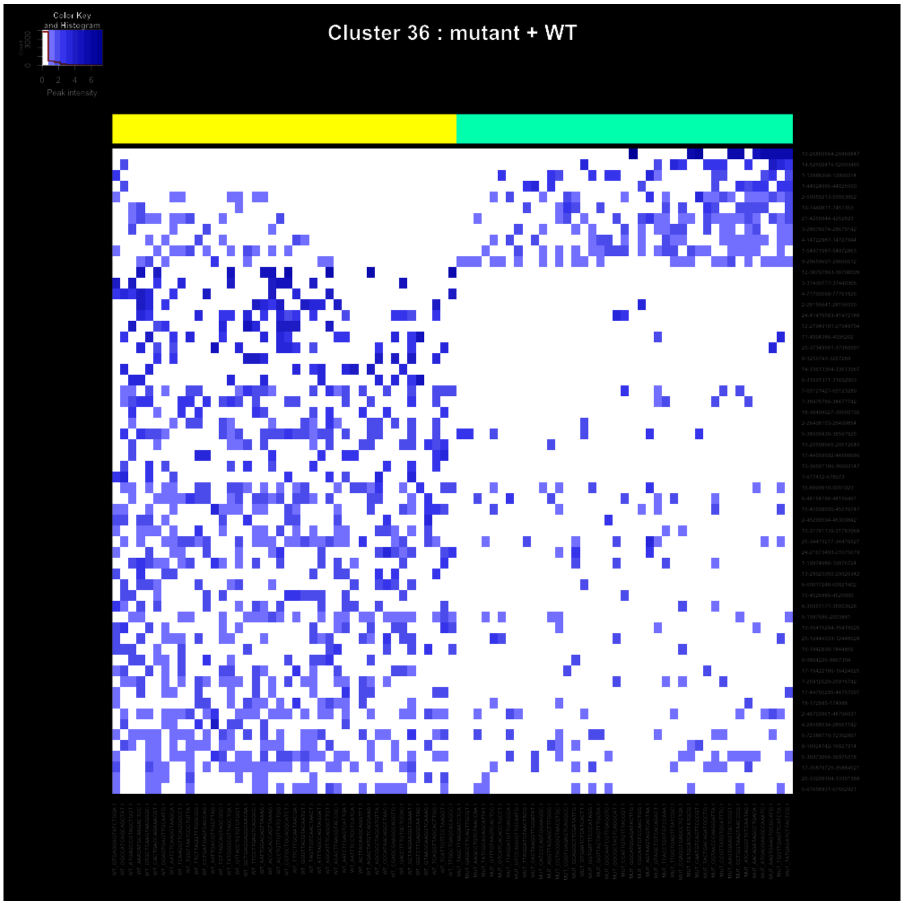
Color Key and Histogram
Peak intensity
Cluster 36 : mutant + WT

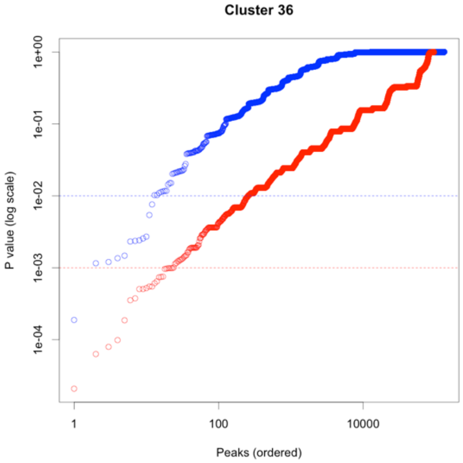
Cluster 36
P value (log scale)
1e+00
1e-01
1e-02
1e-03
1e-04
1
100
10000
Peaks (ordered)

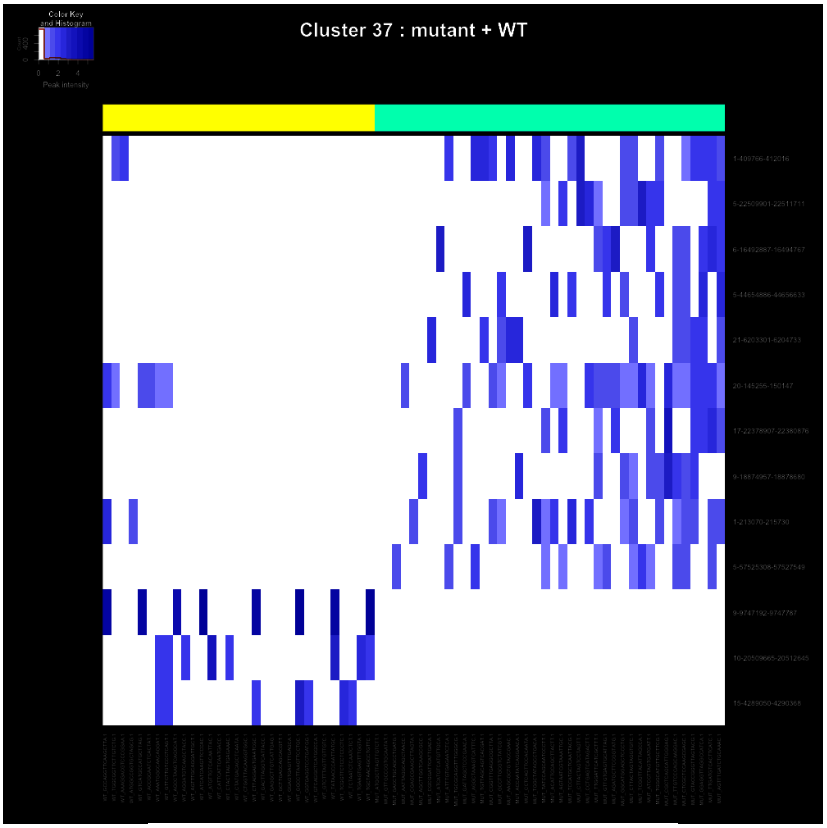
Cluster 37 : mutant + WT
Color Key and Histogram
Peak intensity

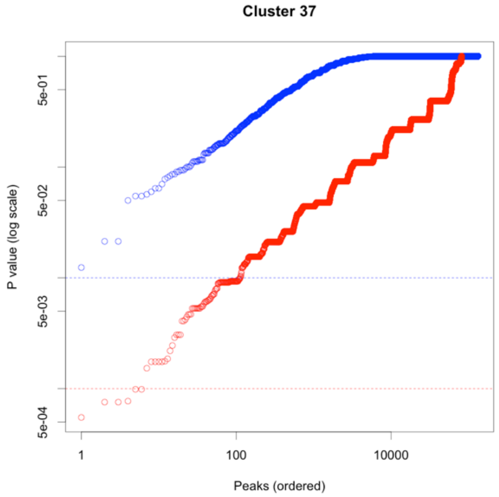
Cluster 37
P value (log scale)
5e-01
5e-02
5e-03
5e-04
1
100
10000
Peaks (ordered)

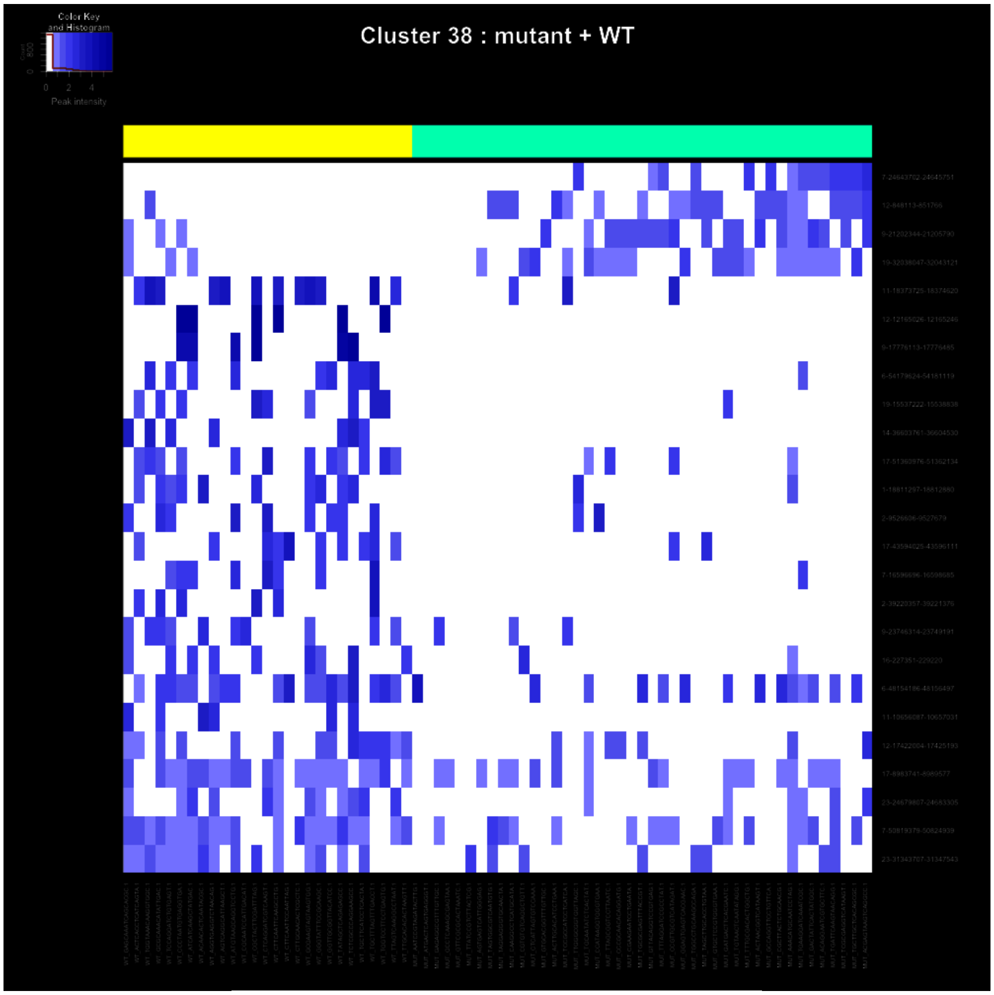
Color Key
and Histogram
0
2
4
Peak intensity
Cluster 38 : mutant + WT

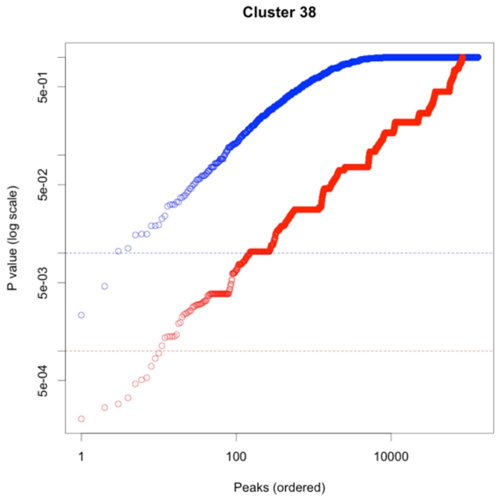
Cluster 38
P value (log scale)
5e-01
5e-02
5e-03
5e-04
1
100
10000
Peaks (ordered)

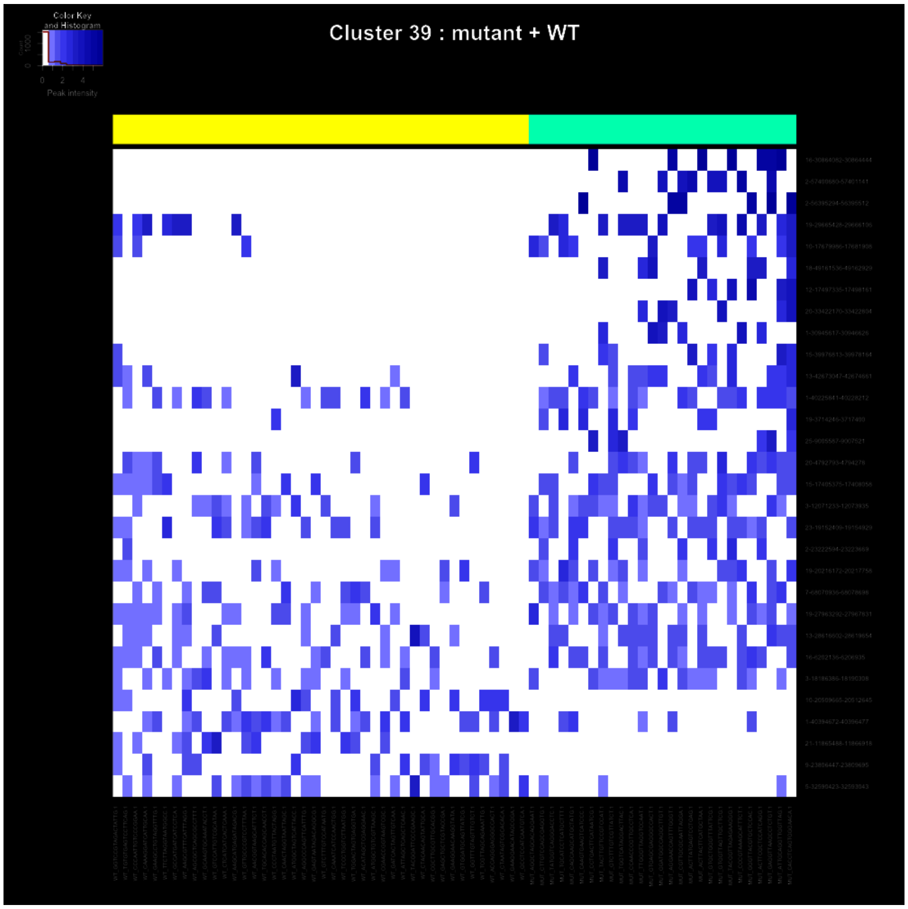
Cluster 39 : mutant + WT
Color Key and Histogram
0
2
4
Peak intensity

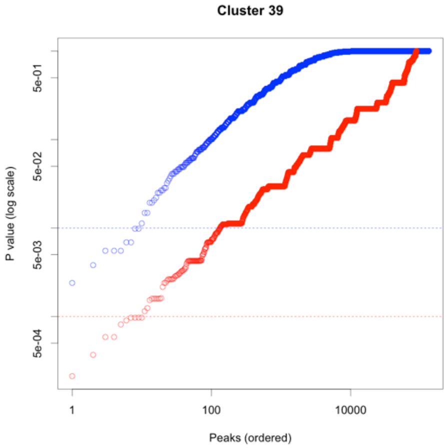
Cluster 39
5e-01
5e-02
5e-03
5e-04
P value (log scale)
1
100
10000
Peaks (ordered)

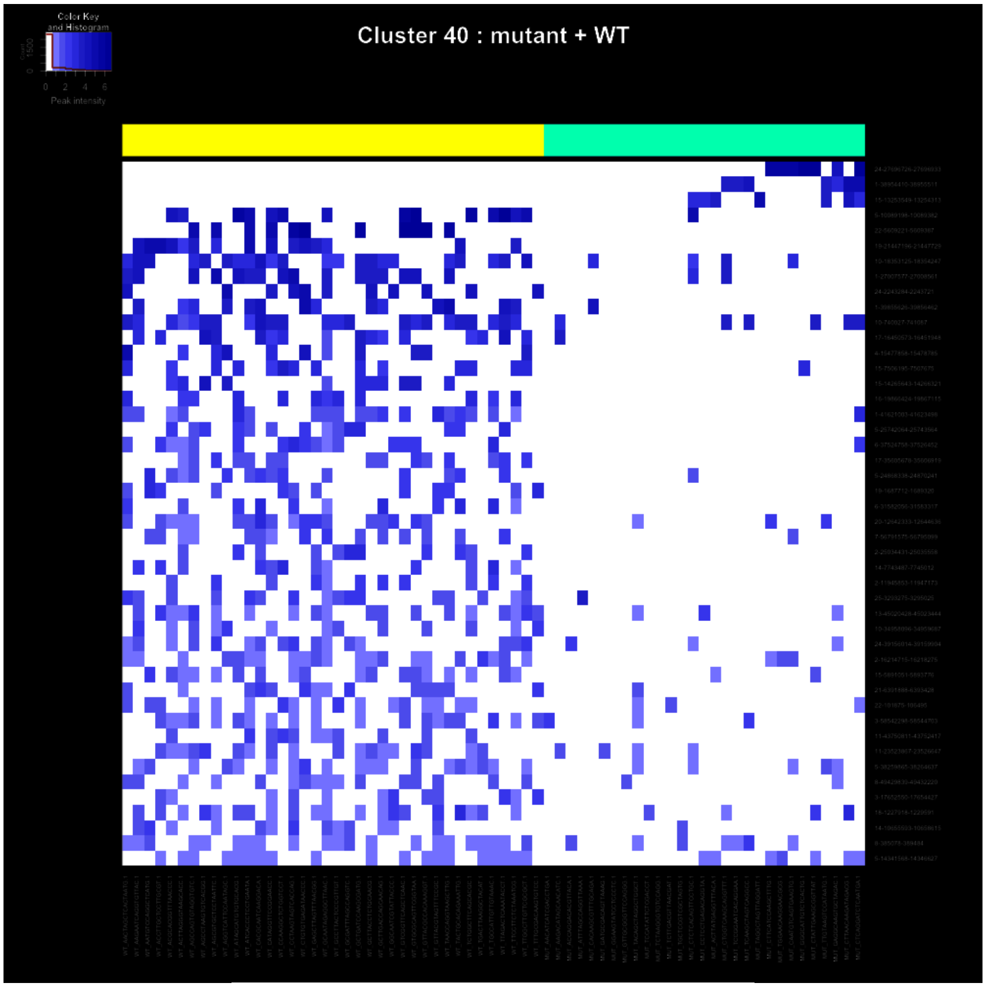
Color Key and Histogram
Peak intensity
Cluster 40 : mutant + WT

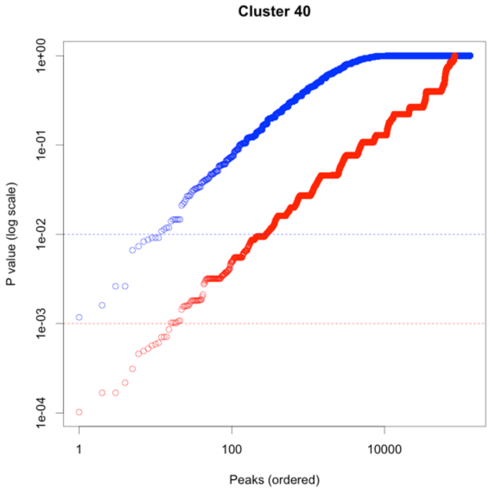
Cluster 40
P value (log scale)
1e+00
1e-01
1e-02
1e-03
1e-04
1
100
10000
Peaks (ordered)

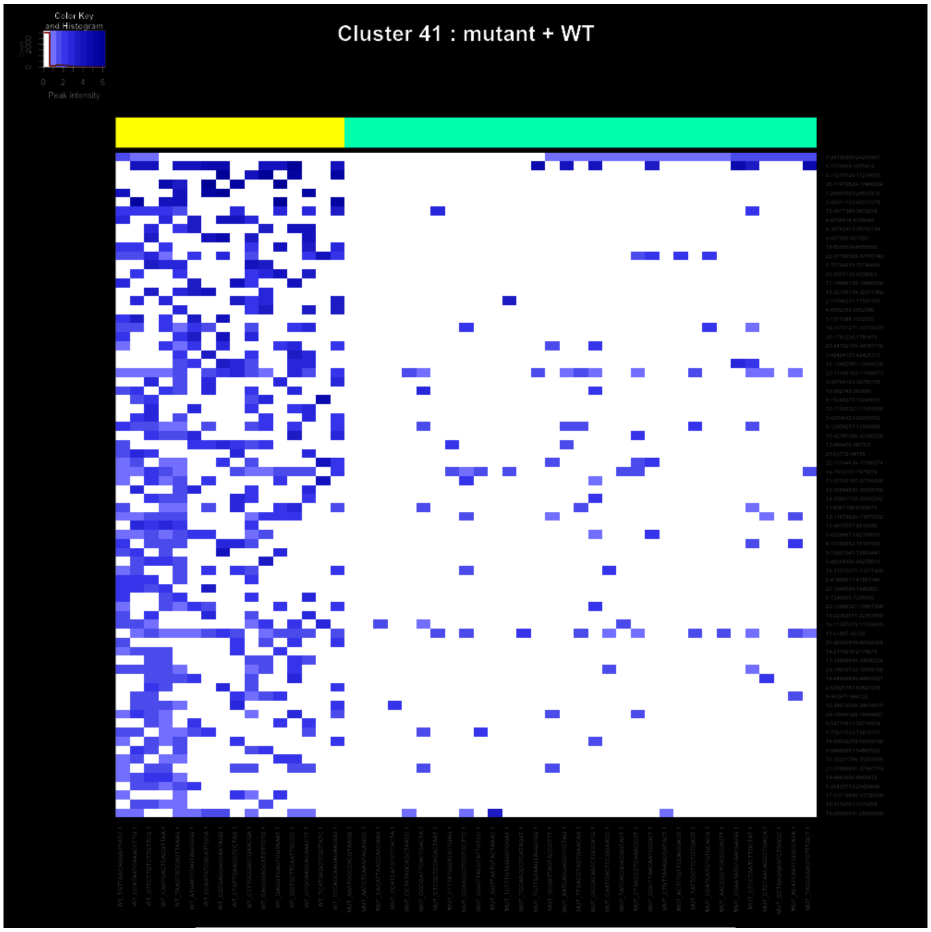
Cluster 41 : mutant + WT
Color Key
and Histogram
Peak intensity

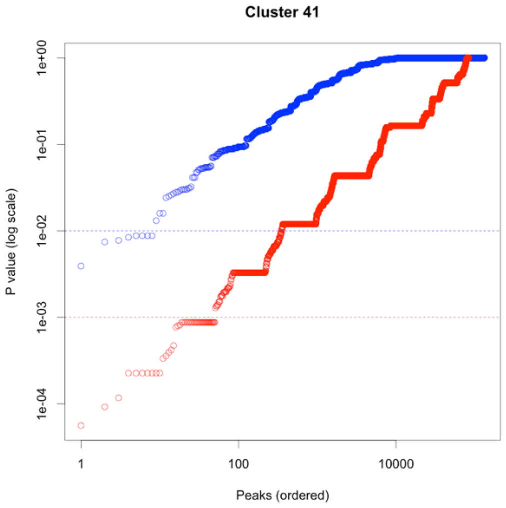
Cluster 41
P value (log scale)
1e+00
1e-01
1e-02
1e-03
1e-04
1
100
10000
Peaks (ordered)

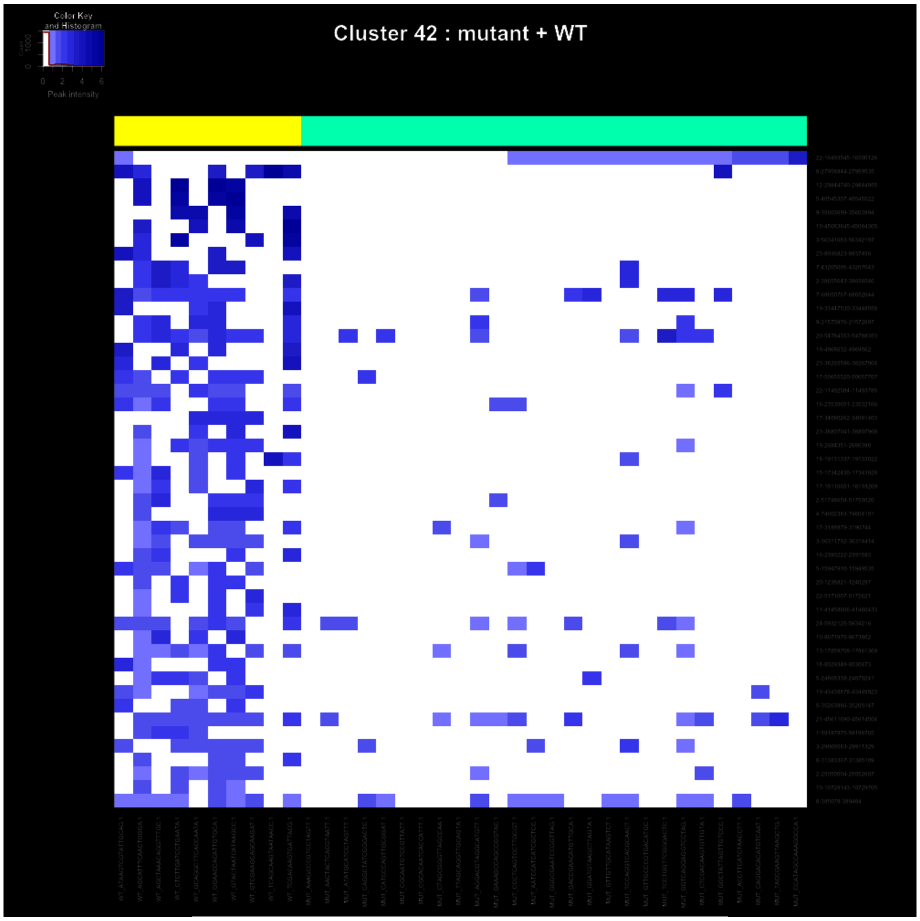
Color Key and Histogram
Peak intensity
0 2 4 6
Cluster 42 : mutant + WT

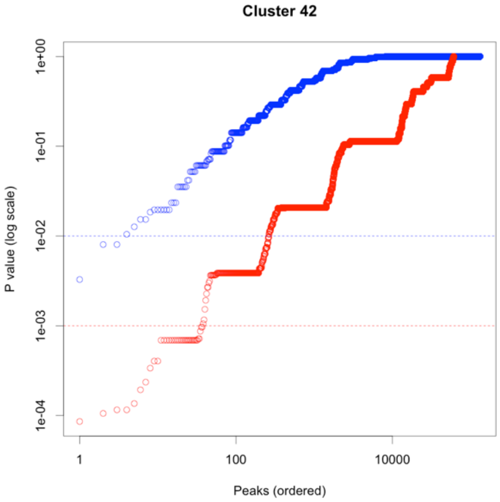
Cluster 42
P value (log scale)
1e+00
1e-01
1e-02
1e-03
1e-04
1
100
10000
Peaks (ordered)
